# Station and train surface microbiomes of Mexico City’s metro (subway/underground)

**DOI:** 10.1101/735027

**Authors:** Apolinar Misael Hernández, Daniela Vargas-Robles, Luis D. Alcaraz, Mariana Peimbert

## Abstract

The metro is one of the more representative urban systems of Mexico City, and it transports approximately 4.5 million commuters every day. Large crowds promote the constant exchange of human and environmental microbes. In this study, we determined the bacterial diversity profile of the Mexico City subway by massive sequencing of the 16S rRNA gene. We identified a total of 50,197 operative taxonomic units (OTUs) and 1058 genera. The metro microbiome was dominated by the phylum *Actinobacteria* and by the genera *Propionibacterium* (15%) (*P. acnes* 13%), *Corynebacterium* (13%), *Streptococcus* (9%), and *Staphylococcus* (5%) (*S. epidermidis;* 4%), reflecting the microbe composition of normal human skin. The metro microbial sources were skin, dust, saliva, and vaginal, with no fecal contribution detected. A total of 420 bacterial genera were universal to the twelve metro lines tested, and they contributed to 99.10% of the abundance. The large OTUs number are probably reflecting the vast human influx, while selection from hosts and environments are constraining the genera diversity, shown by the OTUs to genus ratio. Finally, this study shows that the microbial composition of the Mexico City subway comes from a mixture of environmental and human sources and that commuters are exposed to normal human microbiota.

## Introduction

Public transport systems provide the ideal environment for the transmission of microorganisms, as they carry a multitude of people, and their microbiomes, on a daily basis. The metro can assemble an extensive repository of beneficial bacteria, such as commensals and symbionts, or harmful bacteria, becoming a vehicle for the transmission of infectious diseases. The Mexico City metropolitan area has a population of 21.3 million people, which makes it the largest city in the Western Hemisphere. Mexico City’s metro has more than 4 million users every day, totaling 1,647,475,013 users annually (https://metro.cdmx.gob.mx/) and is the second busiest subway of the American continent, after New York city’s, and the ninth busiest in the world^1^. It has been in use since 1969 and today has 12 lines and 196 stations; it travels through a network of 226.5 km and operates 365 days of the year, covering a radius of around 10 km around the urban sprawl of Mexico City. The various lines were individually constructed at different points in history, which has led to differences in the infrastructure and the wagon type between lines; although all lines have pneumatic wheels, some of also have iron wheels. Most of the lines go through different sections, including underground, superficial, and elevated above ground level, and are ventilated by extractors, fans, or open windows. The size of passenger influx differs by line and by station, with higher influx terminal stations and transfer stations. Terminal stations connect the inner city to the suburban areas, where most of the population live. This mass movement of people breathe the same air and touch the same surfaces, promoting a large-scale interchange of human and environmental microbiota. Several factors differentiate the subways of each city. In Mexico City, there are three rush hours per day: the commute to and from work and another peak at noon that seems to correspond to the schools schedule. The number of trips/person/day is 2.37, similar to London and Seville (2.31 and 2.33, respectively), but much lower than the New York or Paris average (3.79 and 3.70, respectively)^2^.

Previous studies have indicated that a significant proportion of subway surface bacteria come from the skin of passengers, so the lifestyle of these passengers is also important. Among the chemical and physical factors directly influencing the microbiome, the most obvious are temperature and humidity, but there are many other important factors such as differences in the ventilation systems, whether or not the trains are pneumatic, and pollution. Therefore, studies from other networks can be used for reference, but the local diversity will only be understood by directly studying the subways of Mexico City. Multiple studies have characterized the subway microbiomes of various densely populated areas, such as New York City, Boston, Hong Kong, and Barcelona^3–8^. Mexico City is one of the most densely populated areas in the world (6,000 people/km^2^ in Mexico City; 9,800 people/km^2^ in the Greater Mexico City area). However, Mexico City’s subway microbiome has been scarcely explored. Only one previous study has characterized the colony-forming units (CFU; N = 175 isolates) in the air of the subway transport system, identifying three Fungi genera and concluding that the majority of bacteria were Grampositive bacilli^9^.

It has been estimated that humans are capable of emanating 10^6^ particles per hour^10–12^. As a rough estimation, the 4 x 10^6^ Mexico City metro users per day, with an average commuting time of 2.3 hours^13^, release a total of 9.2 x 10^12^ particles in the metro every day. Additionally, the metro architecture favors the presence of multiple aerosol particles from other sources, so the metro can be thought of as a vast shared pool for the horizontal exchange of human-derived, soil, plant, and microbes and their DNA. In previous work, the built environment has been shown to be biologically different when inhabited by people, compared to empty spaces, and even human “cloud” microbiomes can be used to distinguish the microbiota signatures of individuals four hours after they have departed^10,14^. In this study, we revealed the bacterial diversity profile of the Mexico City metro with massive sequencing of the 16S rRNA gene. We took 47 samples from station turnstiles and train handrails, covering all 12 metro lines of Mexico City. This work is the first culture-independent investigation of subway microbiome in Latin America.

## Results

We studied the twelve Mexico City metro lines and collected 47 samples for this study (**Fig. 1**). The samples were taken from station turnstiles (N = 24) and train handrails (N = 23). We sequenced a total of 16.6 x 10^6^ V3-V4 16S rRNA gene reads, spanning 4.98 Gb of coverage. The reads were merged into 5,788,162 sequences (avg = 460 bp); with a mean of 106,057 sequences/sample. This represents the largest published record of Mexico City’s metro microbiological diversity. The sequences were clustered into operational taxonomic units (OTUs; 97% identity 16S rRNA gene) recovering 50,174 OTUs. The OTUs were then homology-matched and summarized as 1,058 distinct bacteria genera (**Table S1**).

**Fig 1.**
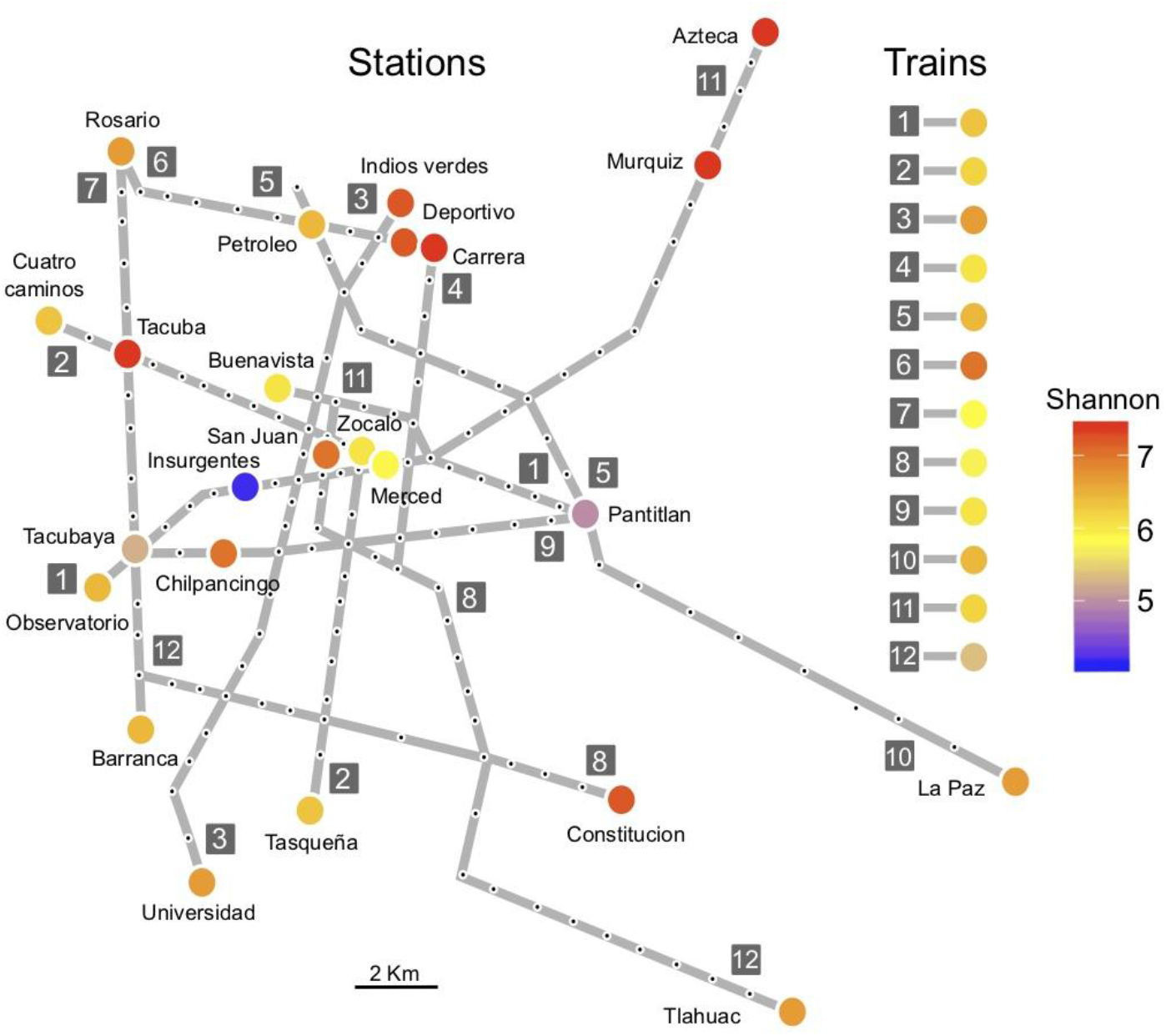
Sampling diversity and location. The metro network, with each black dot representing a station and colored circles showing the sampling locations. The circles are colored according to the Shannon diversity index. Train diversity is also shown in colored circles.

The most diverse station was Carrera, with a Shannon diversity index of H’ = 7.36, followed by Azteca station (H’ = 7.35), Muzquiz (H’ = 7.31), Tacuba station (H’ = 7.30), and Indios Verdes (H’ = 7.16); the lowest diversity was found on metro Line 1 (Insurgentes station; H’ = 4.10) (**Fig. 1**). The train with the highest diversity was found on Line 6 (H’ = 7.38), followed by Line 3 (H’ = 6.96), and line 5 (H’ = 6.63). The lowest diversity train was found on Line 8 (H’ = 5.54). The average Shannon diversity (H’_(average)_ = 6.51) from the stations (turnstiles) was higher than the average train (handrails) diversity (H’_(average)_ = 6.24) (**Fig. 2; Suppl. Table S2**). We performed a comprehensive sampling of the metro’s diversity, as we did not find differences between the observed and expected (Chao1) richness (**Fig. 2**) when comparing genera, and the observed and expected were in the same magnitude order when comparing OTUs. The Simpson diversity index showed a slightly increased dominance for the turnstile when compared to handrails (**Suppl. Table S2**).

**Fig. 2.**
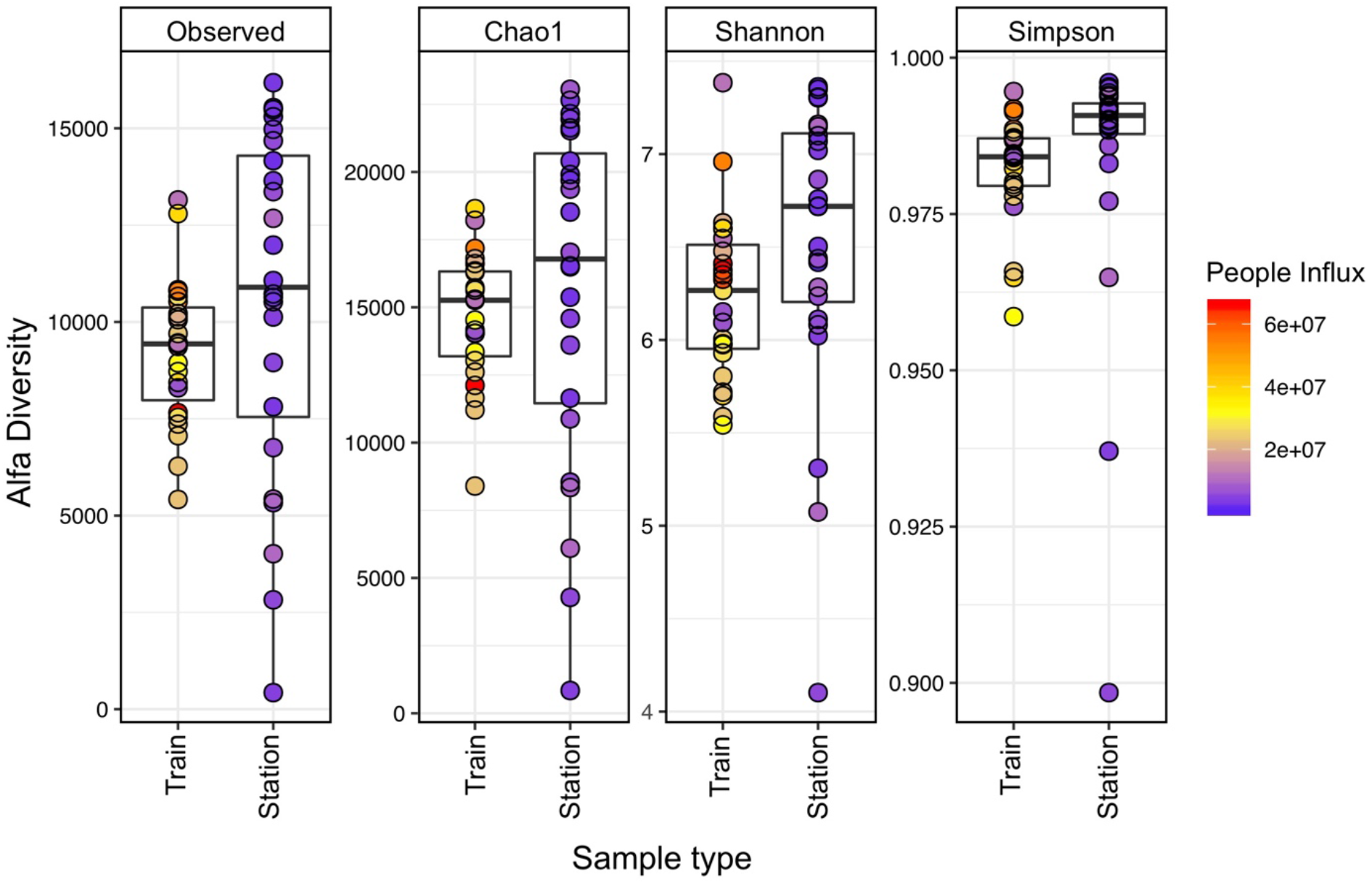
Richness and diversity of Mexico City’s metro. The surface microbiome diversity on the station entrance/exit turnstiles and the metro wagon handrails. A total of 50,174 OTUs were observed, across 47 samples.

We tested multiple factors regarding the microbial richness, diversity, and composition, looking for significant relationships among metro line numbers, station connectivity (regular, transfer, and terminal stations), station surroundings (bus terminals, work-related buildings, schools, hospitals), architectural structure (above ground, below ground stations), train wheels (metallic and pneumatics), train design (individual wagons or connected), and physical variables like temperature, relative humidity, geographical zones, and people influx. However, there were only statistically significant differences when we compared the surfaces of station turnstiles with train handrails (**Fig. S1**). Turnstiles showed higher OTU diversity than handrails (OTU level, Simpson, KW p = 0.004) and had higher diversity dispersions (Observed OTUs and Chao1, p < 0.001, Fligner-Killeen test, **Fig. 2**). Similar results were observed at the genus level (for all metrics, p < 0.003, KW, **Fig. S2**), which had similar dispersions (p > 0.094, Fligner-Killeen test, **Fig. S2)**.

Mexico City’s metro is highly diverse at the OTU level (50,174 OTUs). By merging the OTU diversity with the bacterial genera diversity by line, we found that 420/1,058 genera were ubiquitous throughout the metro system (**Fig. 3**). Moreover, these 420 genera, representing 3,688 OTUs, constituted 99.10% of the whole dataset, expressed in sequence abundance. The other 638/1,058 genera contributed to just 0.9% of the overall richness. We are confident this work has comprehensively described the Mexico City metro bacterial genera diversity, and that the diversity has a fairly homogeneous distribution across the metro system. We concluded the main phyla detected were *Actinobacteria, Firmicutes, Proteobacteria, Cyanobacteria, Bacteroidetes, Chloroflexi, Fusobacteria, a*nd *Thermi* (**Fig. 4A**).

**Fig. 3.**
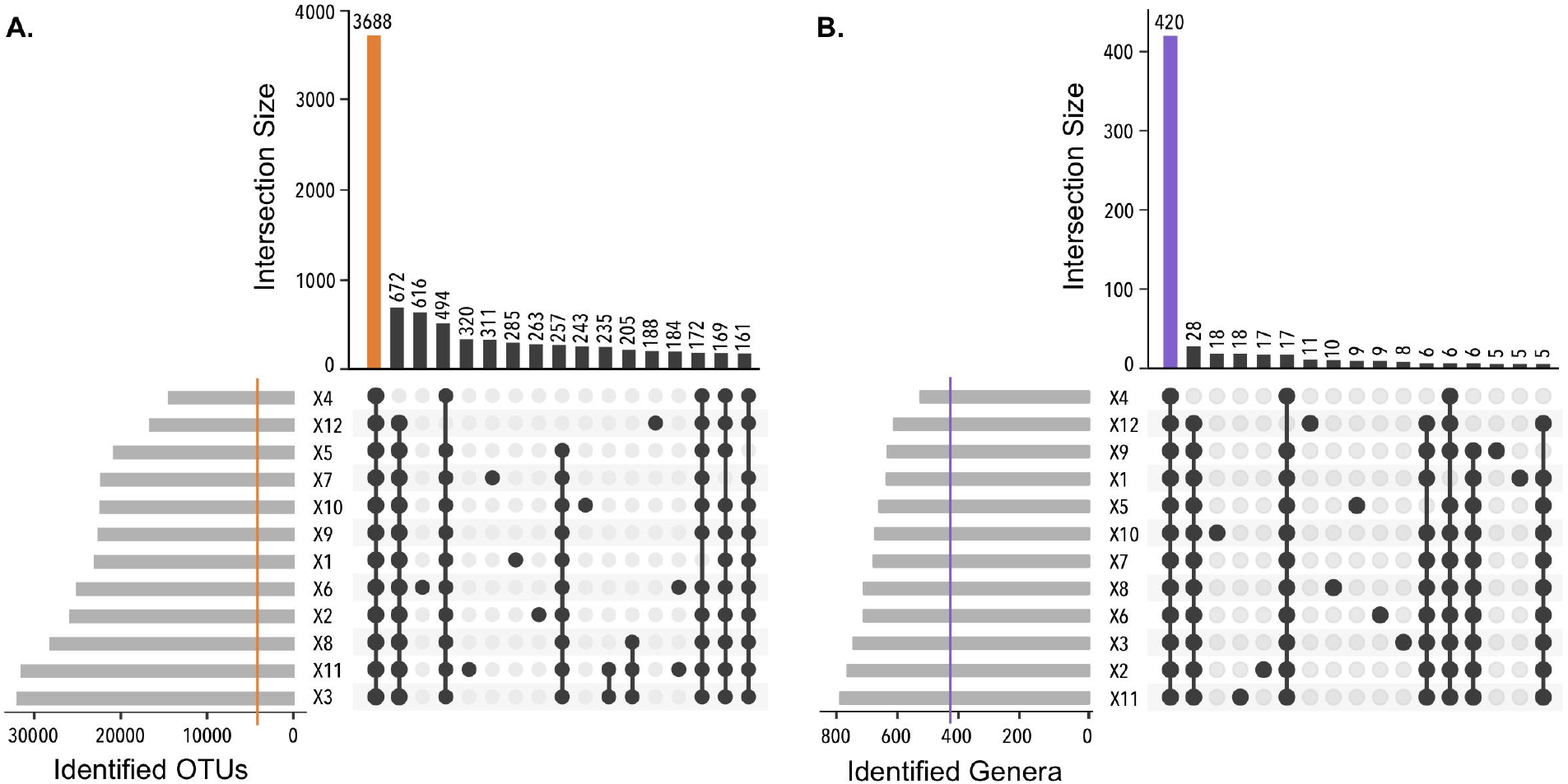
Shared taxa between subway lines, train, and station samples. **A.** Shared OTUs: of the 50,174 OTUs, 3,688 were found in all lines. **B.** Shared genera: of 1,058 genera, 420 were found in all lines. The 420 genera represented 99.10% of the entire dataset. The colored line in cardinalities of sets indicates the number of shared elements. Sets smaller than 150 elements for OTUs, and sets smaller than 5 elements for genera, were excluded from the diagrams.

**Fig. 4.**
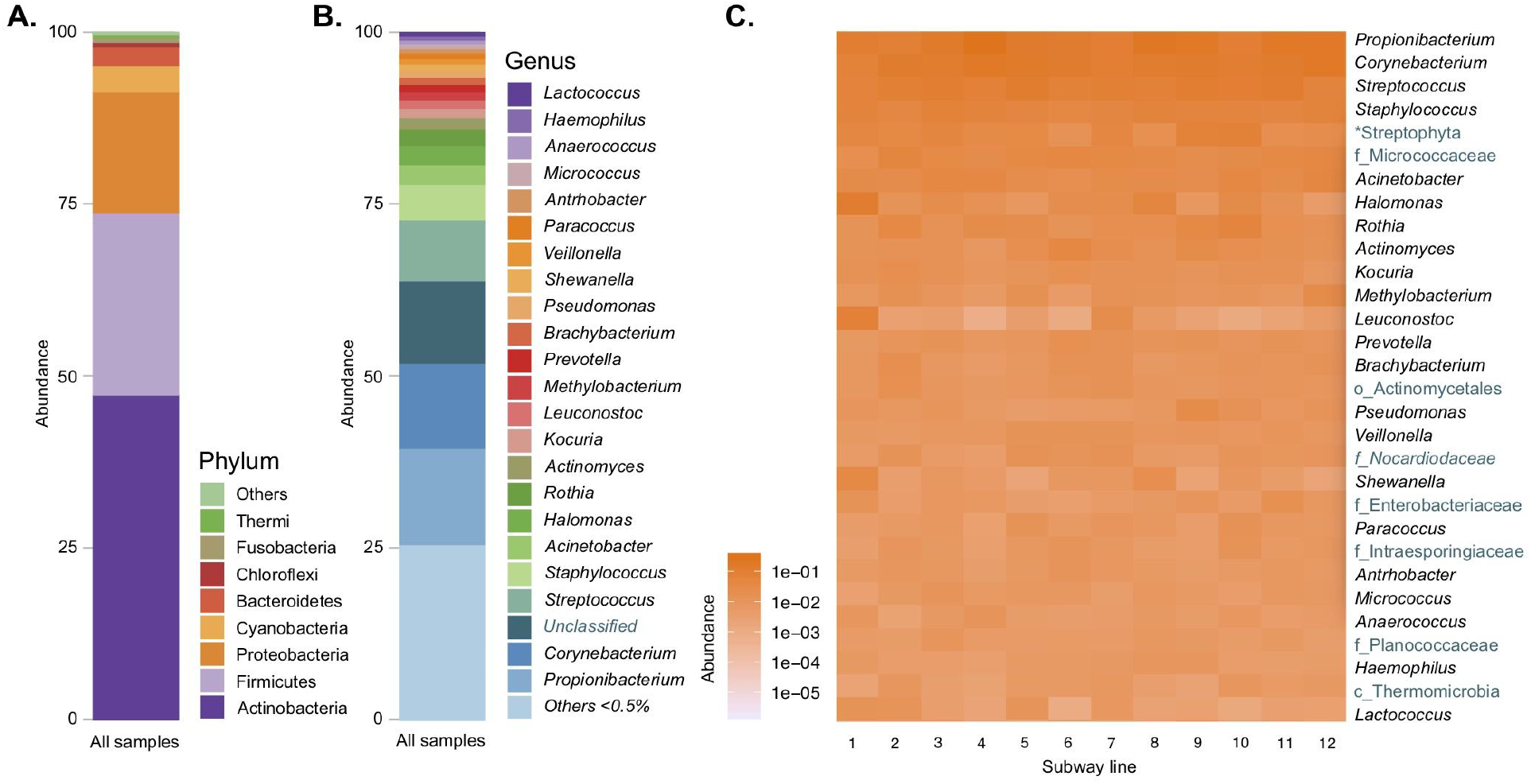
Phylogenetic profile of Mexico City’s metro surfaces. **A.** Phylum abundance in all samples: *Actinobacteria* is the most abundant phylum followed by *Firmicutes* and *Proteobacteria*. **B.** Genus abundance: *Propionibacterium* was the most abundant genus. **C.** Heatmap of the 30 most abundant genera and their abundance in each sampled metro line. The unknown genera were classified to the upper known taxonomic rank, shown in different text color.

There are multiple genera with low abundance (1,034 genera < 0.5%) in our study, and the most abundant known genera were *Propionibacterium* (15%), *Corynebacterium* (13%), *Streptococcus* (9%), and *Staphylococcus* (5%) (**Fig. 4B**). Interestingly, some of the most abundant genera were unclassified, but we were able to assign them to upper taxonomic hierarchies (29.43%). The ubiquity and prevalence of the core genera are exemplified in a heatmap showing the 30 most abundant genera in the system, which are also dominant in each metro line (**Fig. 4C**); the complete 420 genera heatmap and table are also available (**Suppl. Fig. S3, Suppl. Table S3**). The heatmap also shows the upper taxonomic level classifications for unknown genera and points out some candidates for examination, such as members of the families *Micrococcaceae*, *Nocardiaceae, Enterobacteriaceae, Intraesporingiaceae*, and *Planococcaceae;* along with members from the order *Actinomycetales* and the class *Thermomicrobia* (**Fig. 4C**). The fifth most abundant phylotype was classified as chloroplast (*Streptophyta*). Archaea were also marginally detected with a relative abundance of 0.001%, and the most represented genus was *Methanobrevibacter* (0.0004%).

The metro environment seems to be quite constant; however, there are subtle differences at the OTU level that suggest the possibility of biomarkers to discriminate between the environments. The higher diversity and OTU richness of station turnstiles, compared to wagon handrails, seemed to split our study into two distinct groups when we performed *β*-diversity analysis. Constrained analysis of principal coordinates (CAP) ordinations based on unweighted (**Fig. 5)** and weighted (**Fig S4)** UniFrac distances at the OTU level clearly segregated trains from stations (unweighted and weighted, p ≤ 0.001, PERMANOVA). Sample dispersion remained similar between groups (unweighted, p = 0.182, weighted, p = 0.242, BETADISPER). Similar results were observed at the genus level using Bray Curtis dissimilarities. Although, a total of only 7.5% variance was explained in both CAP axes (**Fig. 5a**). The CAP-tested variables showed that exposure to open environments and the turnstile samples were associated with CAP1 dispersion, while humidity and temperature were the best options for explaining CAP2 dispersion (**Fig. 5a**). Seven train communities were clustered closer to stations when we analyzed the same data as a clustering dendrogram, plotting the unweighted Unifrac distances, and two clades were exclusive to handrails (**Fig. 5b**).

**Fig 5.**
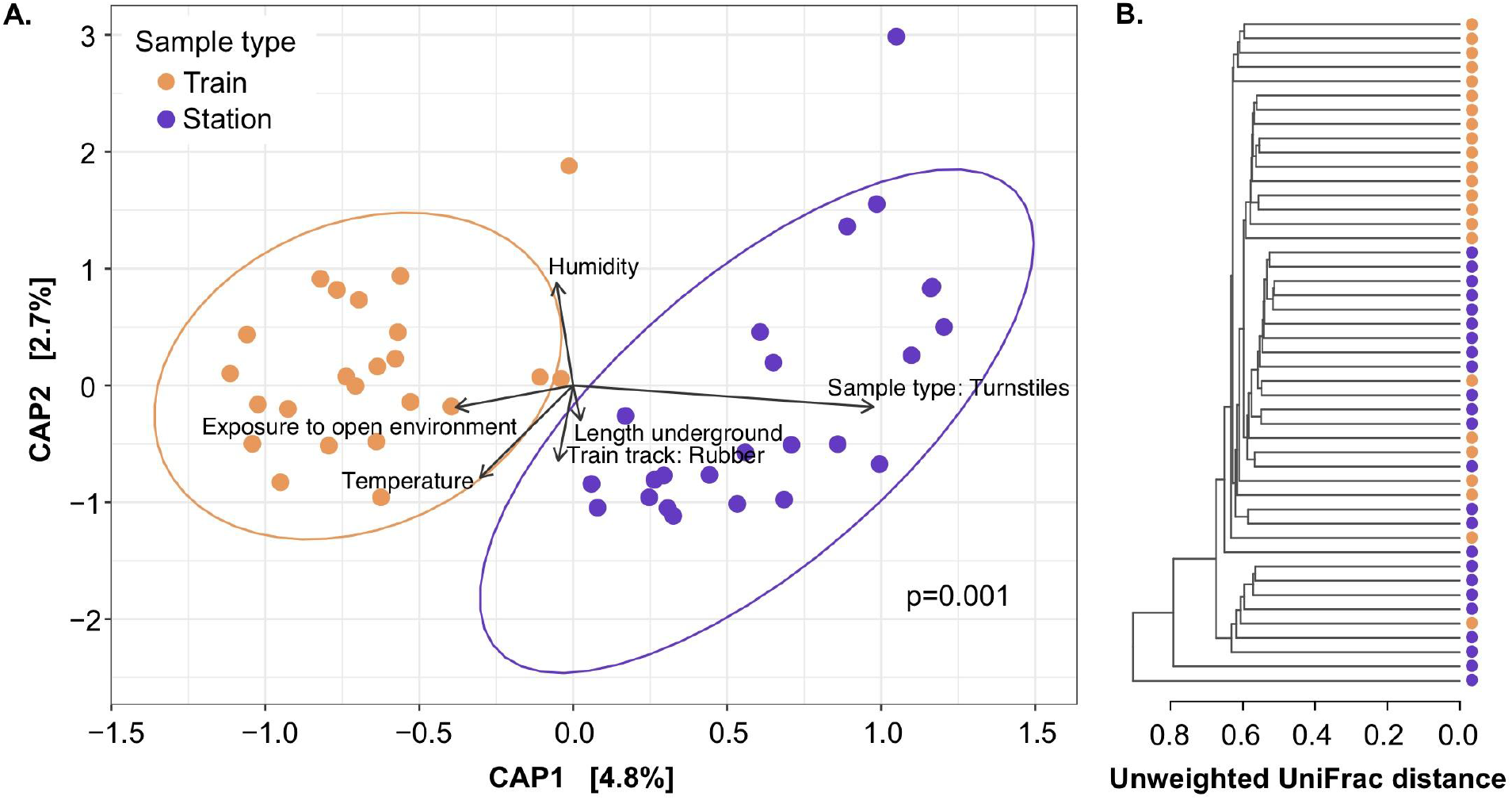
Metro bacterial communities grouped into the sample types collected in this study: for train and station. **A.** Constrained analysis of principal coordinates (CAP) ordinations based on unweighted UniFrac distances for turnstiles and handrails at the OTU level, significantly segregated (p = 0.001, Adonis) ellipses denote the 95% confidence interval of the points distribution by sample type. **B.** Community distances dendrogram using unweighted UniFrac distances.

In the taxa discriminant analysis, turnstiles were inhabited by multiple taxa, mostly from Actinobacteria phylum. The most enriched turnstile OTU was classified at the family level as *Micrococcaceae* (OTU 411738), and the most common significantly enriched genus was *Kocuria*, followed by *Blastococcus* and *Arthrobacter*, along with 34 other known genera. The handrail microbiomes were inhabited by the multi-environment distributed *Rothia, Streptomyces*, and *Veillonella* and the human bacteria *Propionibacterium* and *Gardnerella* (*G. vaginalis*) **(Fig S5)**.

We performed a source-tracking analysis to determine the possible environmental sources of the metro microbial communities. The tracked microbial environments were dust, skin, saliva, vaginal, stool, and soil. The primary environmental source of our data was determined to be dust (mean of 34% of the contribution), skin (32%), saliva (13%), soil (4%), and only 14 samples from the vagina (0.1%). No stool profiles were detected. Train handrails showed a higher level of skin microbiota than turnstiles (p = 1e-06, Student’s t-test), while station turnstiles had a higher level of dust and soil than handrails (p = 5e-4 and p = 3e-05, respectively, Student’s t-test). Saliva and vaginal contributions were not distributed differentially on handrails than turnstiles. We did not detect significant differences in environmental sources between samples exposed to the external environment or completely subterraneous. In multiple samples, the primary environmental source remains unknown (**Fig. 6y and Fig S6**).

**Fig. 6.**
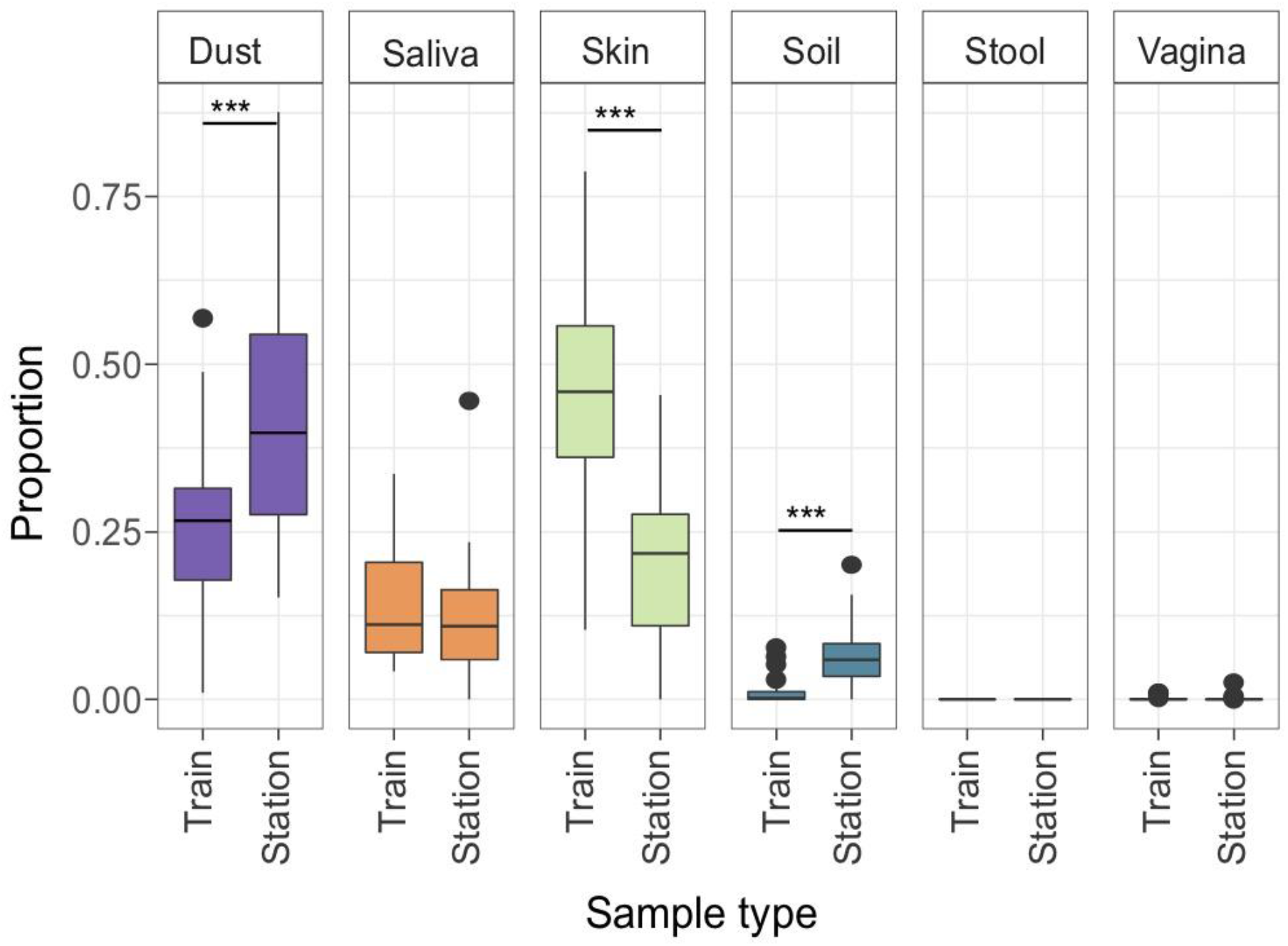
Source tracking comparison between turnstile and handrail microbiota. A source tracker algorithm was used to identify microbes to the genus level, using dust, soil, saliva, skin, feces, and vaginal samples as potential sources (***p < 0.001, Student’s t-test).

## Discussion

The microbiome diversity richness of the Mexico City metro was 50,174 OTUs (16S rRNA gene V3-V4), which is comparable to reports from other cities. For example, in a study of the Hong Kong subway system, using quite similar methodological procedures (16S rRNA gene V4), there were reported to be a total of 55,703 OTUs^6^. However, Shannon’s diversity index (H’) is two magnitude orders larger in Mexico City’s metro (H’_(average)_ = 6.38 ± 0.661) than the Hong Kong subway (H’_(average)_ = 4.13 ± 0.307^15^) or the Barcelona subway airborne bacteria (H’ ~ 1.5^8^). Additionally, the Simpson index of diversity (1-D_(average)_ = 0.98 ± 0.004) is also higher than the Boston surface microbial diversity (1-D ~ 0.75, as estimated from a previous study^4^). The Shannon diversity is higher than some human microbiome systems^16^. Overall bacterial diversity found in Mexico City’s metro is similar to subways in other countries, being overrepresented by human skin and oral taxa^3,5–8^. However, each city metro has its dominant taxa. *Pseudomonas stutzeri*, *Stenotrophomonas maltophila, Enterobacter cloacae*, and *Acinetobacter* are the most abundant species in New York City. *Methylobacterium is the dominant genus in Barcelona subway*. Boston and Hong Kong subways have similar bacterial composition to Mexico City’s metro with a dominance of *Propionibacterium*, *Corynebacterium, Streptococcus*, and *Staphylococcus*^4,6,15^. Neither *Pseudomonas* nor *Methylobacterium*, prevalent in NYC and Barcelona subways, are among the most abundant genera found in this study (**Fig. 4**). The metagenomic study of subways and urban biomes (MetaSUB) is a consortium working to develop sample collection, DNA/RNA isolation, and sequencing standard operating procedures to study global microbial diversity in mass transport systems^17^. This work began before the MetaSUB consortium, but the sampling, DNA isolation, PCR programs, sequencing technologies, and data processing are in line with the MetaSUB requirements.

In Mexico’s subway microbiome, the four most abundant genera were *Propionibacterium* spp. (15%) (*P. acnes* 13%), *Corynebacterium* spp. (13%), *Streptococcus* spp. (9%), and *Staphylococcus* spp. (5%) (*S. epidermidis*, 4%). We detected a large proportion of dust bacteria (compared against taxa profiles of mattress dust, which was mostly human skin debris). *P. acnes* on handrails reached a similar relative abundance to that reported for passengers’ hands after a subway trip (~29%^15^). Despite the similarities with Boston and Hong Kong bacterial diversity, our study has a larger OTU diversity. It is possible that Mexico City metro’s high OTU diversity and heterogeneity is related to the millions of users, while the homogeneous distribution at the genus level is the outcome of host-microbe interactions and their mutual selection^10,14^.

Multiple factors affect the subway microbiome composition, such as the number of users, microenvironmental and climatic conditions, architecture, and even rush hours, and some studies have identified antibiotic resistance genes as morning or afternoon signatures^15^. In this work, no architectonic designs (subterranean, street level, or elevated trains, or rubber or ferrous wheels) affected microbial diversity or composition. The high frequency of surface interaction by users probably promotes the rapid exchange of microbes, preventing them from establishing and growing. Environmental variables (temperature and humidity) appear to be similar among sampled sites; additionally, the type of environment outside the train station (work-related or bus station environment) did not modify the microbial diversity, suggesting the samples were independent of the station context.

The higher diversity we found on the turnstiles compared to handrails may be explained in a number of ways. The station turnstiles are exposed to the outside, which may explain the greater diversity and also why soil bacteria are more highly represented in these samples. Another relevant dissimilarity is the number of people who touch the bars: we observed that the rate of direct contact with one of the turnstiles bars is significantly higher (median, 122 people/hour, N = 8) than with one of the train vertical bars (median 22.9 people/hour, p = 0.001, Kruskal Wallis, N = 9, **Fig. S7**). Differences in usage may also explain the microbial diversity and composition^4^. While handrails are touched almost exclusively with the hands, turnstiles are also touched by clothes, which favors the arrival of dust-associated taxa. Additionally, the material differs between the surfaces, i.e., turnstiles are aluminum, while handrails are made of stainless steel.

The *Actinobacteria* phylum is highly represented in Mexico City’s metro, and we were able to identify 160 known genera and 3,959 undescribed genus-level OTUs. The leading players were found to be *Propionibacterium* and *Corynebacterium*. Interestingly *Propionibacterium* has been described in multiple environments, including soil, freshwater, marine, and host-associated, and while *Corynebacterium* has been described in these same environments, it has been more frequently found in marine environments^18^. *Propionibacterium* is divided into cutaneous and environmental strains, and some consider the cutaneous strain to be a primary human pathogen; it also has multiple biotechnological uses as a source of valuable metabolites like vitamin B12, trehalose, propionic acid, and bacteriocins, and some even propose its use as a probiotic^19^. We found a total of 2,341 OTUs classified as *Propionibacterium*, with 1,358 OTUs best matching *P. acnes* and *P. granulosum* (105 OTUs), as well as 878 unknown *Propionibacterium* species. The large number of *Propionibacterium* OTUs is probably reflecting an ongoing evolutionary process between bacteria and humans. Further work is needed to understand the human microbe partnership in healthy skin.

*Corynebacterium* had more OTUs (4,238), albeit in lower abundances than *Propionibacterium*. It was possible to identify two potential pathogens to the species level: *C. mastitidis* (15 OTUs) and *C. pilosum* (1 OTU), which cause subclinical mastitis and cystitis, respectively (Oliveira et al. 2017). We also found multiple *Corynebacterium* species: *C. simulans* (3 OTUs)*, C. lubricantis* (5 OTUs)*, C. stationis* (25 OTUs)*, C. durum* (53 OTUs), and *C. variabile* (70 OTUs). Although there are no reports of pathogenesis-related to those species, *C. variable* has been found in ripened cheeses^20^. However, most OTUs (4,066) were only classified to the genus level. The vast abundance of *Propionibacterium* and *Corynebacterium* is due to the prolific host microenvironments of the sebaceous glands, with their secreted fatty acids and a low pH (~5) and O2 concentration^21^. The biotechnological potential for further study in the Mexico City’s metro *Actinobacteria* is astounding: with 160 known genera detected, follow up projects could use this diversity to mine for new antibiotics and natural products, explore the potential for biomass decomposition from dust (which is mainly human skin debris in the subway), explore their possible role in bioenergy production, and further investigate *Actinobacteria* species, which are known to be active members of microcommunities and act as symbionts to eukaryotes^18^. It is also interesting that, together, *Actinobacteria* and *Firmicutes* comprise almost ~75% of the total described diversity (**Fig. 4**). *Firmicutes* is represented by 154 known genera, with a higher abundance of *Streptococcus, Staphylococcus, Leuconostoc, Lactobacillus, Veillonella, Anaerococcus*, and *Bacillus*.

The fifth most abundant biological group in Mexico city’s subway was classified into “chloroplast sequences”, which are 1,129 OTUs identified only as Streptophyta (land plants). We were also able to identify unicellular algae (Stramenopiles and Chlorophyta). A second search matching those OTUs to RDP matcher allowed a detailed classification of the plants in the metro system using the mitochondrial ribosomal genes (**Suppl. Table S4**), allowing us to identify plants rooted in the Mexican cultural identity, like maize and its wild relative teosintes (*Zea*), common bean (*Phaseolus vulgaris*), some edible plants like cucumber (*Cucumis sativus*), papaya (*Carica papaya*), lettuce (*Lactuca sativa*), sunflower (*Helianthus annuus*), and pea (*Pisum sativum*); in addition, recreative plants like coffee (*Coffea arabica*) and tobacco (*Nicotiana tabacum*) were found in our dataset. Other species included the most notorious introduced plant of Mexico City’s roadsides, *Eucalyptus globulus*, firs (*Abies*), abele (*Populus alba*), pine (*Pinus*), red cedar (*Juniperus virginiana*), ornamental plants like cycads (*Cycas taitungensis*), and all common plants growing in the city or its surroundings. Some of these plant genera have local centers of diversity or are being regularly consumed by Mexico’s population. It was unexpected to find a plant signal in our study, but we used vigorous DNA extraction and physical bead-beating (MoBio’s PowerSoil) to extract DNA from the surfaces and the deposited bioaerosols. The plant debris, pollen, and plant-derived products we discovered in this study give us a glimpse of the biodiversity of Mexico City. Studying macro species with metagenomic environmental DNA has allowed to track rare species of vertebrates^22^, invading plant species^23^, and lotic communities^24^. Further work on metagenomic DNA degradation in the subway will be useful to determine the spatiotemporal dynamics of microbe transmission and DNA resilience of other taxa, such as the plants species we detected in the metro.

In this paper, we do not intend to identify pathogenic bacteria, because we believe the sequencing of amplicons of the 16S rRNA gene is insufficient for this purpose. In multiple microbiome studies, it is common to define pathogenic groups, and there is even a dysbiosis index that is Bray-Curtis dissimilarity applied to microbiomes^25^. There is a tendency for studies to attempt to distinguish between good and bad or normal and abnormal microbial diversity, and the microbiome is termed “dysbiotic” when it is divergent from supposedly “normal” conditions^26^. When multiple microbiome papers refer to pathogenic bacteria, this is based in reductionistic assumptions that a given bacteria name is unequivocally causing a disease. However, determining pathogenesis is not a linear process. There are reports of typically “non-pathogenic” microbes leading to emerging opportunistic diseases, and of microbes that can be either commensal, innocuous, or pathogenic, e.g., *Escherichia coli* or *Candida albicans*. Categorizing host-microbe interactions into vague definitions like “pathogen” is a shortcoming of current microbiome research^27^. So, we are cautious to define pathogens and invite our readers to be aware of the limitations of microbiome studies involving disease agent identification.

Mexico City’s species pool has the metro as its playground. The species pool is defined as the microbes residing independently of their hosts, capable of colonizing and interacting with other inhabitants and hosts^28^. Species pools can be a part of any given community, and their success is measured by the similarity of species within a community, their frequency, dispersal abilities, and dormancy capabilities^29^. It has been stated that developed countries are reducing their microbiota diversity, compared to non-urban or Occidental lifestyles^30^. There is evidence that the microbiota diversity is convergent and reduced in migrant populations^31^, and changes in the microbiome may be used to track historical migrations^32^. The species pool is involved in the dispersal of microbes between their hosts and the environment, and it is shaped by the host and environmental feedback (*i.e*., immune system responses, host population density, continuous dispersal) that can actively shape the microbial community^28^. We propose that the large OTU number reported in this study (50,174) reflects both the species pool, host diversity, and the *γ* diversity (regional). However, it shows a homogeneous distribution at the genus level, reflecting the outcome of selective feedback between microbes, the environment, and hosts.

Further work could explore the diversity of Mexico City’s microbial pool and its impact on human health. The pool could be used to introduce microbiota to humans during the first years of life, to train and mature the immune system – a process that has been observed with newborn babies who have extensive variation in their skin microbiomes during the first years that subsequently stabilizes^33^.

## Conclusions

Mexico City’s metro provides a fantastic opportunity to test microbiome theories from multiple perspectives, from public health and epidemiology to metacommunity ecology and environment-host interactions that structure the collective microbiome. We did not observe significant differences in the microbiomes regarding people influx, environmental variables, nor architectural design. Actinobacteria dominated the metro surface diversity, and out of the 1,058 bacterial genera found in this study, 420 genera were ubiquitous in all lines and accounted for 99.10% of the total abundance, indicating a homogeneous system at the genus level and higher diversity at OTU level. The most abundant bacterial genus found in Mexico City’s metro surfaces were derived from normal human skin microbiota, with a dominance of *Actinobacteria* phylum (*i.e., Propionibacterium, Corynebacterium, Streptococcus, Staphylococcus*). The wide diversity of Mexico City’s metro microbiome is a delicate balance between environmental and host selection, and it is a major forum for bacterial species to be shared among the residents of the city. This is reflected in the homogeneous genus diversity but the wide heterogeneous diversity at the OTU level.

## Materials and Methods

### Sampling

Station turnstiles from the entrance and vertical handrails inside the train were sampled, each swab collected samples from 3 different bars, covering approximately 300 cm^2^ (100 cm^2^ from each bar). We sampled turnstiles from 24 stations, covering most of the of the subway stations. A total of 23 samples were collected from handrails, at chest level, from two trains on the same line. Sampling was performed with sterile cotton swabs, moistened with transport media (Tris 20mM, EDTA 10mM pH 7.5). Sampling permits were granted by the metro “User Support Manager” (Gerencia de Atención al Usuario, del Sistema de Transporte Colectivo).

### Metadata

Temperature and relative humidity were recorded for the sampled site. Information about the train building architectural design (underground, elevated, superficial trains or trains in tunnels), train wheel type (rubber tyre or steel wheels), and station information, such as climate zone, type of station (way, transfer, or terminal station), train environmental exposure level (open or subterraneous space) was recorded. Additionally, the environmental context of the exterior of the subway stations (work-related, bus stop, or park area) was also collected. People influx statistical data were obtained from the subway office reports, “Sistema de Transporte Colectivo: Cifras de Operación” (https://www.metro.cdmx.gob.mx/operacion/cifras-de-operacion).

### People behavior observations

Frequency of contact with handrail and turnstile bars was determined by counting the number of users touching a specific bar within 20 minutes (N = 8 for turnstiles, N = 9 for handrails). This procedure was performed for six different subway lines and seven different stations, respectively, during rush hour. Additionally, the type of contact that people had with turnstile bars was determined by counting the occurrence of two events: when the turnstile bar was pushed directly with the hand and pushed without hands (N = 730 people in 5 different stations), in rush hours.

### Amplicon generation and massive sequencing

Metagenomic DNA was obtained using the MoBio PowerSoil Kit (MoBio Laboratories, Solana Beach, CA), with the small modifications suggested by MoBio for low biomass samples. Briefly, 125 μL of the sample, 30 μL of C1 solution, and 50 μL phenol: chloroform 1:1 were mixed in the beat tube. The following steps followed the fabricant instructions. We used PCR primers for the V3-V4 region of the 16S rRNA gene, following MiSeq™ Illumina^®^ protocol with 5’ overhangs for sequencing library preparation (341F and 805R) with an expected amplicon size of 464 bp. Each PCR was done in triplicate with a mix reaction of 25 μL including 0.2 μL Phusion DNA polymerase™, 5 μL Buffer 5x, 2.5 μL dNTP (3 mM), 1 μL forward primer (5 pmol/μL), 1 μL reverse primer at (5 pmol/μL), 2 μL DNA, and 13.3 μL water. The PCR reaction was initiated at 98°C, 30s, followed by 35 cycles of 92°C for 10s; 53°C for 30s, 72°C for 40s, and the reaction finished at 72°C for 5 min. Blank samples were used for negative controls. PCR amplicons were purified using the High Pure PCR Product Purification Kit of ROCHE™ (Roche Diagnostics Gmb, Mannheim, Germany). Sequencing was performed using MiSeq™ Illumina^®^ (2 x 300 bp) platform at the Biotechnology Institute of the Universidad Nacional Autónoma de México (UNAM).

### Sequence processing

We used a previously reported protocol for 16S rRNA gene amplicons analysis^34^, which is detailed at GitHub (https://genomica-fciencias-unam.github.io/SOP/). Briefly, sequencing pair-ended reads were merged, using a minimum of 470 bp, a 15 bp of minimum overlapping, and with a quality threshold of 0.95 using PANDASEQ^35^. Sequence clustering into OTUs was performed at 97% identity with cd-hit-est^36^, cd-hit-est is an included OTU picking software in the popular QIIME scripts^37^, which we have previously tested as a reasonable option when picking *de novo* OTUs^34^. Representative sequences and OTU table were built with QIIME (v 1.9) ^37^. Taxonomy assignment was completed with parallel BLAST^38^ against the Greengenes database^39^. Plant-derived sequences were re-classified using the ribosomal database project^40^.

### Diversity metrics and statistical analysis

Diversity calculation and statistical inferences were performed with the R packages “phyloseq2” and “ggplot2” ^41,42^ or using R default functions^43^. Beta diversity ordination analyzes were evaluated with Canonical Analysis of Principal coordinates (CAP) for weighted and unweighted UniFrac distances at the OTU level, exploring the correlation with the following variables: sample_type + temperature_C + Humidity + length_underground + train_track + exposed_open_environment. PERMANOVA were performed per variable using the ANOVA function with 999 permutations. Sample dispersion was also evaluated with *betadisper* R function. The source tracker algorithm^44^ was used at the genus level, using 16S rRNA sequences (V4 region) from mattress dust, soil, saliva, skin, and fecal samples^45^ and vaginal samples^46^ as reference sources. Discriminant taxa analyzes were performed with R’s package “DESeq2” ^47^.

## Acknowledgments

This work was supported by the Secretary of Science, Technology, and Innovation of Mexico City SECITI 102/2017, Autonomous Metropolitan University, and National Council of Science and Technology Graduate fellowship (AMH).

