## Supplementary material for "Station and train surface microbiomes of Mexico City’s metro (subway/underground)": Tables S1, S2. Figures S1, S2, S4, S5, S6, S7

**Running title:** Mexico City’s metro surface microbiome

Apolinar Misael Hernández[a], Daniela Vargas-Robles[a,b], Luis D. Alcaraz[c], Mariana Peimbert[a#]

[a] Universidad Autónoma de Metropolitana, Unidad Cuajimalpa, México City, México
[b] Servicio Autónomo Centro Amazónico de Investigación y Control de Enfermedades Tropicales Simón Bolívar, MPPS, Puerto Ayacucho, Venezuela
[c] Departamento de Biología Celular, Facultad de Ciencias, Universidad Nacional Autónoma de México, México City, México

**Table S1.** Sequencing effort, pair-end merged sequences, and OTUs summary.

| | |
|---|---|
| N samples | 47 |
| Reads | 16,631,760 |
| Merged sequences | 5,788,162 |
| Average sequences/sample | 123,152.38 ± 28,371.93 |
| OTUs | 50,174 |
| Genus | 1,058 |

**Table S2. Alfa diversity (OTUs and Genera)**

| Metric | Level | Site | Min. | Median | Mean | Max. | *Median (p value*)* | *Dispersion (p value**)* |
|---|---|---|---|---|---|---|---|---|
| **Observed** | **OTUs** | Handrail | 5417.00 | 9433.00 | 9254.22 | 13152.00 | *0.081* | *0.001* |
| | | Turnstile | 428.00 | 10895.50 | 10525.83 | 16179.00 | | |
| | **Genus** | Handrail | 373.00 | 460.00 | 454.91 | 574.00 | *0.001* | *0.094* |
| | | Turnstile | 156.00 | 534.00 | 509.58 | 639.00 | | |
| **Chao1** | **OTUs** | Handrail | 8395.82 | 15260.65 | 14629.63 | 18649.01 | *0.148* | *0.001* |
| | | Turnstile | 840.25 | 16785.23 | 15631.35 | 23052.14 | | |
| | **Genus** | Handrail | 406.00 | 493.62 | 498.99 | 639.70 | *0.003* | *0.337* |
| | | Turnstile | 237.05 | 572.83 | 547.51 | 688.52 | | |
| **Shannon** | **OTUs** | Handrail | 5.54 | 6.27 | 6.24 | 7.38 | *0.024* | *0.106* |
| | | Turnstile | 4.10 | 6.72 | 6.52 | 7.36 | | |
| | **Genus** | Handrail | 2.47 | 3.41 | 3.40 | 4.03 | *4.00E-05* | *9.02E-01* |
| | | Turnstile | 3.00 | 4.00 | 3.91 | 4.44 | | |
| **Simpson** | **OTUs** | Handrail | 0.96 | 0.98 | 0.98 | 0.99 | *0.004* | *0.929* |
| | | Turnstile | 0.90 | 0.99 | 0.98 | 1.00 | | |
| | **Genus** | Handrail | 0.75 | 0.91 | 0.90 | 0.95 | *3.50E-04* | *1.34E-01* |
| | | Turnstile | 0.86 | 0.95 | 0.94 | 0.97 | | |

*Kruskal-Wallis test

**Fligner-Killeen Test of Homogeneity of Variances

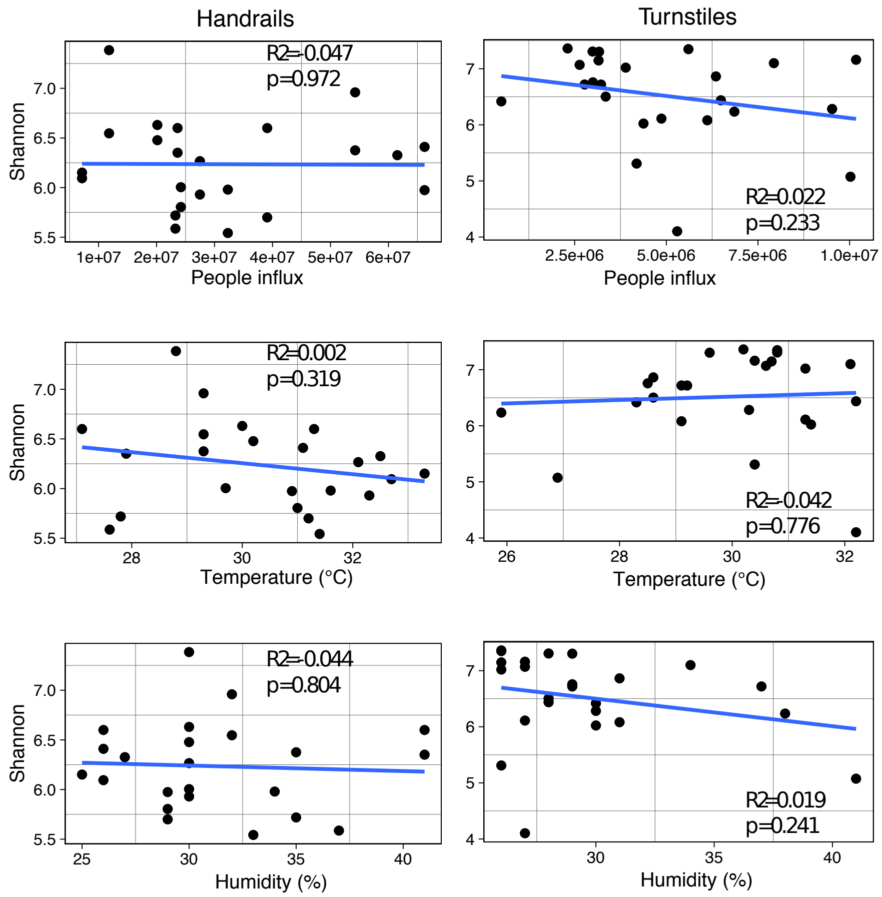


**Fig. S1. Association between temperature, humidity, or people influx with Shannon diversity in handrails and turnstiles.** A linear model was performed for each comparison. The mean temperature and humidity of the sampled points were 30.2 °C and 30.4%. There was no difference between turnstiles and handrail sampling points ($p > 0.05$, Kruskal-Wallis).

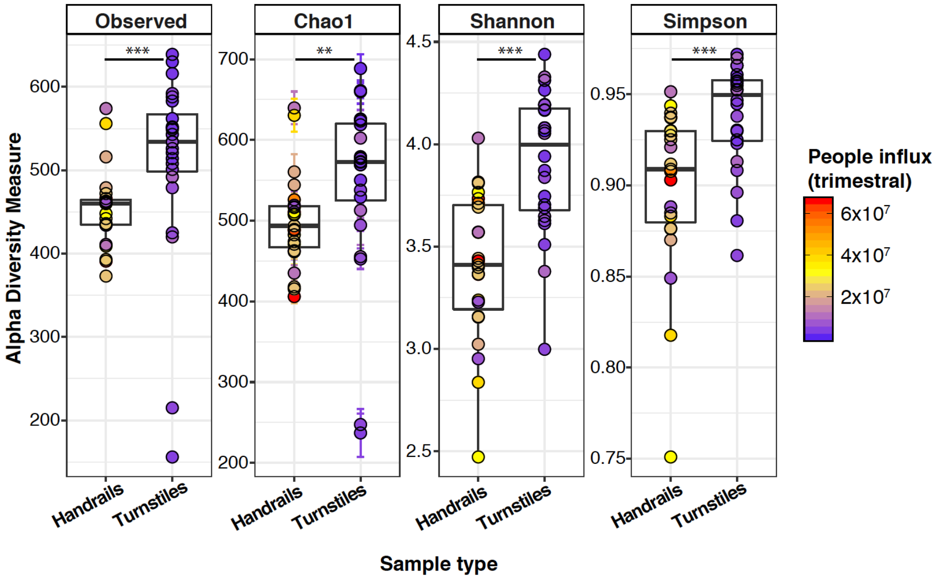


**Fig. S2**. **Alpha diversity measures of turnstile and handrails microbiota at the genus level.** Observed genera, Chao1, Shannon, and Simpson diversity indexes showed that turnstiles have higher microbial diversity than handrails (**p < 0.01, ***p < 0.001, Kruskal-Wallis).

**Fig. S3.** Heatmap of the 420 shared genera and their abundance in each sampled metro line.

Table S3. Genera abundances

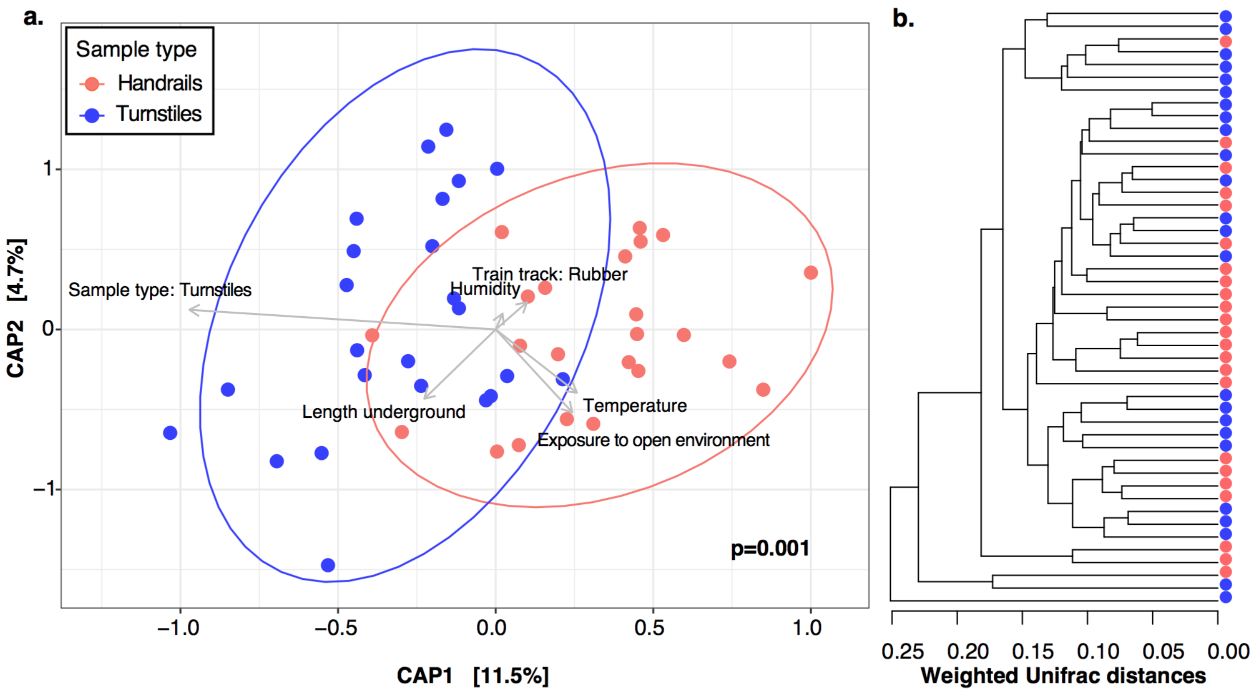


**Fig. S4. Beta diversity based on weighted UniFrac distances for turnstiles and handrails at the OTU level. a.** Canonical Analysis of Principal Coordinates (CAP) "sample type" significantly segregated (p = 0.001, Adonis), ellipses denote the 95% confidence interval of the points distribution by sample type. **b.** Dendrogram tends to group samples by "sample type."

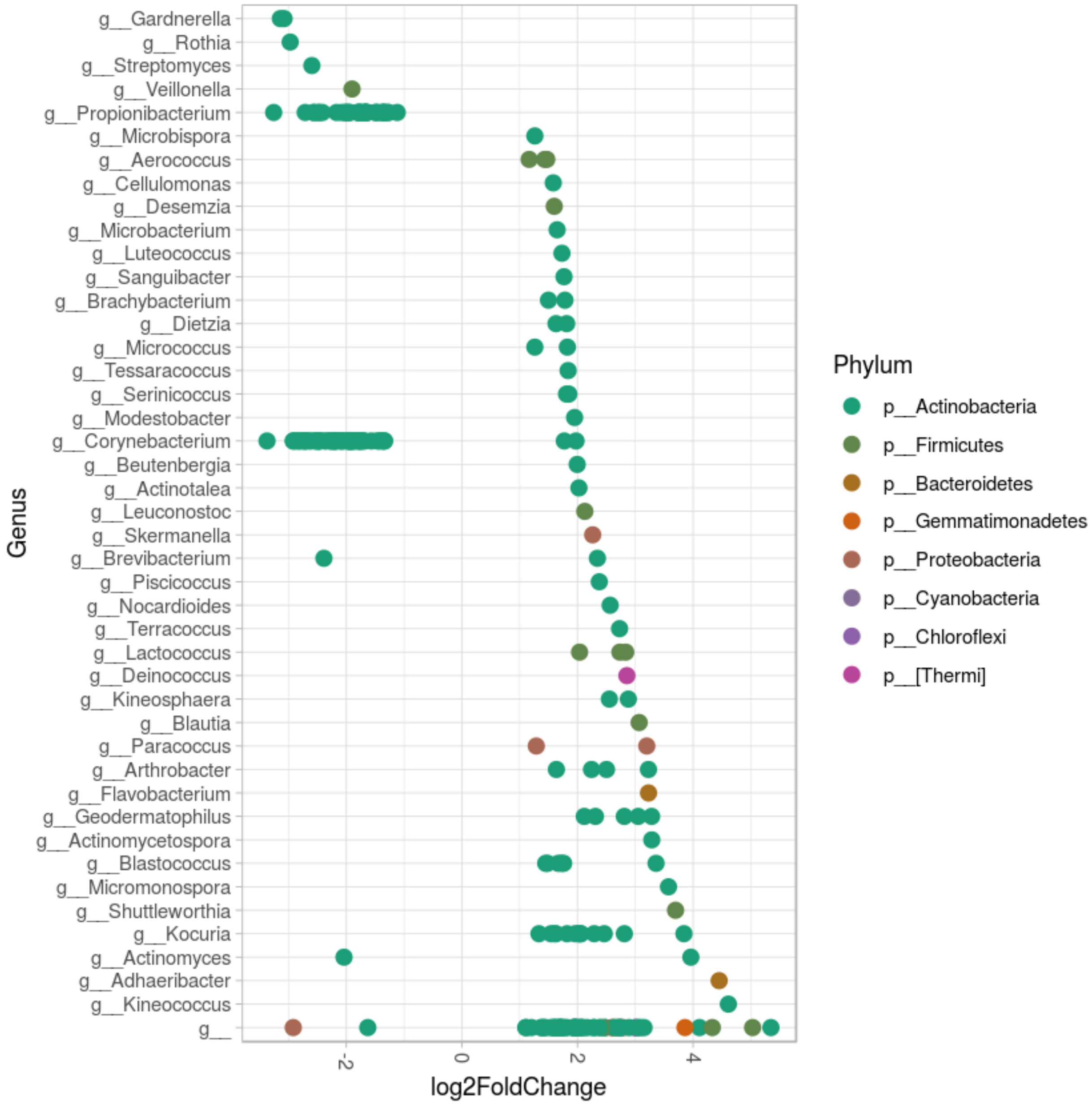


**Fig. S5. Differential OTUs abundances between metro handrails and turnstiles with a majority of *Actinobacteria*.** Significant (p-adj < 0.01) OTUs enriched in turnstiles (negative) or handrails (positive log2 fold change). Each dot is an OTU (97% identity 16S rRNA gene clusters).

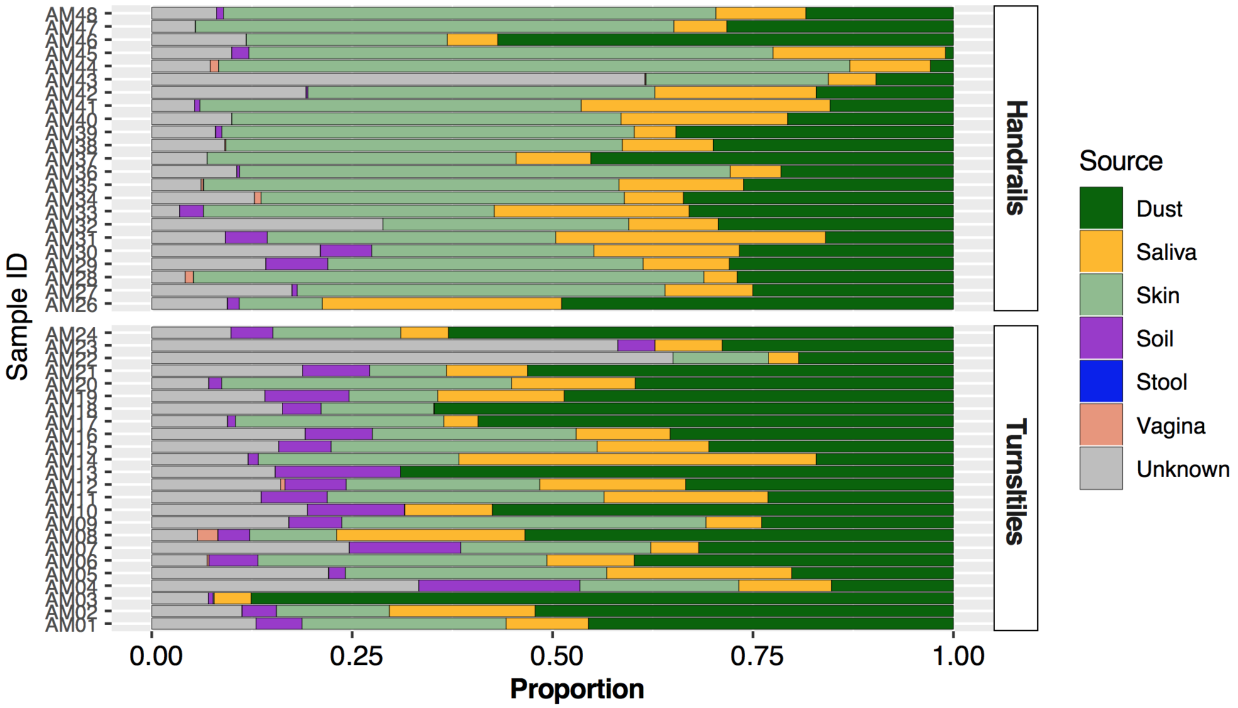


**Fig. S6. Source tracking of turnstile and handrail microbiota.** SourceTracker algorithm was performed at the genus level using dust, soil, saliva, skin, vagina, and feces samples as a source (DNA extraction for streamlined metagenomics of diverse environmental samples, DOI 10.2144/000114559). A substantial proportion of microbiota appears to be from skin and dust.

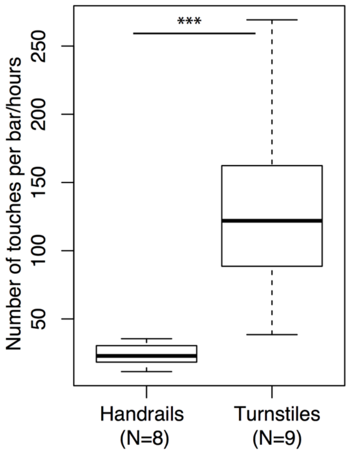


**Fig S7. Handrails and turnstiles users touch rate at peak hour.** The subway users touched turnstiles at a higher rate than handrails (***p = 0.001, Kruskal-Wallis test). More than 4 million users make contact with the turnstile every day to enter the subway's installations. This influx of data comes from the average number of users who go through all the turnstiles, per station, per day. This estimator allows us to compare the inflow of the different trains and stations, but it does not help us to understand how many people touched a train handrail or turnstile. We observed that 78% (570/730) of people touched the turnstiles directly with their hands, while the rest touched with their clothes or carry-on items. Estimating train handrail use proved to be complicated, since some people touch many handrails, some hold tight to just one, and others do not touch anything. However, we observed that the rate of direct contact with one of the turnstiles bars is significantly higher (median, 122 people/hour, N = 8) than with one of the vertical train bars (median 22.9 people/hour, p = 0.001, Kruskal Wallis, N=9), both data were recorded during rush hour.
