## Supplementary figures and images for "Station and train surface microbiomes of Mexico City’s metro (subway/underground)"

### Supplemental Figure S3

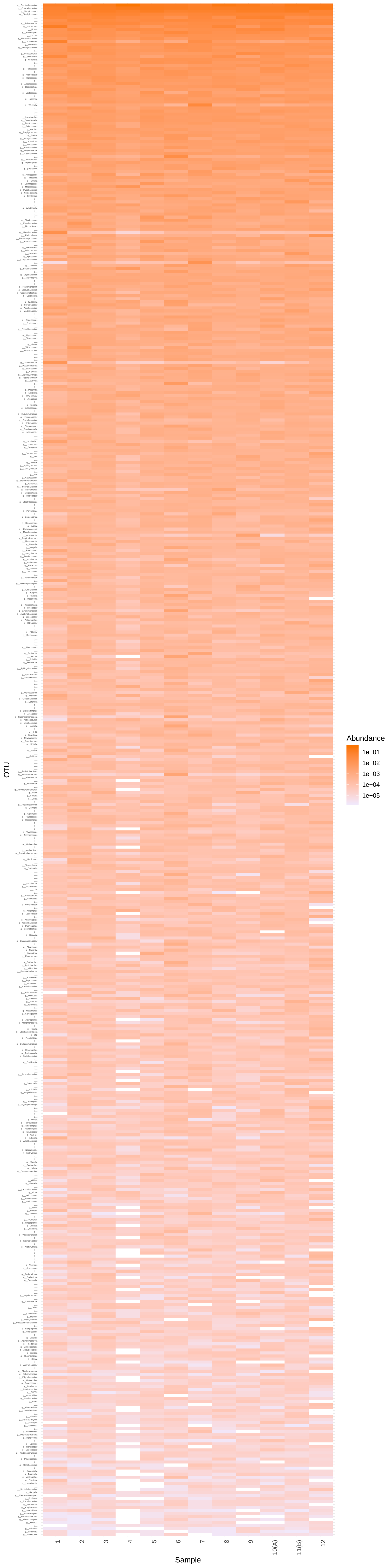
