## Supplemental Table S3 for "Station and train surface microbiomes of Mexico City’s metro (subway/underground)"

| OTU | 1 | 12 | 2 | 3 | 4 | 5 | 6 | 7 | 8 | 9 | A | B | Kingdom | Phylum | Class | Order | Family | Genus | Abundance |
|---|---|---|---|---|---|---|---|---|---|---|---|---|---|---|---|---|---|---|---|
| 414180 | 0.10344245 | 0.15406558 | 0.0875378 | 0.14886856 | 0.29213253 | 0.13888377 | 0.12590533 | 0.09303756 | 0.20900644 | 0.16049254 | 0.09584527 | 0.16585031 | k__Bacteria | p__Actinobact | c__Actinobact | o__Actinomycetales | f__Propionibacteria | g__Propionibacterium | 1.78E+00 |
| 416710 | 0.07090474 | 0.160575 | 0.13495511 | 0.12262001 | 0.16338809 | 0.14742909 | 0.12782521 | 0.10050703 | 0.11891777 | 0.13285929 | 0.11169916 | 0.1349697 | k__Bacteria | p__Actinobact | c__Actinobact | o__Actinomycetales | f__Corynebacteriace | g__Corynebacterium | 1.53E+00 |
| 128 | 0.06432112 | 0.05547658 | 0.08424772 | 0.10320739 | 0.07822082 | 0.1200955 | 0.07391863 | 0.08206616 | 0.07958622 | 0.11064285 | 0.09326702 | 0.11445913 | k__Bacteria | p__Firmicutes | c__Bacilli | o__Lactobacillales | f__Streptococcacea | g__Streptococcus | 1.06E+00 |
| 196 | 0.05858584 | 0.05767756 | 0.04077542 | 0.06592877 | 0.05950694 | 0.04630942 | 0.03932568 | 0.04706649 | 0.05859549 | 0.05003758 | 0.05014696 | 0.04593471 | k__Bacteria | p__Firmicutes | c__Bacilli | o__Bacillales | f__Staphylococcace | g__Staphylococcus | 6.20E-01 |
| 577669 | 0.04324757 | 0.02370887 | 0.03369463 | 0.04881399 | 0.02983373 | 0.03279204 | 0.01489024 | 0.04042043 | 0.01667072 | 0.06660018 | 0.07397936 | 0.01768643 | k__Bacteria | p__Cyanobact | c__Chloroplas | o__Streptophyta | f__ | g__ | 4.42E-01 |
| 406268 | 0.0192957 | 0.03846418 | 0.04879575 | 0.03218915 | 0.02417694 | 0.03348508 | 0.03958082 | 0.0333161 | 0.03157213 | 0.02345358 | 0.02806073 | 0.03308991 | k__Bacteria | p__Actinobact | c__Actinobact | o__Actinomycetales | f__Micrococcaceae | g__ | 3.85E-01 |
| 16 | 0.02905893 | 0.04155366 | 0.02436273 | 0.03924985 | 0.03679124 | 0.02328346 | 0.01834412 | 0.0251531 | 0.02424997 | 0.02229738 | 0.03902767 | 0.02740566 | k__Bacteria | p__Proteobac | c__Gammapro | o__Pseudomonadales | f__Moraxellaceae | g__Acinetobacter | 3.51E-01 |
| 198 | 0.11234454 | 0.0045859 | 0.01172017 | 0.02317642 | 0.01456816 | 0.00722919 | 0.02208502 | 0.02307988 | 0.04743714 | 0.00698347 | 0.02836075 | 0.01322329 | k__Bacteria | p__Proteobac | c__Gammapro | o__Oceanospirillales | f__Halomonadacea | g__Halomonas | 3.15E-01 |
| 429324 | 0.01209881 | 0.01457449 | 0.03596803 | 0.0139754 | 0.02747028 | 0.02389384 | 0.01245053 | 0.0191393 | 0.0184218 | 0.0332524 | 0.05363932 | 0.01724457 | k__Bacteria | p__Actinobact | c__Actinobact | o__Actinomycetales | f__Micrococcaceae | g__Rothia | 2.82E-01 |
| 106592 | 0.01274627 | 0.01237351 | 0.01440504 | 0.01327029 | 0.00760511 | 0.01852758 | 0.03741856 | 0.0207714 | 0.01491558 | 0.02561568 | 0.01765867 | 0.01554563 | k__Bacteria | p__Actinobact | c__Actinobact | o__Actinomycetales | f__Actinomycetacea | g__Actinomyces | 2.11E-01 |
| 407005 | 0.0135946 | 0.00764732 | 0.01812203 | 0.01325876 | 0.00907742 | 0.01103771 | 0.01746391 | 0.01362786 | 0.01414227 | 0.01111978 | 0.01312564 | 0.01382271 | k__Bacteria | p__Actinobact | c__Actinobact | o__Actinomycetales | f__Micrococcaceae | g__Kocuria | 1.56E-01 |
| 423705 | 0.00730587 | 0.02762138 | 0.01530343 | 0.0095699 | 0.00586158 | 0.01537713 | 0.0054503 | 0.01366217 | 0.01255516 | 0.00975546 | 0.01154353 | 0.00706806 | k__Bacteria | p__Proteobac | c__Alphaprote | o__Rhizobiales | f__Methylobacteriac | g__Methylobacterium | 1.41E-01 |
| 2752 | 0.07923416 | 0.00491947 | 0.00280317 | 0.00341796 | 0.00073062 | 0.00371633 | 0.00104924 | 0.02369254 | 0.00568444 | 0.00254076 | 0.00111568 | 0.00235317 | k__Bacteria | p__Firmicutes | c__Bacilli | o__Lactobacillales | f__Leuconostocace | g__Leuconostoc | 1.31E-01 |
| 363378 | 0.00654515 | 0.0096519 | 0.01038974 | 0.01165449 | 0.00731729 | 0.00925426 | 0.01721834 | 0.01295395 | 0.01112798 | 0.0120274 | 0.00956765 | 0.0127232 | k__Bacteria | p__Bacteroide | c__Bacteroidia | o__Bacteroidales | f__Prevotellaceae | g__Prevotella | 1.30E-01 |
| 411630 | 0.00734647 | 0.01255121 | 0.01851979 | 0.00730087 | 0.00506454 | 0.00715925 | 0.01462554 | 0.01414249 | 0.00669258 | 0.00915424 | 0.0108099 | 0.01019363 | k__Bacteria | p__Actinobact | c__Actinobact | o__Actinomycetales | f__Dermabacterace | g__Brachybacterium | 1.24E-01 |
| 407471 | 0.00794906 | 0.00853893 | 0.01724936 | 0.00929516 | 0.00677486 | 0.00829736 | 0.01076983 | 0.01402241 | 0.00894976 | 0.00732166 | 0.0098161 | 0.01030838 | k__Bacteria | p__Actinobact | c__Actinobact | o__Actinomycetales | f__ | g__ | 1.19E-01 |
| 919 | 0.00950041 | 0.0112169 | 0.007614 | 0.01015589 | 0.00560697 | 0.00466369 | 0.00548538 | 0.00551633 | 0.00737479 | 0.02564458 | 0.01430694 | 0.00837823 | k__Bacteria | p__Proteobac | c__Gammapro | o__Pseudomonadales | f__Pseudomonadac | g__Pseudomonas | 1.15E-01 |
| 2872 | 0.00648532 | 0.00695834 | 0.00583265 | 0.00788878 | 0.00688556 | 0.01254777 | 0.0124601 | 0.01168943 | 0.00850439 | 0.00964851 | 0.00827384 | 0.0101491 | k__Bacteria | p__Firmicutes | c__Clostridia | o__Clostridiales | f__Veillonellaceae | g__Veillonella | 1.07E-01 |
| 441149 | 0.0051968 | 0.00985765 | 0.01195505 | 0.00550063 | 0.00425089 | 0.01420406 | 0.01085275 | 0.01372589 | 0.00580793 | 0.00512198 | 0.01091537 | 0.0093767 | k__Bacteria | p__Actinobact | c__Actinobact | o__Actinomycetales | f__Nocardioidaceae | g__ | 1.07E-01 |
| 237 | 0.0340271 | 0.00202952 | 0.00360555 | 0.00690316 | 0.00516417 | 0.00206639 | 0.00737656 | 0.00725872 | 0.01831653 | 0.00225171 | 0.00917154 | 0.00481594 | k__Bacteria | p__Proteobac | c__Gammapro | o__Alteromonadales | f__Shewanellaceae | g__Shewanella | 1.03E-01 |
| 5842 | 0.01202829 | 0.00762861 | 0.00411132 | 0.00816545 | 0.00683574 | 0.00372586 | 0.00325614 | 0.00508257 | 0.00630592 | 0.01018904 | 0.00535104 | 0.01746036 | k__Bacteria | p__Proteobac | c__Gammapro | o__Enterobacteriales | f__Enterobacteriace | g__ | 9.01E-02 |
| 448273 | 0.00465832 | 0.00658112 | 0.00606239 | 0.00727781 | 0.00291142 | 0.01178797 | 0.00609451 | 0.00962356 | 0.0078485 | 0.00409585 | 0.01356159 | 0.00778395 | k__Bacteria | p__Proteobac | c__Alphaprote | o__Rhodobacterales | f__Rhodobacterace | g__Paracoccus | 8.83E-02 |
| 441124 | 0.00358349 | 0.00795596 | 0.00947592 | 0.00650738 | 0.00309407 | 0.00698758 | 0.0113407 | 0.00715334 | 0.00444148 | 0.00415655 | 0.01181073 | 0.0059754 | k__Bacteria | p__Actinobact | c__Actinobact | o__Actinomycetales | f__Intrasporangiace | g__ | 8.25E-02 |
| 405577 | 0.0059853 | 0.00565833 | 0.00911759 | 0.00704342 | 0.00401842 | 0.00658066 | 0.01088783 | 0.0081924 | 0.00617434 | 0.00494566 | 0.00584794 | 0.00707491 | k__Bacteria | p__Actinobact | c__Actinobact | o__Actinomycetales | f__Micrococcaceae | g__Arthrobacter | 8.15E-02 |
| 431705 | 0.00286764 | 0.00672765 | 0.00675847 | 0.01055936 | 0.00571213 | 0.00717515 | 0.00715969 | 0.0040043 | 0.0084133 | 0.00613366 | 0.00441115 | 0.00768461 | k__Bacteria | p__Actinobact | c__Actinobact | o__Actinomycetales | f__Micrococcaceae | g__Micrococcus | 7.76E-02 |
| 621816 | 0.00886149 | 0.00532475 | 0.00263344 | 0.00623456 | 0.01110324 | 0.00492437 | 0.00370263 | 0.00459735 | 0.00678975 | 0.00483582 | 0.00803711 | 0.00438436 | k__Bacteria | p__Firmicutes | c__Clostridia | o__Clostridiales | f__[Tissierellaceae] | g__Anaerococcus | 7.14E-02 |
| 4011 | 0.00526305 | 0.00401227 | 0.00451079 | 0.00989652 | 0.00531361 | 0.00537262 | 0.00552684 | 0.00513894 | 0.00705292 | 0.00629842 | 0.00482836 | 0.00673239 | k__Bacteria | p__Firmicutes | c__Bacilli | o__Bacillales | f__Planococcaceae | g__ | 6.99E-02 |
| 1567 | 0.0066584 | 0.00471995 | 0.00436335 | 0.00412307 | 0.00304979 | 0.00594803 | 0.00563208 | 0.00626377 | 0.0086299 | 0.00846052 | 0.00439943 | 0.0040247 | k__Bacteria | p__Proteobac | c__Gammapro | o__Pasteurellales | f__Pasteurellaceae | g__Haemophilus | 6.63E-02 |
| 581015 | 0.0022736 | 0.00630678 | 0.00844552 | 0.00337761 | 0.0026236 | 0.00612924 | 0.00579791 | 0.00790323 | 0.00341309 | 0.0024049 | 0.00919732 | 0.00553868 | k__Bacteria | p__Chloroflexi | c__Thermomic | o__JG30-KF-CM45 | f__ | g__ | 6.34E-02 |
| 536 | 0.01251122 | 0.00289931 | 0.01079778 | 0.0036562 | 0.00218633 | 0.01150821 | 0.0009727 | 0.00813114 | 0.00241913 | 0.00269684 | 0.00139225 | 0.0030211 | k__Bacteria | p__Firmicutes | c__Bacilli | o__Lactobacillales | f__Streptococcacea | g__Lactococcus | 6.22E-02 |
| 3713 | 0.00597889 | 0.00358205 | 0.00798432 | 0.00834989 | 0.00267341 | 0.00222217 | 0.00540246 | 0.00336469 | 0.00327341 | 0.00941727 | 0.00303062 | 0.0058538 | k__Bacteria | p__Proteobac | c__Gammapro | o__Pseudomonadales | f__Moraxellaceae | g__ | 6.11E-02 |
| 6905 | 0.00374589 | 0.00701446 | 0.00471653 | 0.0042518 | 0.00301658 | 0.00431399 | 0.00715331 | 0.00422486 | 0.0055812 | 0.00774656 | 0.00316187 | 0.00499748 | k__Bacteria | p__Proteobac | c__Betaprotec | o__Neisseriales | f__Neisseriaceae | g__Neisseria | 5.99E-02 |
| 487684 | 0.00244882 | 0.0039468 | 0.00459137 | 0.00679173 | 0.00386898 | 0.00399926 | 0.00513776 | 0.00552123 | 0.00455282 | 0.00544283 | 0.00450959 | 0.00687967 | k__Bacteria | p__Firmicutes | c__Clostridia | o__Clostridiales | f__Lachnospiraceae | g__ | 5.77E-02 |
| 332 | 0.00134194 | 0.00159306 | 0.00096525 | 0.00240544 | 0.001976 | 0.00170398 | 0.00034443 | 0.040293 | 0.00076116 | 0.00215632 | 0.0048049 | 0.00093853 | k__Bacteria | p__Firmicutes | c__Bacilli | o__Lactobacillales | f__Leuconostocace | g__Weissella | 5.50E-02 |
| 594606 | 0.00206846 | 0.00404345 | 0.00340324 | 0.00579651 | 0.00287821 | 0.00732456 | 0.0032402 | 0.0055972 | 0.00527349 | 0.0024367 | 0.00703862 | 0.00544106 | k__Bacteria | p__Proteobac | c__Alphaprote | o__Rhodobacterales | f__Rhodobacterace | g__ | 5.45E-02 |
| 0 | 0.00650883 | 0.0035384 | 0.0037187 | 0.00376187 | 0.00324352 | 0.00549978 | 0.00504527 | 0.00416849 | 0.00461557 | 0.00653255 | 0.004526 | 0.00314954 | k__Bacteria | p__Firmicutes | c__Bacilli | o__Gemellales | f__Gemellaceae | g__ | 5.43E-02 |
| 401629 | 0.00355571 | 0.00452043 | 0.00868897 | 0.00356206 | 0.00156087 | 0.003888 | 0.00446803 | 0.00718275 | 0.00446982 | 0.00287895 | 0.00482836 | 0.00447341 | k__Bacteria | p__Actinobact | c__Actinobact | o__Actinomycetales | f__Cellulomonadace | g__ | 5.41E-02 |
| 7743 | 0.00321595 | 0.00421179 | 0.00557205 | 0.00403085 | 0.00250736 | 0.00725144 | 0.00420333 | 0.00457285 | 0.00530791 | 0.00352064 | 0.00219854 | 0.00402984 | k__Bacteria | p__Firmicutes | c__Bacilli | o__Lactobacillales | f__Lactobacillaceae | g__Lactobacillus | 5.06E-02 |
| 5610 | 0.00277576 | 0.00258756 | 0.00381643 | 0.00418839 | 0.00311068 | 0.00623097 | 0.00454138 | 0.00431063 | 0.00611968 | 0.00598913 | 0.0035158 | 0.00330026 | k__Bacteria | p__Firmicutes | c__Bacilli | o__Lactobacillales | f__Carnobacteriace | g__Granulicatella | 5.05E-02 |
| 415588 | 0.00206633 | 0.00343864 | 0.00665217 | 0.00236125 | 0.00107933 | 0.00460964 | 0.00398965 | 0.00553839 | 0.00300012 | 0.00321713 | 0.00555027 | 0.00442032 | k__Bacteria | p__Actinobact | c__Actinobact | o__Actinomycetales | f__Geodermatophila | g__Blastococcus | 4.59E-02 |
| 2 | 0.00230565 | 0.00566457 | 0.00506114 | 0.00393478 | 0.00366971 | 0.0030519 | 0.00499105 | 0.00304366 | 0.00242925 | 0.00298589 | 0.00385097 | 0.00367361 | k__Bacteria | p__Firmicutes | c__Bacilli | o__Bacillales | f__Bacillaceae | g__Bacillus | 4.47E-02 |
| 447167 | 0.00133766 | 0.00301466 | 0.00547433 | 0.00238047 | 0.0018653 | 0.00398019 | 0.00729683 | 0.00495759 | 0.00268432 | 0.00174298 | 0.00408301 | 0.00329683 | k__Bacteria | p__Actinobact | c__Actinobact | o__Actinomycetales | f__Dietziaceae | g__Dietzia | 4.21E-02 |
| 55238 | 0.00291893 | 0.0035384 | 0.00376156 | 0.0029338 | 0.00311621 | 0.00429809 | 0.00259599 | 0.00413174 | 0.00342929 | 0.00368251 | 0.00394238 | 0.00324031 | k__Bacteria | p__Bacteroide | c__Bacteroidia | o__Bacteroidales | f__Porphyromonada | g__Porphyromonas | 4.16E-02 |
| 418998 | 0.00106842 | 0.00348541 | 0.00819177 | 0.00380029 | 0.00103505 | 0.00400244 | 0.0031796 | 0.00315149 | 0.00300215 | 0.00283848 | 0.00233449 | 0.00526123 | k__Bacteria | p__[Thermi] | c__Deinococci | o__Deinococcales | f__Deinococcaceae | g__Deinococcus | 4.14E-02 |
| 464306 | 0.00219668 | 0.00299907 | 0.00241913 | 0.00236125 | 0.0020701 | 0.00310595 | 0.00451906 | 0.00344311 | 0.00239079 | 0.00567696 | 0.003783 | 0.00433983 | k__Bacteria | p__Fusobacte | c__Fusobacte | o__Fusobacteriales | f__Leptotrichiaceae | g__Leptotrichia | 3.93E-02 |
| 5363 | 0.00068806 | 0.00240986 | 0.00475768 | 0.00559477 | 0.0026236 | 0.00148462 | 0.00755515 | 0.0013797 | 0.0034718 | 0.002376 | 0.00464554 | 0.00222472 | k__Bacteria | p__Firmicutes | c__Bacilli | o__Bacillales | f__Staphylococcace | g__Jeotgalicoccus | 3.92E-02 |
| 597541 | 0.00530151 | 0.0019017 | 0.00170419 | 0.00058983 | 0.0004262 | 0.00096326 | 0.02315977 | 0.00083321 | 0.00084821 | 0.00055209 | 0.00182353 | 0.00076384 | k__Bacteria | p__Firmicutes | c__Clostridia | o__Clostridiales | f__Clostridiaceae | g__ | 3.89E-02 |
| 417274 | 0.00149793 | 0.00363817 | 0.00421076 | 0.00298183 | 0.00133947 | 0.00220945 | 0.00668769 | 0.00311228 | 0.00222884 | 0.00247716 | 0.00332125 | 0.00434668 | k__Bacteria | p__Actinobact | c__Actinobact | o__Actinomycetales | f__Brevibacteriacea | g__Brevibacterium | 3.81E-02 |
| 85 | 0.00226505 | 0.00734492 | 0.00296262 | 0.00384256 | 0.00222508 | 0.00315363 | 0.00142237 | 0.00202911 | 0.00363173 | 0.002376 | 0.00258528 | 0.0036719 | k__Bacteria | p__Proteobac | c__Gammapro | o__Pseudomonadales | f__Moraxellaceae | g__Enhydrobacter | 3.75E-02 |
| 7642 | 0.00437626 | 0.00195781 | 0.00730025 | 0.0033315 | 0.00119556 | 0.0016245 | 0.00292766 | 0.00333529 | 0.00291105 | 0.00313042 | 0.00200166 | 0.00246106 | k__Bacteria | p__Firmicutes | c__Bacilli | o__Lactobacillales | f__Aerococcaceae | g__Aerococcus | 3.66E-02 |
| 526192 | 0.00288688 | 0.00300219 | 0.00161675 | 0.00232091 | 0.00133394 | 0.00326172 | 0.00581705 | 0.00428122 | 0.00313778 | 0.00288473 | 0.00284779 | 0.00260321 | k__Bacteria | p__Fusobacte | c__Fusobacte | o__Fusobacteriales | f__Fusobacteriacea | g__Fusobacterium | 3.60E-02 |
| 436075 | 0.00179067 | 0.00254703 | 0.00456394 | 0.00162925 | 0.00107933 | 0.00483536 | 0.00328485 | 0.00371023 | 0.00165189 | 0.00200312 | 0.00279857 | 0.00370615 | k__Bacteria | p__Actinobact | c__Actinobact | o__Actinomycetales | f__Cellulomonadace | g__Cellulomonas | 3.36E-02 |
| 618776 | 0.00317108 | 0.0023101 | 0.00194936 | 0.00285502 | 0.00396307 | 0.00231754 | 0.00163923 | 0.00263686 | 0.00344548 | 0.00258122 | 0.00364236 | 0.0022487 | k__Bacteria | p__Firmicutes | c__Clostridia | o__Clostridiales | f__[Tissierellaceae] | g__Peptoniphilus | 3.28E-02 |
| 16922 | 0.00306637 | 0.0034667 | 0.00125843 | 0.0033853 | 0.00160515 | 0.00197738 | 0.0036803 | 0.00178895 | 0.00370258 | 0.00341658 | 0.00201807 | 0.00194727 | k__Bacteria | p__Proteobac | c__Betaprotec | o__Neisseriales | f__Neisseriaceae | g__ | 3.13E-02 |
| 5405 | 0.00334202 | 0.01007276 | 0.00201623 | 0.00185212 | 0.00083025 | 0.0012144 | 0.00076221 | 0.00569033 | 0.00099194 | 0.00116487 | 0.00072425 | 0.00167839 | k__Bacteria | p__Firmicutes | c__Bacilli | o__Lactobacillales | f__Leuconostocace | g__ | 3.03E-02 |
| 365341 | 0.00223086 | 0.00158059 | 0.00199394 | 0.00239007 | 0.00138929 | 0.0020505 | 0.00383976 | 0.00185757 | 0.00604275 | 0.00324893 | 0.0014696 | 0.00208086 | k__Bacteria | p__Bacteroide | c__Bacteroidia | o__Bacteroidales | f__[Paraprevotellace | g__[Prevotella] | 3.02E-02 |
| 613011 | 0.00237831 | 0.00281825 | 0.00130815 | 0.00292419 | 0.00322138 | 0.00266723 | 0.00140642 | 0.00206342 | 0.00264383 | 0.00245693 | 0.0023673 | 0.00216478 | k__Bacteria | p__Firmicutes | c__Clostridia | o__Clostridiales | f__[Tissierellaceae] | g__Finegoldia | 2.84E-02 |
| 48372 | 0.00050002 | 0.0007638 | 0.00093953 | 0.00577921 | 0.00034317 | 0.00658066 | 0.00209848 | 0.0045802 | 0.00234625 | 0.00089317 | 0.00280326 | 0.00073644 | k__Bacteria | p__Firmicutes | c__Bacilli | o__Lactobacillales | f__Aerococcaceae | g__Alloiococcus | 2.84E-02 |
| 4427 | 0.00221377 | 0.00123143 | 0.00349068 | 0.00337761 | 0.01005712 | 0.00075344 | 0.00104286 | 0.00086752 | 0.00166201 | 0.00111285 | 0.00129381 | 0.00119028 | k__Bacteria | p__Firmicutes | c__Bacilli | o__Bacillales | f__Staphylococcace | g__Macrococcus | 2.83E-02 |
| 405472 | 0.00108765 | 0.0064003 | 0.00212767 | 0.00140446 | 0.00088007 | 0.00783003 | 0.00165199 | 0.00144831 | 0.00102231 | 0.00135854 | 0.00125162 | 0.00162187 | k__Bacteria | p__Actinobact | c__Actinobact | o__Actinomycetales | f__Dermacoccaceae | g__Dermacoccus | 2.81E-02 |
| 447452 | 0.00199367 | 0.00281514 | 0.00326608 | 0.00125652 | 0.00169925 | 0.00253371 | 0.00218458 | 0.00315149 | 0.0017855 | 0.00160712 | 0.00217276 | 0.00182739 | k__Bacteria | p__Actinobact | c__Actinobact | o__Actinomycetales | f__Mycobacteriacea | g__Mycobacterium | 2.63E-02 |
| 609633 | 0.0012992 | 0.00162735 | 0.00276202 | 0.00101059 | 0.00100184 | 0.00310912 | 0.00658245 | 0.00253884 | 0.00124701 | 0.00086715 | 0.00140397 | 0.00229322 | k__Bacteria | p__Firmicutes | c__Clostridia | o__Clostridiales | f__Clostridiaceae | g__Clostridium | 2.57E-02 |
| 405883 | 0.00101927 | 0.001618 | 0.0020608 | 0.00215184 | 0.00128966 | 0.00144012 | 0.0017668 | 0.0021786 | 0.00401028 | 0.00125737 | 0.00389316 | 0.00189075 | k__Bacteria | p__Actinobact | c__Actinobact | o__Actinomycetales | f__Micrococcaceae | g__Nesterenkonia | 2.46E-02 |
| 1391 | 0.0018441 | 0.00163359 | 0.00126357 | 0.00358703 | 0.0009963 | 0.00082338 | 0.00183377 | 0.00087732 | 0.00188267 | 0.00435021 | 0.00298374 | 0.00199694 | k__Bacteria | p__Proteobac | c__Gammapro | o__Pseudomonadales | f__Pseudomonadac | g__ | 2.41E-02 |
| 1302 | 0.001827 | 0.00123766 | 0.00081266 | 0.00126997 | 0.00045941 | 0.00703209 | 0.00081324 | 0.0053791 | 0.00149399 | 0.00168517 | 0.00079457 | 0.00089229 | k__Bacteria | p__Firmicutes | c__Bacilli | o__Lactobacillales | f__ | g__ | 2.37E-02 |
| 1891 | 0.00177572 | 0.00207628 | 0.00129615 | 0.00207114 | 0.00398521 | 0.00088696 | 0.00110345 | 0.00149978 | 0.00342726 | 0.00118222 | 0.00256653 | 0.00174861 | k__Bacteria | p__Proteobac | c__Gammapro | o__Aeromonadales | f__Aeromonadacea | g__ | 2.36E-02 |
| 6802 | 0.00302791 | 0.00128754 | 0.00115727 | 0.00241697 | 0.00128412 | 0.00103955 | 0.00066973 | 0.00143361 | 0.00154865 | 0.00137588 | 0.00114381 | 0.00686597 | k__Bacteria | p__Proteobac | c__Gammapro | o__Enterobacteriales | f__Enterobacteriace | g__Erwinia | 2.33E-02 |
| 424461 | 0.00132912 | 0.00178011 | 0.0020848 | 0.00092029 | 0.00060332 | 0.00440618 | 0.00525894 | 0.00147282 | 0.0009717 | 0.00124003 | 0.00071957 | 0.00161844 | k__Bacteria | p__Actinobact | c__Actinobact | o__Actinomycetales | f__Nocardiaceae | g__Rhodococcus | 2.24E-02 |
| 374772 | 0.00097654 | 0.00056116 | 0.00197508 | 0.00114508 | 0.00045941 | 0.00112221 | 0.00142875 | 0.00194089 | 0.00107292 | 0.00232975 | 0.00125162 | 0.00774113 | k__Bacteria | p__Actinobact | c__Actinobact | o__Actinomycetales | f__Actinomycetacea | g__ | 2.20E-02 |
| 363484 | 0.00325228 | 0.00090097 | 0.00220139 | 0.00216913 | 0.00114575 | 0.00134792 | 0.00132988 | 0.00261235 | 0.0016114 | 0.00131229 | 0.00199228 | 0.00167667 | k__Bacteria | p__Bacteroide | c__Flavobacte | o__Flavobacteriales | f__[Weeksellaceae] | g__Wautersiella | 2.16E-02 |
| 613047 | 0.00060045 | 0.00206069 | 0.00409589 | 0.00070703 | 0.00095756 | 0.00159271 | 0.00313496 | 0.00189923 | 0.00114175 | 0.0010666 | 0.00260169 | 0.00166982 | k__Bacteria | p__Firmicutes | c__Clostridia | o__Clostridiales | f__Peptostreptococ | g__ | 2.15E-02 |
| 5137 | 0.00089961 | 0.01078044 | 0.00065493 | 0.00201158 | 0.00035978 | 0.00063581 | 0.00060913 | 0.00064206 | 0.00115592 | 0.00031507 | 0.00168524 | 0.0009214 | k__Bacteria | p__Proteobac | c__Gammapro | o__Alteromonadales | f__[Chromatiaceae] | g__Rheinheimera | 2.07E-02 |
| 7312 | 0.00290611 | 0.00021199 | 0.00112641 | 0.00234781 | 0.0017546 | 0.00068032 | 0.00183377 | 0.00231093 | 0.00305883 | 0.00091918 | 0.00216807 | 0.00130846 | k__Bacteria | p__Proteobac | c__Gammapro | o__Oceanospirillales | f__Halomonadacea | g__ | 2.06E-02 |
| 416077 | 0.00083978 | 0.00238803 | 0.00309806 | 0.00084536 | 0.00065313 | 0.0019583 | 0.00173491 | 0.00269813 | 0.00111543 | 0.00083536 | 0.00251497 | 0.00153966 | k__Bacteria | p__Actinobact | c__Actinobact | o__Actinomycetales | f__Nocardioidaceae | g__Nocardioides | 2.02E-02 |
| 9569 | 0.00136758 | 0.00241609 | 0.00141787 | 0.00220371 | 0.00041513 | 0.00170716 | 0.00175086 | 0.00141891 | 0.00141706 | 0.0024049 | 0.00150476 | 0.00173833 | k__Bacteria | p__Proteobac | c__Betaprotec | o__Burkholderiales | f__Oxalobacteracea | g__ | 1.98E-02 |
| 426444 | 0.00044233 | 0.00102255 | 0.00151217 | 0.00212878 | 0.00628224 | 0.00082974 | 0.00102054 | 0.00130863 | 0.00100206 | 0.00076309 | 0.00125162 | 0.00137354 | k__Bacteria | p__Actinobact | c__Actinobact | o__Bifidobacteriales | f__Bifidobacteriacea | g__Bifidobacterium | 1.89E-02 |
| 420344 | 0.00152784 | 0.00075444 | 0.00295062 | 0.0007224 | 0.00041513 | 0.00641217 | 0.00087064 | 0.00144831 | 0.00061541 | 0.00158978 | 0.00050393 | 0.00095565 | k__Bacteria | p__Actinobact | c__Actinobact | o__Actinomycetales | f__Intrasporangiace | g__Arsenicicoccus | 1.88E-02 |
| 363726 | 0.00079491 | 0.00166788 | 0.00764314 | 0.00125268 | 0.00050369 | 0.00072801 | 0.00128524 | 0.00109297 | 0.00092514 | 0.00052896 | 0.00090708 | 0.00118343 | k__Bacteria | p__Bacteroide | c__Flavobacte | o__Flavobacteriales | f__Flavobacteriacea | g__Flavobacterium | 1.85E-02 |

| | | | | | | | | | | | | | | | | | | | |
|---|---|---|---|---|---|---|---|---|---|---|---|---|---|---|---|---|---|---|---|
| 619366 | 0.00064319 | 0.00059545 | 0.00217396 | 0.00143904 | 0.00079704 | 0.00133839 | 0.00227707 | 0.00161986 | 0.0012794 | 0.00206093 | 0.00262747 | 0.00127592 | k__Bacteria | p__Firmicutes | c__Clostridia | o__Clostridiales | f__Peptostreptococ | g__Peptostreptococcus | 1.81E-02 |
| 270 | 0.01614385 | 0.000106 | 0.00015259 | 8.45E-05 | 1.11E-05 | 5.40E-05 | 0.00029021 | 0.00036024 | 0.00017207 | 0.00017921 | 7.97E-05 | 0.00011132 | k__Bacteria | p__Proteobac | c__Gammapro | o__Vibrionales | f__Vibrionaceae | g__Photobacterium | 1.77E-02 |
| 318 | 0.00076713 | 0.00087603 | 0.00111098 | 0.00107592 | 0.00068634 | 0.00163722 | 0.00303609 | 0.00128657 | 0.00104053 | 0.00236154 | 0.00192431 | 0.00158248 | k__Bacteria | p__Firmicutes | c__Clostridia | o__Clostridiales | f__Veillonellaceae | g__Selenomonas | 1.74E-02 |
| 201 | 0.00387196 | 0.00074509 | 0.00090867 | 0.00196355 | 0.00055904 | 0.00124937 | 0.00149891 | 0.00128902 | 0.00166201 | 0.00165337 | 0.00057893 | 0.00138895 | k__Bacteria | p__Firmicutes | c__Bacilli | o__Lactobacillales | f__Streptococcacea | g__ | 1.74E-02 |
| 420264 | 0.0002842 | 0.00190482 | 0.00266773 | 0.00154663 | 0.00125091 | 0.00315363 | 0.00086427 | 0.00146547 | 0.00119236 | 0.00087582 | 0.00067972 | 0.00144889 | k__Bacteria | p__Actinobact | c__Actinobact | o__Actinomycetales | f__Intrasporangiace | g__Kytococcus | 1.73E-02 |
| 848 | 0.00098936 | 0.00213551 | 0.0009241 | 0.00100483 | 0.00047048 | 0.00100776 | 0.00465619 | 0.001017 | 0.00092514 | 0.00172274 | 0.00124928 | 0.00122625 | k__Bacteria | p__Firmicutes | c__Bacilli | o__Lactobacillales | f__Aerococcaceae | g__ | 1.73E-02 |
| 363489 | 0.00187187 | 0.00158371 | 0.00096868 | 0.00112971 | 0.00029336 | 0.00213633 | 0.00108113 | 0.00308532 | 0.00131787 | 0.00126315 | 0.00119303 | 0.001043 | k__Bacteria | p__Bacteroide | c__Flavobacte | o__Flavobacteriales | f__[Weeksellaceae] | g__Chryseobacterium | 1.70E-02 |
| 609729 | 0.00043592 | 0.00081056 | 0.006347 | 0.00081847 | 0.00032103 | 0.0018216 | 0.00137453 | 0.00146547 | 0.00035224 | 0.0006995 | 0.00103364 | 0.00103101 | k__Bacteria | p__Proteobac | c__Alphaprote | o__Rhodospirillales | f__Rhodospirillacea | g__Skermanella | 1.65E-02 |
| 407618 | 0.00168597 | 0.00175206 | 0.00236941 | 0.00101444 | 0.00146678 | 0.00134474 | 0.00107156 | 0.00171543 | 0.00072473 | 0.00078044 | 0.00110865 | 0.00132387 | k__Bacteria | p__Actinobact | c__Actinobact | o__Actinomycetales | f__Microbacteriacea | g__Cryobacterium | 1.64E-02 |
| 430883 | 0.00048079 | 0.00151512 | 0.00147788 | 0.00054372 | 0.0006476 | 0.00069622 | 0.00398009 | 0.00156839 | 0.00091502 | 0.00047982 | 0.000668 | 0.00304679 | k__Bacteria | p__Actinobact | c__Actinobact | o__Actinomycetales | f__Gordoniaceae | g__Gordonia | 1.60E-02 |
| 436581 | 0.00118595 | 0.00109737 | 0.00177106 | 0.00137179 | 0.00124538 | 0.00128752 | 0.00136497 | 0.00139195 | 0.00132596 | 0.00106949 | 0.00101255 | 0.00151226 | k__Bacteria | p__Actinobact | c__Actinobact | o__Actinomycetales | f__Micrococcaceae | g__Microbispora | 1.56E-02 |
| 431267 | 6.41E-06 | 8.11E-05 | 0.00064464 | 0.00012873 | 3.87E-05 | 9.22E-05 | 0.00014032 | 0.01345877 | 0.00028949 | 6.36E-05 | 0.00027423 | 9.76E-05 | k__Bacteria | p__Cyanobact | c__Oscillatorio | o__Chroococcales | f__Xenococcaceae | g__ | 1.53E-02 |
| 609800 | 0.0004466 | 0.00201081 | 0.00158075 | 0.00115853 | 0.00112361 | 0.00058177 | 0.00131713 | 0.00264666 | 0.0005648 | 0.00061279 | 0.00182587 | 0.00126564 | k__Bacteria | p__Firmicutes | c__Clostridia | o__Clostridiales | f__Ruminococcacea | g__ | 1.51E-02 |
| 7516 | 0.00079491 | 0.00338565 | 0.00097554 | 0.00136219 | 0.00049815 | 0.00070257 | 0.00095356 | 0.0012155 | 0.00150411 | 0.00066771 | 0.00208838 | 0.0009625 | k__Bacteria | p__Firmicutes | c__Bacilli | o__Bacillales | f__[Exiguobacterac | g__Exiguobacterium | 1.51E-02 |
| 1768 | 0.00106842 | 0.00088538 | 0.00094125 | 0.00172147 | 0.00157748 | 0.00134792 | 0.00141918 | 0.00096309 | 0.00151221 | 0.00147416 | 0.00097505 | 0.00117145 | k__Bacteria | p__Firmicutes | c__Bacilli | o__Gemellales | f__ | g__ | 1.51E-02 |
| 12383 | 0.00055772 | 0.00057363 | 0.00253914 | 0.00100291 | 0.00202028 | 0.00032426 | 0.00382701 | 0.00083076 | 0.00079963 | 0.00061857 | 0.00107818 | 0.00062854 | k__Bacteria | p__Firmicutes | c__Bacilli | o__Lactobacillales | f__Aerococcaceae | g__Facklamia | 1.48E-02 |
| 88867 | 0.00049575 | 0.00130001 | 0.00290947 | 0.00078388 | 0.00074723 | 0.00085199 | 0.00242058 | 0.0012155 | 0.00096158 | 0.00093652 | 0.00086957 | 0.00125365 | k__Bacteria | p__Firmicutes | c__Bacilli | o__Bacillales | f__Planococcaceae | g__Planomicrobium | 1.47E-02 |
| 433202 | 0.0010321 | 0.00114725 | 0.00227511 | 0.00053219 | 0.00012731 | 0.00143376 | 0.00152442 | 0.00135764 | 0.00080368 | 0.00082669 | 0.00131725 | 0.00177601 | k__Bacteria | p__Actinobact | c__Actinobact | o__Actinomycetales | f__Geodermatophila | g__Geodermatophilus | 1.42E-02 |
| 5848 | 0.00209838 | 0.00075444 | 0.00044062 | 0.00084152 | 0.00035424 | 0.00042917 | 0.00035719 | 0.00071803 | 0.0006316 | 0.00070817 | 0.00033752 | 0.0062734 | k__Bacteria | p__Proteobac | c__Gammapro | o__Enterobacteriales | f__Enterobacteriace | g__Klebsiella | 1.39E-02 |
| 459305 | 0.00338049 | 0.00089162 | 0.00104069 | 0.0009568 | 0.00156641 | 0.00085835 | 0.00051984 | 0.00066902 | 0.00109721 | 0.00055498 | 0.0013571 | 0.00103443 | k__Bacteria | p__Proteobac | c__Alphaprote | o__Rhizobiales | f__Rhizobiaceae | g__Agrobacterium | 1.39E-02 |
| 427741 | 0.00023292 | 0.00134366 | 0.0015036 | 0.0019501 | 0.0013118 | 0.00082974 | 0.00049751 | 0.00145812 | 0.0017855 | 0.00086715 | 0.00052268 | 0.00147287 | k__Bacteria | p__Actinobact | c__Actinobact | o__Bifidobacteriales | f__Bifidobacteriacea | g__Gardnerella | 1.38E-02 |
| 783 | 0.00139322 | 0.00046451 | 0.00223568 | 0.00226519 | 0.00117896 | 0.00022889 | 0.00307436 | 0.00024751 | 0.00024495 | 0.00065904 | 0.00026954 | 0.00143177 | k__Bacteria | p__Proteobac | c__Gammapro | o__Pseudomonadales | f__Moraxellaceae | g__Psychrobacter | 1.37E-02 |
| 586158 | 0.000406 | 0.00118155 | 0.00081609 | 0.00083384 | 0.00065313 | 0.00114764 | 0.0027395 | 0.00195314 | 0.00094133 | 0.00125448 | 0.00096567 | 0.00070732 | k__Bacteria | p__TM7 | c__TM7-3 | o__ | f__ | g__ | 1.36E-02 |
| 610992 | 0.00112825 | 0.0004801 | 0.00223396 | 0.00060328 | 0.00018266 | 0.0010173 | 0.00087702 | 0.00105621 | 0.00079963 | 0.00066482 | 0.00353923 | 0.00088886 | k__Bacteria | p__Proteobac | c__Alphaprote | o__Rhodospirillales | f__Acetobacteracea | g__ | 1.35E-02 |
| 582162 | 0.00157272 | 0.00052998 | 0.0013973 | 0.0003881 | 0.00118449 | 0.00119851 | 0.00032211 | 0.00189923 | 0.00055873 | 0.00084114 | 0.00294155 | 0.00040932 | k__Bacteria | p__Proteobac | c__Alphaprote | o__Rickettsiales | f__mitochondria | g__ | 1.32E-02 |
| 436034 | 0.00101073 | 0.00061104 | 0.00170419 | 0.00050145 | 0.00028229 | 0.00139879 | 0.00112578 | 0.00117629 | 0.00067209 | 0.00092496 | 0.00062816 | 0.003117 | k__Bacteria | p__Actinobact | c__Actinobact | o__Actinomycetales | f__Geodermatophila | g__Modestobacter | 1.32E-02 |
| 423061 | 0.00088252 | 0.00105373 | 0.001255 | 0.00079349 | 0.0003321 | 0.0014274 | 0.00126291 | 0.00102681 | 0.00084821 | 0.00071396 | 0.00236261 | 0.00116802 | k__Bacteria | p__Actinobact | c__Actinobact | o__Actinomycetales | f__Intrasporangiace | g__Serinicoccus | 1.31E-02 |
| 610501 | 0.00292107 | 0.00102255 | 0.00082981 | 0.00094335 | 0.0003155 | 0.00065171 | 0.00060594 | 0.00079155 | 0.00122272 | 0.00070528 | 0.00159148 | 0.00137182 | k__Bacteria | p__Proteobac | c__Alphaprote | o__Sphingomonadales | f__Sphingomonada | g__ | 1.30E-02 |
| 609840 | 0.00026069 | 0.00368493 | 0.00129786 | 0.00117198 | 0.00078044 | 0.00055952 | 0.0001754 | 0.0021884 | 0.00049597 | 0.00031507 | 0.00085551 | 0.00109438 | k__Bacteria | p__Firmicutes | c__Clostridia | o__Clostridiales | f__Ruminococcacea | g__Faecalibacterium | 1.29E-02 |
| 440949 | 0.00074148 | 0.00066092 | 0.00068579 | 0.00087226 | 0.00078044 | 0.00139879 | 0.0022388 | 0.00097044 | 0.00125106 | 0.00124292 | 0.00050862 | 0.00122282 | k__Bacteria | p__Actinobact | c__Actinobact | o__Actinomycetales | f__Pseudonocardia | g__ | 1.26E-02 |
| 446671 | 0.00110902 | 0.00078874 | 0.0028066 | 0.00048801 | 0.00017712 | 0.00084245 | 0.00116405 | 0.00196294 | 0.000581 | 0.00067349 | 0.00102427 | 0.00094538 | k__Bacteria | p__Actinobact | c__Actinobact | o__Actinomycetales | f__Dermatophilacea | g__Piscicoccus | 1.26E-02 |
| 428390 | 0.00048079 | 0.00133119 | 0.00132529 | 0.00104902 | 0.00050369 | 0.00129706 | 0.00144151 | 0.00117875 | 0.00077331 | 0.0007313 | 0.00125631 | 0.00109095 | k__Bacteria | p__Actinobact | c__Actinobact | o__Actinomycetales | f__Intrasporangiace | g__Phycicoccus | 1.25E-02 |
| 409275 | 0.00043164 | 0.00180505 | 0.00148988 | 0.00120272 | 0.00034317 | 0.00082656 | 0.00180826 | 0.00111503 | 0.00045144 | 0.00066771 | 0.00129147 | 0.00081693 | k__Bacteria | p__Actinobact | c__Actinobact | o__Actinomycetales | f__Intrasporangiace | g__Terracoccus | 1.22E-02 |
| 609777 | 0.0004637 | 0.00148083 | 0.00173677 | 0.00118735 | 0.0009797 | 0.00074072 | 0.00044011 | 0.00181836 | 0.00077331 | 0.0004278 | 0.00105474 | 0.00096422 | k__Bacteria | p__Firmicutes | c__Clostridia | o__Clostridiales | f__Lachnospiraceae | g__Blautia | 1.21E-02 |
| 448507 | 0.00069234 | 0.00114725 | 0.00169905 | 0.00079349 | 0.00066974 | 0.0012144 | 0.00150848 | 0.00126452 | 0.000581 | 0.00066771 | 0.00065159 | 0.00113377 | k__Bacteria | p__Actinobact | c__Actinobact | o__Actinomycetales | f__Nocardioidaceae | g__Aeromicrobium | 1.20E-02 |
| 431910 | 0.00067738 | 0.00135925 | 0.00162532 | 0.00054949 | 0.00033764 | 0.0008806 | 0.0011768 | 0.00149978 | 0.000581 | 0.00079489 | 0.00124459 | 0.00113034 | k__Bacteria | p__Actinobact | c__Actinobact | o__Actinomycetales | f__Geodermatophila | g__ | 1.19E-02 |
| 15009 | 0.00042737 | 0.00164606 | 0.00129443 | 0.00119504 | 0.00075276 | 0.00078841 | 0.00093124 | 0.00077194 | 0.0013786 | 0.00079778 | 0.00063284 | 0.0012194 | k__Bacteria | p__Proteobac | c__Betaprotec | o__Burkholderiales | f__Comamonadacea | g__ | 1.18E-02 |
| 586284 | 0.00030557 | 0.00134054 | 0.00148817 | 0.00077043 | 0.00070295 | 0.00096008 | 0.00152761 | 0.00075234 | 0.00104053 | 0.00095387 | 0.00112271 | 0.00067992 | k__Bacteria | p__Firmicutes | c__Clostridia | o__Clostridiales | f__[Mogibacteriacea | g__ | 1.16E-02 |
| 658768 | 0.00054703 | 0.00066715 | 0.00114356 | 0.00127573 | 0.00059778 | 0.00056269 | 0.00192945 | 0.00122286 | 0.00071258 | 0.00069661 | 0.00104771 | 0.00090256 | k__Bacteria | p__Firmicutes | c__Clostridia | o__Clostridiales | f__ | g__ | 1.13E-02 |
| 431472 | 0.00038036 | 0.00078562 | 0.00136301 | 0.00055333 | 0.00040959 | 0.00102048 | 0.00096632 | 0.00272018 | 0.00055265 | 0.00049717 | 0.00066331 | 0.00097449 | k__Bacteria | p__Actinobact | c__Actinobact | o__Actinomycetales | f__Pseudonocardia | g__Pseudonocardia | 1.09E-02 |
| 603515 | 0.00943416 | 0.00056739 | 5.31E-05 | 0.00021326 | 2.77E-05 | 2.23E-05 | 9.89E-05 | 8.82E-05 | 8.70E-05 | 0.00018788 | 1.41E-05 | 9.25E-05 | k__Bacteria | p__Proteobac | c__Alphaprote | o__Rhodospirillales | f__Acetobacteracea | g__Gluconobacter | 1.09E-02 |
| 12074 | 0.00032053 | 0.00014964 | 0.00524116 | 0.00040155 | 0.00046494 | 0.00020982 | 0.0020315 | 0.00036024 | 0.00044334 | 0.00034108 | 0.00040314 | 0.00037164 | k__Bacteria | p__Firmicutes | c__Bacilli | o__Lactobacillales | f__Carnobacteriace | g__Trichococcus | 1.07E-02 |
| 406908 | 0.00079704 | 0.0010662 | 0.0013973 | 0.0006225 | 0.00047601 | 0.00111585 | 0.00062827 | 0.00146302 | 0.00060731 | 0.00101457 | 0.00074066 | 0.00071246 | k__Bacteria | p__Actinobact | c__Actinobact | o__Actinomycetales | f__Microbacteriacea | g__Cryocola | 1.06E-02 |
| 8896 | 0.00015813 | 0.00061415 | 0.00107498 | 0.00124115 | 0.00024908 | 0.00080112 | 0.00243653 | 0.00090183 | 0.00136848 | 0.00026593 | 0.00079926 | 0.00062169 | k__Bacteria | p__Firmicutes | c__Bacilli | o__Bacillales | f__Staphylococcace | g__Salinicoccus | 1.05E-02 |
| 364283 | 0.00060686 | 0.00187988 | 0.00046977 | 0.0006321 | 0.00037085 | 0.00045143 | 0.00081324 | 0.0012155 | 0.00056885 | 0.00184703 | 0.00064691 | 0.00095908 | k__Bacteria | p__Bacteroide | c__Flavobacte | o__Flavobacteriales | f__Flavobacteriacea | g__Capnocytophaga | 1.05E-02 |
| 435657 | 0.00012394 | 0.00045828 | 0.00031375 | 0.00030741 | 0.0004428 | 0.00039738 | 0.00595418 | 0.00013478 | 0.00058909 | 0.00043069 | 0.00053206 | 0.00031855 | k__Bacteria | p__Actinobact | c__Coriobacte | o__Coriobacteriales | f__Coriobacteriacea | g__ | 1.00E-02 |
| 10386 | 0.00020514 | 0.00240362 | 0.00049206 | 0.00050914 | 0.00137822 | 0.00081384 | 0.00103329 | 0.00125226 | 0.00022673 | 0.00049428 | 0.00084379 | 0.00022436 | k__Bacteria | p__Proteobac | c__Gammapro | o__Pseudomonadales | f__Moraxellaceae | g__Moraxella | 9.88E-03 |
| 7998 | 0.00102782 | 0.00065157 | 0.00058978 | 0.00048416 | 0.00049262 | 0.00137971 | 0.0007367 | 0.00122286 | 0.00062148 | 0.00120534 | 0.00060472 | 0.00062511 | k__Bacteria | p__Proteobac | c__Gammapro | o__Pasteurellales | f__Pasteurellaceae | g__Aggregatibacter | 9.64E-03 |
| 2182 | 0.00054917 | 0.00156812 | 0.00119499 | 0.00039771 | 0.00037085 | 0.00065807 | 0.00068248 | 0.00092388 | 0.00070651 | 0.00116198 | 0.00044533 | 0.00071075 | k__Bacteria | p__Proteobac | c__Betaprotec | o__Burkholderiales | f__Burkholderiaceae | g__Lautropia | 9.37E-03 |
| 610051 | 0.00021155 | 0.0010132 | 0.0004732 | 0.00094911 | 0.00194279 | 0.00053726 | 0.00031573 | 0.00051708 | 0.00051217 | 0.00048271 | 0.00128444 | 0.00063539 | k__Bacteria | p__Firmicutes | c__Clostridia | o__Clostridiales | f__[Tissierellaceae] | g__WAL_1855D | 8.87E-03 |
| 18551 | 0.00116885 | 0.00096332 | 0.0004732 | 0.00055717 | 0.00027122 | 0.00052772 | 0.00049751 | 0.00049993 | 0.0007976 | 0.00093363 | 0.00058597 | 0.00129133 | k__Bacteria | p__Proteobac | c__Gammapro | o__Xanthomonadales | f__Xanthomonadac | g__ | 8.57E-03 |
| 6654 | 0.00075644 | 0.00077938 | 0.00122928 | 0.00074161 | 0.0002214 | 0.00055952 | 0.00052302 | 0.00070578 | 0.00081177 | 0.00077755 | 0.00042658 | 0.00065765 | k__Bacteria | p__Firmicutes | c__Bacilli | o__Lactobacillales | f__Carnobacteriace | g__Desemzia | 8.19E-03 |
| 447628 | 0.00024574 | 0.0006578 | 0.00080923 | 0.00046879 | 0.00019926 | 0.00128434 | 0.00064421 | 0.00072293 | 0.00043726 | 0.00059255 | 0.00117896 | 0.00070903 | k__Bacteria | p__Actinobact | c__Actinobact | o__Actinomycetales | f__Sporichthyaceae | g__ | 7.95E-03 |
| 471778 | 0.00024146 | 0.00032734 | 0.00045948 | 0.00101828 | 0.00072509 | 0.00053726 | 0.00056767 | 0.00109052 | 0.00084619 | 0.00068794 | 0.00064925 | 0.0006645 | k__Bacteria | p__Actinobact | c__Coriobacte | o__Coriobacteriales | f__Coriobacteriacea | g__Atopobium | 7.81E-03 |
| 437341 | 0.00035472 | 0.00066092 | 0.00061035 | 0.00089147 | 0.00028782 | 0.00045779 | 0.00137453 | 0.00066657 | 0.00045346 | 0.00041623 | 0.00109224 | 0.00050694 | k__Bacteria | p__Actinobact | c__Actinobact | o__Actinomycetales | f__Intrasporangiace | g__Knoellia | 7.77E-03 |
| 610168 | 0.00034189 | 0.00088226 | 0.0005332 | 0.00059368 | 9.41E-05 | 0.00104591 | 0.00067929 | 0.00102681 | 0.0004069 | 0.0006995 | 0.00040314 | 0.00091797 | k__Bacteria | p__Proteobac | c__Alphaprote | o__Rhodobacterales | f__Rhodobacterace | g__Rubellimicrobium | 7.62E-03 |
| 648 | 0.00057481 | 0.00045516 | 0.00062064 | 0.00087995 | 0.00054797 | 0.0002575 | 0.00061232 | 0.00048522 | 0.00059921 | 0.00058099 | 0.00128912 | 0.00064567 | k__Bacteria | p__Firmicutes | c__Bacilli | o__Lactobacillales | f__Enterococcaceae | g__Enterococcus | 7.55E-03 |
| 363540 | 0.00057695 | 0.0002307 | 0.00102354 | 0.00033815 | 0.0002214 | 0.00133203 | 0.00044011 | 0.00119835 | 0.00052836 | 0.00061279 | 0.00047815 | 0.00047954 | k__Bacteria | p__Bacteroide | c__Cytophagia | o__Cytophagales | f__Cytophagaceae | g__Hymenobacter | 7.46E-03 |
| 609698 | 0.00048079 | 0.00071703 | 0.00062921 | 0.00120656 | 0.00039852 | 0.00051819 | 0.0005007 | 0.00056364 | 0.00047775 | 0.00054052 | 0.00073832 | 0.00059771 | k__Bacteria | p__Proteobac | c__Alphaprote | o__Caulobacterales | f__Caulobacteracea | g__ | 7.37E-03 |
| 597451 | 0.00015599 | 0.00118778 | 0.00074065 | 0.00038426 | 0.00033764 | 0.00086788 | 0.00064102 | 0.00074254 | 0.00045144 | 0.00024569 | 0.00085317 | 0.00068163 | k__Bacteria | p__Actinobact | c__Acidimicrob | o__Acidimicrobiales | f__ | g__ | 7.29E-03 |
| 426939 | 0.0001624 | 0.00066715 | 0.0006035 | 0.00041308 | 0.00028782 | 0.00056905 | 0.00205064 | 0.00057099 | 0.00042309 | 0.00035553 | 0.00048284 | 0.00069191 | k__Bacteria | p__Actinobact | c__Actinobact | o__Actinomycetales | f__Streptomycetace | g__Streptomyces | 7.28E-03 |
| 13662 | 0.00055344 | 0.00039281 | 0.00036004 | 0.00052643 | 0.00135054 | 0.00043553 | 0.00071756 | 0.00053914 | 0.00078141 | 0.00022257 | 0.00081332 | 0.00051037 | k__Bacteria | p__Proteobac | c__Gammapro | o__Xanthomonadales | f__Xanthomonadac | g__Luteimonas | 7.20E-03 |
| 609978 | 0.00091884 | 0.00061104 | 0.00038919 | 0.00045342 | 0.00013284 | 0.00080748 | 0.00058681 | 0.00049747 | 0.00052431 | 0.00041623 | 0.00112271 | 0.00059429 | k__Bacteria | p__Proteobac | c__Alphaprote | o__Sphingomonadales | f__Sphingomonada | g__Kaistobacter | 7.05E-03 |
| 12146 | 0.00223514 | 0.00059233 | 0.00038747 | 0.0003612 | 0.00057564 | 0.00025433 | 0.00022005 | 0.00036514 | 0.00043524 | 0.00051451 | 0.00031173 | 0.00078439 | k__Bacteria | p__Proteobac | c__Gammapro | o__Enterobacteriales | f__Enterobacteriace | g__Enterobacter | 7.04E-03 |
| 410092 | 0.00037181 | 0.00033358 | 0.00109212 | 0.00046303 | 0.00013284 | 0.0008202 | 0.00065378 | 0.00128412 | 0.00047573 | 0.0005174 | 0.00031173 | 0.00052749 | k__Bacteria | p__Actinobact | c__Actinobact | o__Actinomycetales | f__Nocardioidaceae | g__Friedmanniella | 6.98E-03 |
| 24575 | 0.00050643 | 0.00048322 | 0.00112298 | 0.00045726 | 0.00028229 | 0.00037831 | 0.00059 | 0.00098515 | 0.00023483 | 0.00044225 | 0.00094458 | 0.00041446 | k__Bacteria | p__Firmicutes | c__Bacilli | o__Lactobacillales | f__Carnobacteriace | g__ | 6.84E-03 |
| 18766 | 0.00082696 | 0.0024005 | 0.0002366 | 0.00035352 | 0.00049262 | 0.00019074 | 0.00038589 | 0.00033818 | 0.00047978 | 0.00039889 | 0.00037971 | 0.00033396 | k__Bacteria | p__Proteobac | c__Betaprotec | o__Burkholderiales | f__Comamonadacea | g__Comamonas | 6.82E-03 |
| 424742 | 0.00025428 | 0.00091344 | 0.00060178 | 0.0004246 | 0.00035978 | 0.00047368 | 0.00130437 | 0.00036269 | 0.00056075 | 0.00034686 | 0.00060237 | 0.00055832 | k__Bacteria | p__Actinobact | c__Actinobact | o__Actinomycetales | f__Bogoriellaceae | g__Georgenia | 6.76E-03 |
| 12166 | 0.00171375 | 0.0004084 | 0.001231 | 0.00053219 | 0.00016052 | 0.00039738 | 0.00015627 | 0.0001887 | 0.0003907 | 0.0006706 | 0.00014298 | 0.00057373 | k__Bacteria | p__Firmicutes | c__Bacilli | o__Lactobacillales | f__Carnobacteriace | g__Carnobacterium | 6.57E-03 |
| 5482 | 0.00282277 | 0.00045204 | 0.00035833 | 0.00099522 | 0.00026568 | 0.00017167 | 0.00025832 | 8.33E-05 | 0.00028949 | 0.00033819 | 0.00015001 | 0.00024833 | k__Bacteria | p__Firmicutes | c__Bacilli | o__Bacillales | f__Listeriaceae | g__Brochothrix | 6.43E-03 |
| 444195 | 0.00030984 | 0.00116908 | 0.00030518 | 0.0006859 | 8.86E-05 | 0.00037195 | 0.00047838 | 0.00041905 | 0.00034617 | 0.00147416 | 0.00036799 | 0.0003391 | k__Bacteria | p__Proteobac | c__Alphaprote | o__Rickettsiales | f__mitochondria | g__Zea | 6.36E-03 |
| 446598 | 0.0004872 | 0.00028993 | 0.00046805 | 0.00066284 | 0.00065867 | 0.00043871 | 0.00082918 | 0.00051708 | 0.000498 | 0.0004567 | 0.00033752 | 0.00065423 | k__Bacteria | p__Actinobact | c__Actinobact | o__Actinomycetales | f__Microbacteriacea | g__ | 6.30E-03 |
| 9204 | 0.00025215 | 0.00057986 | 0.00024003 | 0.00066861 | 0.00040406 | 0.00068032 | 0.00044011 | 0.00086507 | 0.00049597 | 0.00033819 | 0.00063753 | 0.00028601 | k__Bacteria | p__Firmicutes | c__Clostridia | o__Clostridiales | f__Veillonellaceae | g__Dialister | 5.89E-03 |
| 21753 | 0.00016667 | 0.00093526 | 0.00061207 | 0.00035159 | 6.64E-05 | 0.0006231 | 0.00044329 | 0.00065186 | 0.00030163 | 0.00038155 | 0.00076644 | 0.00050352 | k__Bacteria | p__Actinobact | c__Thermoleo | o__Solirubrobacterales | f__ | g__ | 5.80E-03 |
| 7729 | 0.0004231 | 0.00054245 | 0.00050749 | 0.00044189 | 7.20E-05 | 0.00042282 | 0.00047838 | 0.00030633 | 0.00041297 | 0.00093363 | 0.00037736 | 0.00078781 | k__Bacteria | p__Proteobac | c__Betaprotec | o__ | f__ | g__ | 5.71E-03 |
| 595433 | 0.00019873 | 0.00028681 | 0.00026746 | 0.00048032 | 0.00017159 | 0.00037195 | 0.00082281 | 0.00085526 | 0.00038463 | 0.000792 | 0.00040783 | 0.00054291 | k__Bacteria | p__Proteobac | c__Epsilonpro | o__Campylobacterales | f__Campylobacterac | g__Campylobacter | 5.58E-03 |
| 610684 | 0.00045728 | 0.00040216 | 0.00029489 | 0.00063979 | 0.00011624 | 0.00046096 | 0.00033805 | 0.00036759 | 0.0007389 | 0.00030928 | 0.00078285 | 0.00059257 | k__Bacteria | p__Proteobac | c__Alphaprote | o__Sphingomonadales | f__Sphingomonada | g__Sphingomonas | 5.50E-03 |
| 426696 | 0.00039318 | 0.00019329 | 0.00053663 | 0.000415 | 0.00021033 | 0.00046732 | 0.00100459 | 0.00034309 | 0.00049395 | 0.00061568 | 0.00023673 | 0.00058572 | k__Bacteria | p__Actinobact | c__Actinobact | o__Actinomycetales | f__Actinomycetacea | g__N09 | 5.50E-03 |
| 27858 | 0.00106628 | 0.00052686 | 0.00014916 | 0.00045726 | 0.0003155 | 0.00022571 | 0.00033167 | 0.00033328 | 0.00042107 | 0.00063591 | 0.0003047 | 0.00067307 | k__Bacteria | p__Proteobac | c__Gammapro | o__Xanthomonadales | f__Xanthomonadac | g__Stenotrophomonas | 5.44E-03 |

| | | | | | | | | | | | | | | | | | | | |
|---|---|---|---|---|---|---|---|---|---|---|---|---|---|---|---|---|---|---|---|
| 14252 | 0.00010684 | 0.0003741 | 0.00028117 | 0.00046879 | 0.00048155 | 0.00024479 | 0.0009727 | 0.00062981 | 0.00044739 | 0.00041623 | 0.00037736 | 0.00061141 | k__Bacteria | p__Firmicutes | c__Clostridia | o__Clostridiales | f__Veillonellaceae | g__Megasphaera | 5.41E-03 |
| 110225 | 8.76E-05 | 0.0002837 | 0.0007698 | 0.00017868 | 0.0001107 | 0.0006231 | 0.0003827 | 0.00073518 | 0.00025102 | 0.00080934 | 0.0007266 | 0.00035794 | k__Bacteria | p__Actinobact | c__Rubrobact | o__Rubrobacterales | f__Rubrobacteracea | g__Rubrobacter | 5.32E-03 |
| 420579 | 0.00019873 | 0.00015276 | 0.00071322 | 0.00034391 | 0.00016052 | 0.0005468 | 0.000472 | 0.00065186 | 0.00054051 | 0.00044803 | 0.00040549 | 0.00062683 | k__Bacteria | p__Actinobact | c__Actinobact | o__Actinomycetales | f__Williamsiaceae | g__Williamsia | 5.26E-03 |
| 609632 | 0.00032053 | 0.0003554 | 0.00068408 | 0.00062442 | 0.00024354 | 0.00022253 | 0.00032848 | 0.00091408 | 0.00041095 | 0.00014742 | 0.00052971 | 0.00047269 | k__Bacteria | p__Firmicutes | c__Clostridia | o__Clostridiales | f__Lachnospiraceae | g__Coprococcus | 5.25E-03 |
| 611582 | 0.00015385 | 0.00039281 | 0.00051434 | 0.00064939 | 0.00021587 | 0.00048322 | 0.00036357 | 0.00045581 | 0.00046358 | 0.0002428 | 0.0006891 | 0.00057031 | k__Bacteria | p__Proteobac | c__Alphaprote | o__Caulobacterales | f__Caulobacteracea | g__Phenylobacterium | 5.19E-03 |
| 611480 | 0.00045087 | 0.00109114 | 0.00050577 | 0.00035544 | 0.00019926 | 0.00040692 | 0.0005294 | 0.00037249 | 0.0002409 | 0.00026882 | 0.00038908 | 0.0003254 | k__Bacteria | p__Proteobac | c__Alphaprote | o__Rhizobiales | f__Bradyrhizobiacea | g__ | 5.14E-03 |
| 435636 | 0.00010684 | 0.00153383 | 0.00043205 | 0.0002978 | 9.96E-05 | 0.00043871 | 0.00028065 | 0.00058325 | 0.00019232 | 0.00013007 | 0.00051799 | 0.00036479 | k__Bacteria | p__Actinobact | c__Actinobact | o__Actinomycetales | f__Micromonospora | g__ | 4.98E-03 |
| 769 | 0.00029488 | 0.00049257 | 0.00073551 | 0.00021518 | 4.98E-05 | 4.45E-05 | 4.46E-05 | 0.0006053 | 0.00189886 | 0.0001214 | 0.00014766 | 0.00023634 | k__Bacteria | p__Proteobac | c__Gammapro | o__Oceanospirillales | f__Oceanospirillace | g__Marinomonas | 4.89E-03 |
| 12211 | 0.00047011 | 0.00050816 | 0.00048348 | 0.00070895 | 0.00019926 | 0.00034334 | 0.00035081 | 0.00045826 | 0.00028746 | 0.00023991 | 0.00027189 | 0.0004607 | k__Bacteria | p__Firmicutes | c__Bacilli | o__Bacillales | f__Planococcaceae | g__Staphylococcus | 4.78E-03 |
| 15639 | 0.00035044 | 0.00032422 | 0.00020917 | 0.00038618 | 0.00028229 | 0.00033698 | 0.00044967 | 0.00043866 | 0.00076116 | 0.00052029 | 0.00034689 | 0.00034938 | k__Bacteria | p__Proteobac | c__Gammapro | o__Pasteurellales | f__Pasteurellaceae | g__ | 4.76E-03 |
| 397576 | 0.00014531 | 0.00052998 | 0.0005452 | 0.00031509 | 0.00019373 | 0.00036877 | 0.00055492 | 0.00071558 | 0.00034617 | 0.0001214 | 0.00055315 | 0.00035794 | k__Bacteria | p__Actinobact | c__Actinobact | o__Actinomycetales | f__Beutenbergiacea | g__Beutenbergia | 4.75E-03 |
| 611811 | 0.00018163 | 0.00062663 | 0.00038061 | 0.00025553 | 0.00011624 | 0.00049593 | 0.00054535 | 0.0006053 | 0.00020649 | 0.00026304 | 0.00050393 | 0.00055832 | k__Bacteria | p__Proteobac | c__Alphaprote | o__Rhizobiales | f__Bradyrhizobiacea | g__Balneimonas | 4.74E-03 |
| 616224 | 0.00057908 | 0.00046763 | 0.00059321 | 0.00026898 | 0.0003321 | 0.00033062 | 0.0005294 | 0.00032348 | 0.00051824 | 0.00027749 | 0.00025782 | 0.00021922 | k__Bacteria | p__Firmicutes | c__Clostridia | o__Clostridiales | f__[Tissierellaceae] | g__Parvimonas | 4.70E-03 |
| 612991 | 0.00012821 | 0.00023382 | 0.00024174 | 0.00024977 | 0.00012731 | 0.00023843 | 0.00140642 | 0.00051708 | 0.00030568 | 0.0005492 | 0.00037971 | 0.00020894 | k__Bacteria | p__Firmicutes | c__Clostridia | o__Clostridiales | f__Lachnospiraceae | g__Moryella | 4.59E-03 |
| 428135 | 0.00035258 | 0.00024629 | 0.00051091 | 0.00032278 | 0.00014945 | 0.00056905 | 0.00059638 | 0.00057344 | 0.00019636 | 0.00033241 | 0.00016641 | 0.00056688 | k__Bacteria | p__Actinobact | c__Actinobact | o__Actinomycetales | f__Beutenbergiacea | g__Salana | 4.58E-03 |
| 10161 | 0.00091671 | 2.49E-05 | 0.00015087 | 0.00041692 | 0.00024908 | 8.27E-05 | 0.00034762 | 0.00037249 | 0.00085631 | 0.00017343 | 0.00053206 | 0.0004333 | k__Bacteria | p__Proteobac | c__Gammapro | o__Alteromonadales | f__Shewanellaceae | g__ | 4.56E-03 |
| 608894 | 0.00109834 | 2.49E-05 | 0.00018688 | 0.00011528 | 0.00015498 | 7.63E-05 | 2.87E-05 | 7.84E-05 | 0.00014575 | 0.00253787 | 7.03E-06 | 9.93E-05 | k__Bacteria | p__Proteobac | c__Alphaprote | o__Rhodospirillales | f__Acetobacteracea | g__Acetobacter | 4.55E-03 |
| 609876 | 0.00106201 | 0.00093526 | 0.00048863 | 0.00026322 | 0.00032657 | 0.00030519 | 7.02E-05 | 0.00033573 | 0.00014778 | 0.0001214 | 0.00025782 | 0.00023634 | k__Bacteria | p__Firmicutes | c__Clostridia | o__Clostridiales | f__Lachnospiraceae | g__[Ruminococcus] | 4.55E-03 |
| 428344 | 0.00055772 | 0.00057674 | 0.0004972 | 0.00031701 | 0.00022694 | 0.00023207 | 0.00046562 | 0.00047297 | 0.000332 | 0.00030061 | 0.00021798 | 0.00032711 | k__Bacteria | p__Actinobact | c__Actinobact | o__Actinomycetales | f__Microbacteriacea | g__Microbacterium | 4.52E-03 |
| 420377 | 0.00028634 | 0.00034605 | 0.00028289 | 0.00083384 | 0.00014391 | 0.00028929 | 0.00029021 | 0.00028182 | 0.00028746 | 0.00098855 | 0.00019923 | 0.00020038 | k__Bacteria | p__Proteobac | c__Alphaprote | o__Rickettsiales | f__mitochondria | g__Nelumbo | 4.43E-03 |
| 609983 | 4.27E-05 | 0.000106 | 0.00020574 | 0.00048608 | 0.00021033 | 0.00078205 | 8.29E-05 | 0.000397 | 0.00150411 | 0.00016476 | 0.00015938 | 0.00019695 | k__Bacteria | p__Proteobac | c__Alphaprote | o__Rhizobiales | f__Methylocystacea | g__ | 4.34E-03 |
| 586417 | 0.00020086 | 0.00027434 | 0.00080409 | 0.00039578 | 0.00011624 | 0.00029247 | 0.0003827 | 0.00054159 | 0.00032592 | 0.00032663 | 0.00030236 | 0.00031855 | k__Bacteria | p__Actinobact | c__Actinobact | o__Actinomycetales | f__Nocardioidaceae | g__Propionicimonas | 4.28E-03 |
| 449468 | 0.00035685 | 0.00013717 | 0.0010184 | 0.00021134 | 0.00062546 | 0.00024161 | 0.00016584 | 0.00017399 | 0.00019434 | 0.00014453 | 0.00017579 | 0.0007587 | k__Bacteria | p__Actinobact | c__Actinobact | o__Actinomycetales | f__Dermabacterace | g__Dermabacter | 4.20E-03 |
| 448725 | 0.00032907 | 0.00042399 | 0.00051777 | 0.00013833 | 0.00022694 | 0.00034334 | 0.00046881 | 0.00037004 | 0.00024292 | 0.00020523 | 0.0004219 | 0.00049324 | k__Bacteria | p__Actinobact | c__Actinobact | o__Actinomycetales | f__Sanguibacterac | g__Sanguibacter | 4.18E-03 |
| 25268 | 0.00086115 | 0.00019017 | 0.0001543 | 0.00010567 | 3.87E-05 | 8.90E-05 | 0.00150848 | 0.00012008 | 6.88E-05 | 0.00012429 | 0.00059769 | 0.00030314 | k__Bacteria | p__Firmicutes | c__Bacilli | o__Turicibacterales | f__Turicibacteracea | g__Turicibacter | 4.16E-03 |
| 434933 | 0.00023292 | 0.00049569 | 0.0003669 | 0.00022863 | 0.00012731 | 0.00040692 | 0.00059319 | 0.00036269 | 0.00021053 | 0.00017343 | 0.00061175 | 0.00034424 | k__Bacteria | p__Actinobact | c__Actinobact | o__Actinomycetales | f__Cellulomonadace | g__Actinotalea | 4.15E-03 |
| 583880 | 0.00010471 | 0.00052686 | 0.00054692 | 0.00035928 | 0.00023247 | 9.86E-05 | 0.00037313 | 0.00049747 | 0.00022066 | 0.0002428 | 0.00045002 | 0.00041446 | k__Bacteria | p__Firmicutes | c__Clostridia | o__Clostridiales | f__Ruminococcacea | g__Ruminococcus | 4.07E-03 |
| 610457 | 0.00147015 | 0.00023693 | 0.0002846 | 0.00026706 | 5.54E-05 | 0.00026386 | 0.00010843 | 0.00024506 | 0.00026722 | 0.00017343 | 0.00037736 | 0.00027573 | k__Bacteria | p__Proteobac | c__Alphaprote | o__Rhodobacterales | f__Rhodobacterace | g__Amaricoccus | 4.03E-03 |
| 610907 | 0.00029916 | 0.00038346 | 0.0003789 | 0.00036696 | 0.00016052 | 0.00038785 | 0.00031573 | 0.00033818 | 0.00018624 | 0.0002746 | 0.00044533 | 0.00038192 | k__Bacteria | p__Proteobac | c__Alphaprote | o__Rhizobiales | f__Hyphomicrobiace | g__Devosia | 3.92E-03 |
| 595047 | 7.91E-05 | 0.00118466 | 0.00058807 | 0.00028435 | 9.41E-05 | 0.00026068 | 4.15E-05 | 0.00062491 | 0.00038868 | 7.80E-05 | 3.75E-05 | 0.00024662 | k__Bacteria | p__Firmicutes | c__Clostridia | o__Clostridiales | f__Lachnospiraceae | g__Roseburia | 3.91E-03 |
| 9657 | 0.00123082 | 0.00040528 | 0.00021431 | 9.80E-05 | 8.30E-05 | 0.00022889 | 0.00023919 | 0.00036269 | 0.00017005 | 0.00015609 | 0.00045705 | 0.00022093 | k__Bacteria | p__Firmicutes | c__Bacilli | o__Bacillales | f__Paenibacillaceae | g__ | 3.87E-03 |
| 363834 | 0.0002842 | 0.00072639 | 0.00027775 | 0.00030933 | 0.00021033 | 0.0001526 | 0.00022962 | 0.00029407 | 0.00036439 | 0.00040467 | 0.00037736 | 0.00022949 | k__Bacteria | p__Bacteroide | c__Flavobacte | o__Flavobacteriales | f__[Weeksellaceae] | g__ | 3.86E-03 |
| 420245 | 0.00013889 | 0.00010288 | 0.00026574 | 0.00021518 | 0.00019926 | 0.00018121 | 0.00195815 | 0.00016909 | 0.00019029 | 9.83E-05 | 0.00010079 | 0.00021751 | k__Bacteria | p__Actinobact | c__Actinobact | o__Actinomycetales | f__Yaniellaceae | g__Yaniella | 3.84E-03 |
| 427270 | 5.13E-05 | 0.0004271 | 0.00011144 | 9.41E-05 | 2.77E-05 | 5.72E-05 | 0.00255134 | 0.00011028 | 7.09E-05 | 2.89E-05 | 7.73E-05 | 0.00022607 | k__Bacteria | p__Actinobact | c__Actinobact | o__Actinomycetales | f__Promicromonosp | g__Xylanimicrobium | 3.83E-03 |
| 375356 | 0.00010043 | 0.00073574 | 0.00053492 | 0.00018636 | 3.87E-05 | 0.00037195 | 0.00022962 | 0.00033573 | 0.00019232 | 0.00013874 | 0.00052034 | 0.00041617 | k__Bacteria | p__Bacteroide | c__Cytophagia | o__Cytophagales | f__Cytophagaceae | g__Adhaeribacter | 3.80E-03 |
| 448932 | 0.00065815 | 8.73E-05 | 0.0008058 | 0.00016907 | 0.00018266 | 0.0004101 | 0.00028065 | 0.00023281 | 0.00025305 | 0.00028905 | 7.97E-05 | 0.00030656 | k__Bacteria | p__Actinobact | c__Actinobact | o__Actinomycetales | f__Propionibacteria | g__Luteococcus | 3.75E-03 |
| 611331 | 0.00075431 | 0.00021511 | 0.0002486 | 0.00019789 | 7.20E-05 | 0.00039102 | 0.00055811 | 0.0002181 | 0.0001579 | 0.00066771 | 0.00011485 | 0.00015756 | k__Bacteria | p__Firmicutes | c__Clostridia | o__Clostridiales | f__Lachnospiraceae | g__Oribacterium | 3.75E-03 |
| 396712 | 5.77E-05 | 0.00029305 | 0.00066522 | 0.00015947 | 8.86E-05 | 0.00030201 | 0.00020411 | 0.0003823 | 0.00021053 | 0.00026882 | 0.0006891 | 0.00029115 | k__Bacteria | p__Actinobact | c__Actinobact | o__Actinomycetales | f__Pseudonocardia | g__Actinomycetospora | 3.61E-03 |
| 624015 | 0.00014103 | 0.00011847 | 0.0002846 | 0.0002978 | 0.00012177 | 0.00046096 | 0.00016903 | 0.00076214 | 0.00014575 | 0.00031507 | 0.00040314 | 0.00038877 | k__Bacteria | p__[Thermi] | c__Deinococci | o__Deinococcales | f__Trueperaceae | g__Truepera | 3.61E-03 |
| 18949 | 0.00029061 | 0.00064533 | 0.00016288 | 0.00020558 | 9.96E-05 | 0.00019074 | 0.00020411 | 0.00047542 | 0.00018624 | 0.00016765 | 0.00059534 | 0.0002843 | k__Bacteria | p__Proteobac | c__Gammapro | o__Xanthomonadales | f__Xanthomonadac | g__Lysobacter | 3.51E-03 |
| 589958 | 0.00012821 | 0.00032734 | 0.00039776 | 0.00022479 | 0.00019926 | 0.00029565 | 3.83E-05 | 0.00028917 | 0.00018827 | 0.00095676 | 0.00012422 | 0.00033739 | k__Bacteria | p__TM7 | c__TM7-3 | o__CW040 | f__ | g__ | 3.51E-03 |
| 597358 | 7.05E-05 | 0.00026499 | 0.00026574 | 0.00028819 | 4.98E-05 | 0.00042917 | 0.00022643 | 0.00065922 | 0.00034414 | 0.00016765 | 0.00042893 | 0.0003117 | k__Bacteria | p__Actinobact | c__Acidimicrob | o__Acidimicrobiales | f__C111 | g__ | 3.51E-03 |
| 448617 | 0.00036754 | 0.00019329 | 0.00053835 | 0.0001806 | 0.00052583 | 0.00014306 | 0.000472 | 7.60E-05 | 0.00023483 | 0.00033241 | 0.00014532 | 0.00025176 | k__Bacteria | p__Actinobact | c__Actinobact | o__Actinomycetales | f__Microbacteriacea | g__Leucobacter | 3.46E-03 |
| 426514 | 0.00035685 | 0.00011535 | 0.00056235 | 0.00022479 | 7.20E-05 | 0.00020982 | 0.00053578 | 0.00034309 | 0.00031783 | 0.00020523 | 0.00016876 | 0.00027916 | k__Bacteria | p__Actinobact | c__Actinobact | o__Actinomycetales | f__Dermatophilacea | g__Kineosphaera | 3.39E-03 |
| 72400 | 0.00020941 | 0 | 0.00013716 | 0.00034775 | 0.00014391 | 0.00033698 | 0.0005294 | 0.00010293 | 0.00035629 | 0.00028616 | 0.00077816 | 0.00013872 | k__Bacteria | p__Spirochaet | c__Spirochaet | o__Spirochaetales | f__Spirochaetaceae | g__Treponema | 3.37E-03 |
| 20050 | 0.00030984 | 0.00038969 | 0.00020574 | 0.00039194 | 0.00012177 | 0.00014942 | 0.0002647 | 0.0002132 | 0.00021661 | 0.00054052 | 0.00024376 | 0.00029971 | k__Bacteria | p__Proteobac | c__Betaprotec | o__Burkholderiales | f__Oxalobacteracea | g__Janthinobacterium | 3.35E-03 |
| 109653 | 0.00011753 | 0.00074197 | 0.0002126 | 0.00012296 | 0.00012731 | 0.00046096 | 0.0001467 | 0.00031368 | 0.00020041 | 0.00022835 | 0.00030236 | 0.00030314 | k__Bacteria | p__Actinobact | c__Thermoleo | o__Gaiellales | f__Gaiellaceae | g__ | 3.28E-03 |
| 431939 | 5.13E-05 | 0.00013405 | 9.77E-05 | 0.00018829 | 0.0002048 | 0.00011445 | 0.00016265 | 0.00186737 | 7.49E-05 | 7.52E-05 | 0.00018048 | 9.42E-05 | k__Bacteria | p__Actinobact | c__Actinobact | o__Actinomycetales | f__Intrasporangiace | g__Janibacter | 3.25E-03 |
| 610468 | 2.99E-05 | 9.04E-05 | 0.00037033 | 2.50E-05 | 0 | 8.58E-05 | 0.00203788 | 0.00015439 | 7.09E-05 | 0.00014453 | 6.80E-05 | 0.00014729 | k__Bacteria | p__Firmicutes | c__Clostridia | o__Clostridiales | f__Clostridiaceae | g__Sarcina | 3.22E-03 |
| 8945 | 0.00015599 | 0.00014652 | 0.00055721 | 0.00017868 | 4.98E-05 | 0.0004101 | 0.00040821 | 0.00035779 | 0.00031378 | 0.00020234 | 0.00027189 | 8.73E-05 | k__Bacteria | p__Proteobac | c__Gammapro | o__Pasteurellales | f__Pasteurellaceae | g__Actinobacillus | 3.14E-03 |
| 19641 | 0.00016667 | 0.00147148 | 0.00010458 | 0.00029396 | 5.54E-05 | 8.27E-05 | 0.00011162 | 0.00011028 | 0.00019434 | 8.09E-05 | 0.00032814 | 0.00013187 | k__Bacteria | p__Proteobac | c__Gammapro | o__Alteromonadales | f__[Chromatiaceae] | g__ | 3.13E-03 |
| 24681 | 0.00019445 | 0.00026811 | 0.00014573 | 0.00023055 | 0.00089114 | 0.00010173 | 0.00021686 | 0.00012498 | 0.00020041 | 0.00011851 | 0.00032111 | 0.00028944 | k__Bacteria | p__Firmicutes | c__Bacilli | o__Bacillales | f__ | g__ | 3.10E-03 |
| 610022 | 8.12E-05 | 0.00072639 | 0.00051949 | 0.00025937 | 3.32E-05 | 0.00028929 | 0.00021686 | 0.00020095 | 5.67E-05 | 0.00013874 | 0.00031408 | 0.00026032 | k__Bacteria | p__Chloroflexi | c__Ellin6529 | o__ | f__ | g__ | 3.10E-03 |
| 612115 | 0.00033548 | 0.0001777 | 0.00011487 | 0.00032662 | 0.00011624 | 0.00075662 | 0.00022324 | 0.00016419 | 0.00015385 | 0.00015031 | 0.0003258 | 0.00023121 | k__Bacteria | p__Proteobac | c__Alphaprote | o__Sphingomonadales | f__Erythrobacterac | g__ | 3.08E-03 |
| 377715 | 8.33E-05 | 0.00033981 | 0.00037547 | 0.00013833 | 0.00018266 | 0.00017485 | 0.00025513 | 0.00024506 | 0.00022471 | 0.00036131 | 0.00050159 | 0.00013872 | k__Bacteria | p__Firmicutes | c__Erysipelotri | o__Erysipelotrichales | f__Erysipelotrichace | g__Bulleidia | 3.02E-03 |
| 14443 | 8.76E-05 | 0.0005861 | 0.00012859 | 0.00017292 | 9.96E-05 | 0.0003497 | 0.00036357 | 0.00031858 | 9.92E-05 | 0.00023413 | 0.00035861 | 0.00022093 | k__Bacteria | p__Actinobact | c__Thermoleo | o__Solirubrobacterales | f__Solirubrobactera | g__ | 3.02E-03 |
| 359872 | 0.0001624 | 0.00021823 | 0.00029489 | 0.00033238 | 0.00029889 | 0.00017167 | 7.02E-05 | 0.00029162 | 0.00016802 | 7.23E-05 | 0.00065394 | 0.0002843 | k__Bacteria | p__Bacteroide | c__Bacteroidia | o__Bacteroidales | f__Bacteroidaceae | g__Bacteroides | 3.02E-03 |
| 612673 | 4.06E-05 | 0.00025876 | 0.00040976 | 0.00026898 | 4.43E-05 | 0.00024161 | 0.00029978 | 0.00021075 | 0.00038666 | 0.0001821 | 0.00037033 | 0.00028944 | k__Bacteria | p__Proteobac | c__Alphaprote | o__Rhodospirillales | f__Rhodospirillacea | g__ | 3.00E-03 |
| 410759 | 8.55E-05 | 0.00023693 | 0.00062921 | 0.00015947 | 4.43E-05 | 0.00026068 | 0.00036357 | 0.00046317 | 0.00017005 | 0.00017343 | 6.33E-05 | 0.00032883 | k__Bacteria | p__Actinobact | c__Actinobact | o__Actinomycetales | f__Kineosporiaceae | g__Kineococcus | 2.98E-03 |
| 610126 | 9.83E-05 | 0.00033046 | 0.00073551 | 0.00019021 | 2.21E-05 | 0.00020028 | 0.00026789 | 0.00045091 | 5.87E-05 | 8.09E-05 | 0.00034689 | 0.00019182 | k__Bacteria | p__Acidobacte | c__[Chloracido | o__RB41 | f__Ellin6075 | g__ | 2.97E-03 |
| 365025 | 0.00096372 | 5.61E-05 | 0.00018345 | 9.22E-05 | 0.00065867 | 8.58E-05 | 0.00017222 | 0.00026712 | 4.86E-05 | 0.00027749 | 5.39E-05 | 0.00011132 | k__Bacteria | p__Bacteroide | c__Flavobacte | o__Flavobacteriales | f__Flavobacteriacea | g__Myroides | 2.97E-03 |
| 435200 | 0.00015172 | 0.00029617 | 0.00022803 | 0.00013449 | 9.96E-05 | 0.00040692 | 0.00043054 | 0.00023036 | 0.00020446 | 0.00020234 | 0.00028595 | 0.00028772 | k__Bacteria | p__Actinobact | c__Actinobact | o__Actinomycetales | f__Propionibacteria | g__ | 2.96E-03 |
| 412648 | 3.21E-05 | 0.00031175 | 0.00074065 | 0.0001806 | 6.09E-05 | 0.00037831 | 0.0002073 | 0.000397 | 0.00012349 | 0.00021101 | 0.00013126 | 0.00014215 | k__Bacteria | p__Actinobact | c__Actinobact | o__Actinomycetales | f__Kineosporiaceae | g__ | 2.92E-03 |
| 587246 | 8.33E-05 | 0.00018705 | 0.00036861 | 0.00023824 | 9.41E-05 | 0.0001971 | 0.00044011 | 8.82E-05 | 0.00064578 | 0.00015609 | 0.00022267 | 0.00019182 | k__Bacteria | p__Firmicutes | c__Clostridia | o__Clostridiales | f__Peptostreptococ | g__Filifactor | 2.91E-03 |
| 610982 | 2.99E-05 | 1.56E-05 | 0.00041662 | 0.00012296 | 0.00019373 | 7.31E-05 | 0.00056448 | 0.00023526 | 0.00020446 | 0.00035553 | 0.00052268 | 0.00017469 | k__Bacteria | p__Firmicutes | c__Clostridia | o__Clostridiales | f__Lachnospiraceae | g__Shuttleworthia | 2.91E-03 |
| 614508 | 0.00015813 | 0.00038346 | 0.00056921 | 0.00025553 | 5.54E-05 | 0.00030201 | 0.00021367 | 0.00016909 | 0.00015993 | 0.00023702 | 0.00020626 | 0.0001627 | k__Bacteria | p__Proteobac | c__Alphaprote | o__Rhizobiales | f__Methylobacteriac | g__ | 2.87E-03 |
| 365019 | 0.0002436 | 0.00032422 | 0.0003429 | 0.000244 | 2.21E-05 | 0.0001367 | 0.00051984 | 0.00022546 | 0.00017814 | 0.00015609 | 0.00020626 | 0.00026717 | k__Bacteria | p__Bacteroide | c__Sphingoba | o__Sphingobacteriales | f__Sphingobacteria | g__Sphingobacterium | 2.87E-03 |
| 363774 | 0.00030771 | 0.00027434 | 0.00024174 | 0.00030164 | 3.32E-05 | 0.00024479 | 0.00022324 | 0.00028672 | 0.00035022 | 0.00015898 | 0.00016407 | 0.00026375 | k__Bacteria | p__Bacteroide | c__Sphingoba | o__Sphingobacteriales | f__Sphingobacteria | g__Pedobacter | 2.85E-03 |
| 442773 | 1.07E-05 | 6.86E-05 | 5.83E-05 | 1.34E-05 | 3.32E-05 | 1.59E-05 | 0.0022388 | 2.21E-05 | 0.00015993 | 5.49E-05 | 0.00010547 | 5.14E-05 | k__Bacteria | p__Actinobact | c__Actinobact | o__Actinomycetales | f__Pseudonocardia | g__Saccharomonospo | 2.83E-03 |
| 113020 | 8.55E-05 | 0.00048945 | 0.00025546 | 0.00021518 | 8.86E-05 | 0.00011763 | 0.00067929 | 4.90E-05 | 0.00021661 | 0.00012718 | 0.0002883 | 0.00021922 | k__Bacteria | p__Firmicutes | c__Bacilli | o__Bacillales | f__Bacillaceae | g__ | 2.83E-03 |
| 19866 | 5.13E-05 | 0.00054245 | 0.00087267 | 9.61E-05 | 9.96E-05 | 0.00011445 | 0.00034443 | 0.00012253 | 0.00011944 | 8.38E-05 | 0.0002086 | 0.00015585 | k__Bacteria | p__Firmicutes | c__Bacilli | o__Bacillales | f__Planococcaceae | g__Sporosarcina | 2.81E-03 |
| 38948 | 0.00018804 | 0.00017146 | 9.94E-05 | 0.00024977 | 0.00012177 | 7.63E-05 | 0.00018497 | 0.00014949 | 0.0002571 | 0.00018499 | 0.00013594 | 0.00094538 | k__Bacteria | p__Proteobac | c__Gammapro | o__Enterobacteriales | f__Enterobacteriace | g__Citrobacter | 2.76E-03 |
| 53099 | 0.00010898 | 2.18E-05 | 4.29E-05 | 0.00013833 | 0.00012177 | 0.00017803 | 0.00164242 | 8.33E-05 | 0.00011741 | 0.00011273 | 7.03E-05 | 7.19E-05 | k__Bacteria | p__Firmicutes | c__Bacilli | o__Gemellales | f__Gemellaceae | g__Gemella | 2.71E-03 |
| 617413 | 0.00013035 | 0.00025564 | 0.00017316 | 0.00027859 | 0.00032103 | 0.00034652 | 0.00013395 | 0.00013723 | 0.00016397 | 0.00014163 | 0.00020392 | 0.00035623 | k__Bacteria | p__Proteobac | c__Alphaprote | o__Rhizobiales | f__ | g__ | 2.64E-03 |
| 611161 | 7.91E-05 | 0.00011223 | 0.00021774 | 0.0004246 | 0.00014945 | 0.00011763 | 0.00041459 | 0.00016174 | 0.00010324 | 0.00027171 | 0.00043361 | 0.00014215 | k__Bacteria | p__Proteobac | c__Alphaprote | o__Caulobacterales | f__Caulobacteracea | g__Brevundimonas | 2.63E-03 |
| 451281 | 6.84E-05 | 0.0004271 | 0.00033775 | 0.00014217 | 0.00011624 | 0.00023525 | 0.00035719 | 0.00021565 | 8.70E-05 | 0.00010406 | 0.00023439 | 0.00028601 | k__Bacteria | p__Actinobact | c__Actinobact | o__Actinomycetales | f__Streptomycetace | g__ | 2.61E-03 |
| 363483 | 0.0005043 | 0.00064533 | 0.00015773 | 0.00013065 | 4.98E-05 | 0.00019392 | 0.0001467 | 0.00012253 | 0.00030163 | 9.83E-05 | 0.00012422 | 0.00011817 | k__Bacteria | p__Bacteroide | c__Flavobacte | o__Flavobacteriales | f__[Weeksellaceae] | g__Cloacibacterium | 2.59E-03 |
| 612474 | 0.00013889 | 9.35E-05 | 0.00022117 | 0.0001537 | 0.00012177 | 0.00024797 | 0.00031573 | 0.00020585 | 0.0002328 | 0.00049428 | 0.00015704 | 0.00015756 | k__Bacteria | p__Firmicutes | c__Clostridia | o__Clostridiales | f__Lachnospiraceae | g__Catonella | 2.54E-03 |
| 610781 | 9.83E-05 | 0.00020576 | 8.74E-05 | 0.00014794 | 0.00029889 | 0.00033062 | 0.00013395 | 0.00040925 | 0.00021661 | 0.0001214 | 0.00019688 | 0.00027231 | k__Bacteria | p__Firmicutes | c__Clostridia | o__Clostridiales | f__[Tissierellaceae] | g__1-68 | 2.52E-03 |

| | | | | | | | | | | | | | | | | | | | |
|---|---|---|---|---|---|---|---|---|---|---|---|---|---|---|---|---|---|---|---|
| 610936 | 6.84E-05 | 0.00061727 | 0.00033947 | 0.00021518 | 1.66E-05 | 6.68E-05 | 0.00014351 | 0.00018625 | 0.00014373 | 0.00013296 | 0.00034924 | 0.00022436 | k__Bacteria | p__Proteobac | c__Epsilonpro | o__Campylobacterales | f__Campylobacterac | g__Arcobacter | 2.50E-03 |
| 416337 | 4.27E-06 | 0.000106 | 0.00029318 | 0.00021326 | 0.00050922 | 0.00016213 | 3.83E-05 | 9.56E-05 | 2.83E-05 | 5.78E-05 | 0.00073363 | 0.00024491 | k__Bacteria | p__Actinobact | c__Actinobact | o__Actinomycetales | f__Actinomycetacea | g__Actinobaculum | 2.49E-03 |
| 364240 | 5.98E-05 | 0.00023693 | 0.00024688 | 0.00021903 | 6.64E-05 | 0.00029883 | 0.00038589 | 0.00021565 | 0.00014778 | 0.00010695 | 0.00033752 | 0.00013187 | k__Bacteria | p__Bacteroide | c__Sphingoba | o__Sphingobacteriales | f__Sphingobacteria | g__ | 2.45E-03 |
| 586656 | 0.00013248 | 0.00014029 | 0.00023317 | 0.00017292 | 0.00015498 | 0.00013034 | 0.00040502 | 0.00014704 | 0.00019636 | 0.0003035 | 0.00027892 | 0.00015585 | k__Bacteria | p__Firmicutes | c__Clostridia | o__Clostridiales | f__[Mogibacteriacea | g__Mogibacterium | 2.45E-03 |
| 23690 | 0.00050643 | 7.48E-05 | 8.40E-05 | 0.00018829 | 0.00029889 | 2.54E-05 | 0.00072075 | 6.37E-05 | 7.09E-05 | 8.38E-05 | 0.00010547 | 0.00017469 | k__Bacteria | p__Firmicutes | c__Bacilli | o__Bacillales | f__Planococcaceae | g__Kurthia | 2.40E-03 |
| 406147 | 1.92E-05 | 1.25E-05 | 0.00012859 | 4.80E-05 | 0.00012177 | 0.00045779 | 0.00030297 | 0.00085526 | 0.00022673 | 3.18E-05 | 0.00011485 | 7.02E-05 | k__Bacteria | p__Actinobact | c__Actinobact | o__Bifidobacteriales | f__Bifidobacteriacea | g__Scardovia | 2.39E-03 |
| 610449 | 0.00019659 | 0.00015899 | 8.06E-05 | 0.00010567 | 3.87E-05 | 4.13E-05 | 0.00012438 | 4.66E-05 | 0.00014575 | 0.00014163 | 7.27E-05 | 0.00120741 | k__Bacteria | p__Proteobac | c__Alphaprote | o__Rhizobiales | f__Brucellaceae | g__Ochrobactrum | 2.36E-03 |
| 370945 | 0.00011539 | 0.00033981 | 0.00026232 | 0.00010759 | 3.32E-05 | 0.00015895 | 0.00025513 | 0.00022546 | 0.00019839 | 0.00017054 | 0.00020157 | 0.00028259 | k__Bacteria | p__Bacteroide | c__[Saprospira | o__[Saprospirales] | f__Chitinophagacea | g__Flavisolibacter | 2.35E-03 |
| 51013 | 3.21E-05 | 7.48E-05 | 1.89E-05 | 1.34E-05 | 3.32E-05 | 6.36E-06 | 0.00202512 | 1.72E-05 | 2.43E-05 | 1.16E-05 | 3.05E-05 | 3.77E-05 | k__Bacteria | p__Firmicutes | c__Bacilli | o__Bacillales | f__Planococcaceae | g__Rummeliibacillus | 2.33E-03 |
| 16589 | 0.00013889 | 0.00016835 | 7.20E-05 | 0.00019981 | 0.0001107 | 0.00012716 | 0.00029978 | 0.00015929 | 0.00027329 | 0.00047693 | 9.38E-05 | 0.00019353 | k__Bacteria | p__Proteobac | c__Betaprotec | o__Neisseriales | f__Neisseriaceae | g__Kingella | 2.31E-03 |
| 626785 | 6.20E-05 | 0.00025564 | 0.00036861 | 0.00012296 | 5.54E-06 | 0.0002575 | 0.00022962 | 0.00030143 | 2.02E-05 | 0.00019366 | 0.00014532 | 0.00030656 | k__Bacteria | p__Chloroflexi | c__Chloroflexi | o__AKIW781 | f__ | g__ | 2.27E-03 |
| 609766 | 2.99E-05 | 0.00025876 | 0.00039776 | 0.00024016 | 4.43E-05 | 0.00012398 | 0.00012438 | 0.00016174 | 0.00015183 | 9.83E-05 | 0.00044533 | 9.93E-05 | k__Bacteria | p__Proteobac | c__Alphaprote | o__Rhizobiales | f__Aurantimonadac | g__Aurantimonas | 2.18E-03 |
| 69324 | 0.00011325 | 0.00020888 | 0.0001663 | 0.00029588 | 0.00011624 | 9.86E-05 | 0.00023281 | 0.00023281 | 0.00010324 | 0.00017054 | 0.00018282 | 0.00021237 | k__Bacteria | p__Firmicutes | c__Clostridia | o__Clostridiales | f__Veillonellaceae | g__ | 2.13E-03 |
| 55469 | 0.0001218 | 0.00036787 | 9.26E-05 | 0.000244 | 7.20E-05 | 8.27E-05 | 0.00021367 | 9.56E-05 | 0.00023483 | 0.00011851 | 0.00030939 | 0.00013016 | k__Bacteria | p__Firmicutes | c__Bacilli | o__Bacillales | f__[Exiguobacterac | g__ | 2.08E-03 |
| 449445 | 0.0001624 | 0.0002494 | 0.00016459 | 8.07E-05 | 0.00011624 | 0.00019392 | 0.00031573 | 0.00012253 | 0.00018219 | 8.09E-05 | 0.00016407 | 0.00023977 | k__Bacteria | p__Actinobact | c__Actinobact | o__Actinomycetales | f__Cellulomonadace | g__Sediminihabitans | 2.07E-03 |
| 80183 | 9.83E-05 | 9.04E-05 | 0.00016802 | 0.0002344 | 6.64E-05 | 0.00026704 | 0.00020411 | 0.00015439 | 0.00014373 | 0.00023991 | 0.00024845 | 0.00013872 | k__Bacteria | p__Proteobac | c__Betaprotec | o__Burkholderiales | f__ | g__ | 2.05E-03 |
| 366092 | 0.00084619 | 0.00024629 | 0.00013544 | 0.00010567 | 1.11E-05 | 5.40E-05 | 0.00012119 | 6.13E-05 | 0.00015993 | 6.07E-05 | 0.00011251 | 0.00010276 | k__Bacteria | p__Bacteroide | c__[Saprospira | o__[Saprospirales] | f__Chitinophagacea | g__ | 2.02E-03 |
| 86054 | 0.00013035 | 0.0002837 | 9.43E-05 | 0.00021134 | 0.00012731 | 0.00015577 | 0.00015627 | 6.13E-05 | 0.00023685 | 0.00011851 | 0.00014766 | 0.00029115 | k__Bacteria | p__Proteobac | c__Betaprotec | o__Rhodocyclales | f__Rhodocyclaceae | g__ | 2.01E-03 |
| 375033 | 2.56E-05 | 0.00053933 | 0.00013201 | 9.80E-05 | 4.43E-05 | 4.77E-05 | 0.00011481 | 0.00010783 | 0.00012956 | 9.54E-05 | 0.00048284 | 0.00019353 | k__Bacteria | p__Bacteroide | c__Cytophagia | o__Cytophagales | f__Cytophagaceae | g__Pontibacter | 2.01E-03 |
| 610839 | 0.00020086 | 0.00044269 | 0.00020402 | 0.00014217 | 0.0001107 | 6.68E-05 | 8.93E-05 | 0.00012498 | 0.00011944 | 5.20E-05 | 0.00025079 | 0.00016099 | k__Bacteria | p__Proteobac | c__Alphaprote | o__Rhodobacterales | f__Rhodobacterace | g__Rhodobacter | 1.96E-03 |
| 610912 | 0.00014103 | 0.00026811 | 0.00015259 | 0.00018829 | 0.00013838 | 0.0001367 | 0.00012119 | 0.00028427 | 7.90E-05 | 0.00011851 | 8.91E-05 | 0.00017469 | k__Bacteria | p__Firmicutes | c__Clostridia | o__Clostridiales | f__Lachnospiraceae | g__Dorea | 1.89E-03 |
| 622850 | 4.91E-05 | 0 | 0.00114699 | 2.69E-05 | 6.09E-05 | 0.00010809 | 6.06E-05 | 0.00012253 | 1.62E-05 | 6.36E-05 | 0.00012188 | 0.00011132 | k__Bacteria | p__Firmicutes | c__Clostridia | o__Clostridiales | f__[Tissierellaceae] | g__Gallicola | 1.89E-03 |
| 6067 | 0.00010684 | 9.35E-05 | 6.17E-05 | 0.00055333 | 2.77E-05 | 2.23E-05 | 7.97E-05 | 8.33E-05 | 0.00011336 | 0.00063591 | 3.52E-05 | 6.85E-05 | k__Bacteria | p__Proteobac | c__Gammapro | o__Vibrionales | f__Vibrionaceae | g__Vibrio | 1.88E-03 |
| 417557 | 1.92E-05 | 0.00026811 | 0.00046291 | 6.92E-05 | 0 | 0.000213 | 9.25E-05 | 5.88E-05 | 0.00018624 | 6.65E-05 | 0.00022032 | 0.00020894 | k__Bacteria | p__Gemmatim | c__Gemm-1 | o__ | f__ | g__ | 1.87E-03 |
| 411759 | 0.00010257 | 0.0001247 | 0.00013716 | 8.07E-05 | 7.20E-05 | 0.00011445 | 0.00069843 | 5.88E-05 | 3.64E-05 | 0.00011273 | 7.50E-05 | 0.00023121 | k__Bacteria | p__Actinobact | c__Actinobact | o__Actinomycetales | f__Microbacteriacea | g__Agromyces | 1.84E-03 |
| 35775 | 0.00028634 | 0.00015276 | 0.00027603 | 0.00010951 | 3.32E-05 | 0.00016213 | 7.34E-05 | 0.00015929 | 0.00017005 | 0.00022546 | 7.03E-05 | 8.05E-05 | k__Bacteria | p__Firmicutes | c__Bacilli | o__Lactobacillales | f__Lactobacillaceae | g__ | 1.80E-03 |
| 26840 | 0.00016454 | 0.0001247 | 0.00016288 | 0.00015755 | 9.96E-05 | 0.00017167 | 0.00013076 | 0.0002132 | 9.11E-05 | 4.34E-05 | 0.00019923 | 0.00023806 | k__Bacteria | p__Proteobac | c__Gammapro | o__Alteromonadales | f__Alteromonadace | g__Cellvibrio | 1.80E-03 |
| 614149 | 0.00014531 | 4.36E-05 | 0.00021774 | 0.00010951 | 5.54E-05 | 0.00028294 | 9.57E-05 | 0.00027447 | 9.72E-05 | 0.00017632 | 8.20E-05 | 0.0001901 | k__Bacteria | p__Proteobac | c__Alphaprote | o__Rhizobiales | f__Beijerinckiaceae | g__ | 1.77E-03 |
| 369946 | 1.07E-05 | 9.66E-05 | 8.92E-05 | 8.26E-05 | 6.09E-05 | 0.00015577 | 0.00033167 | 0.00038475 | 0.00011134 | 9.25E-05 | 0.00025548 | 8.56E-05 | k__Bacteria | p__Bacteroide | c__Bacteroidia | o__Bacteroidales | f__ | g__ | 1.76E-03 |
| 610478 | 4.91E-05 | 0.00020888 | 0.00078523 | 2.31E-05 | 9.96E-05 | 2.54E-05 | 0.00022324 | 0.00015684 | 1.01E-05 | 4.34E-05 | 0.00011954 | 5.14E-06 | k__Bacteria | p__Firmicutes | c__Clostridia | o__Clostridiales | f__Clostridiaceae | g__Proteiniclasticum | 1.75E-03 |
| 577684 | 8.55E-06 | 0.00019329 | 0.00028117 | 1.54E-05 | 0 | 0.00035606 | 0.00019454 | 4.17E-05 | 5.67E-05 | 5.49E-05 | 0.00039377 | 0.00015242 | k__Bacteria | p__Chloroflexi | c__Thermomic | o__AKYG1722 | f__ | g__ | 1.75E-03 |
| 614514 | 8.55E-05 | 0.00015276 | 0.00050577 | 5.38E-05 | 0.00011624 | 6.99E-05 | 0.0001467 | 4.17E-05 | 1.21E-05 | 9.54E-05 | 0.00021095 | 0.00024833 | k__Bacteria | p__Acidobacte | c__Solibactere | o__Solibacterales | f__ | g__ | 1.74E-03 |
| 615312 | 7.69E-05 | 0.00014029 | 0.0001663 | 0.00026514 | 0.00015498 | 0.00027976 | 0.00013713 | 0.00016909 | 0.00010122 | 5.78E-05 | 5.16E-05 | 0.00013359 | k__Bacteria | p__Proteobac | c__Alphaprote | o__Rhodospirillales | f__Acetobacteracea | g__Roseomonas | 1.73E-03 |
| 611559 | 5.56E-05 | 0.00050504 | 9.26E-05 | 0.00022863 | 3.32E-05 | 4.77E-05 | 0.00018497 | 9.80E-05 | 0.00021053 | 4.91E-05 | 8.44E-05 | 0.00014215 | k__Bacteria | p__Proteobac | c__Alphaprote | o__Rhodospirillales | f__ | g__ | 1.73E-03 |
| 25217 | 0.00027352 | 4.05E-05 | 0.00028117 | 0.00017868 | 0 | 5.09E-05 | 8.29E-05 | 0.00013968 | 0.00012146 | 4.62E-05 | 0.00042893 | 7.54E-05 | k__Bacteria | p__Proteobac | c__Gammapro | o__Xanthomonadales | f__Xanthomonadac | g__Pseudoxanthomon | 1.72E-03 |
| 156955 | 7.05E-05 | 0.00016211 | 0.00021088 | 0.00022095 | 0.00011624 | 0.00011763 | 0.00024876 | 9.31E-05 | 0.00014575 | 6.36E-05 | 0.00012657 | 0.0001353 | k__Bacteria | p__Firmicutes | c__Bacilli | o__Bacillales | f__Planococcaceae | g__Planococcus | 1.71E-03 |
| 372548 | 5.98E-05 | 7.17E-05 | 5.49E-05 | 8.26E-05 | 0.00018819 | 3.81E-05 | 0.00080367 | 5.64E-05 | 6.88E-05 | 2.89E-06 | 0.00018985 | 6.68E-05 | k__Bacteria | p__Bacteroide | c__Flavobacte | o__Flavobacteriales | f__Flavobacteriacea | g__ | 1.68E-03 |
| 50867 | 0.00027138 | 2.81E-05 | 0.0001063 | 0.00016139 | 4.43E-05 | 0.00012716 | 9.89E-05 | 0.00016664 | 0.00014373 | 0.00012718 | 7.97E-05 | 0.00030828 | k__Bacteria | p__Proteobac | c__Gammapro | o__Enterobacteriales | f__Enterobacteriace | g__Serratia | 1.66E-03 |
| 339828 | 8.97E-05 | 0.00039904 | 0.00025203 | 1.34E-05 | 0 | 0.00019392 | 0.00012438 | 8.58E-05 | 2.02E-05 | 9.83E-05 | 0.00014298 | 0.00022093 | k__Bacteria | p__Proteobac | c__Deltaprote | o__Myxococcales | f__ | g__ | 1.64E-03 |
| 447987 | 7.69E-05 | 7.79E-05 | 0.00010287 | 0.00010183 | 2.77E-05 | 0.00018756 | 0.00018816 | 0.00016909 | 6.48E-05 | 0.00020812 | 0.00021798 | 0.00021751 | k__Bacteria | p__Actinobact | c__Actinobact | o__Actinomycetales | f__Frankiaceae | g__ | 1.64E-03 |
| 430625 | 5.77E-05 | 0.00011535 | 0.00021602 | 0.0001537 | 0.00014391 | 0.00024479 | 0.00016584 | 9.56E-05 | 0.00010527 | 0.00011562 | 3.75E-05 | 0.00017983 | k__Bacteria | p__Actinobact | c__Actinobact | o__Actinomycetales | f__Intrasporangiace | g__Marihabitans | 1.63E-03 |
| 449755 | 9.62E-05 | 0.00015899 | 0.0002246 | 0.00012104 | 2.77E-05 | 9.54E-05 | 0.0001754 | 0.00012743 | 0.00026722 | 5.49E-05 | 0.00013594 | 0.00013016 | k__Bacteria | p__Actinobact | c__Actinobact | o__Actinomycetales | f__Propionibacteria | g__Tessaracoccus | 1.61E-03 |
| 413470 | 7.91E-05 | 0.00017458 | 0.00013201 | 0.00015947 | 9.41E-05 | 9.86E-05 | 0.00021049 | 0.00021565 | 0.00011944 | 0.0001214 | 2.81E-05 | 0.00018154 | k__Bacteria | p__Actinobact | c__Actinobact | o__Actinomycetales | f__Actinomycetacea | g__Varibaculum | 1.61E-03 |
| 12220 | 0.00024787 | 0.00010911 | 0.00033775 | 0.00014986 | 0.00021587 | 5.72E-05 | 8.29E-05 | 2.45E-05 | 7.09E-05 | 3.47E-05 | 5.63E-05 | 0.00018668 | k__Bacteria | p__Firmicutes | c__Bacilli | o__Lactobacillales | f__Enterococcaceae | g__Vagococcus | 1.57E-03 |
| 461352 | 4.27E-05 | 0.00013094 | 7.20E-05 | 0.00017868 | 0.00049262 | 4.77E-05 | 7.02E-05 | 3.92E-05 | 0.00022471 | 5.20E-05 | 0.00011016 | 9.59E-05 | k__Bacteria | p__Proteobac | c__Gammapro | o__Aeromonadales | f__Aeromonadacea | g__Aeromonas | 1.56E-03 |
| 619233 | 5.56E-05 | 8.42E-05 | 0.00024688 | 0.00010183 | 5.54E-06 | 0.00010491 | 9.25E-05 | 0.00029897 | 0.00012551 | 7.52E-05 | 0.00016641 | 0.0001764 | k__Bacteria | p__Planctomy | c__Phycispha | o__WD2101 | f__ | g__ | 1.53E-03 |
| 594161 | 5.56E-05 | 0.00019952 | 7.54E-05 | 0.00018252 | 0.00018819 | 7.31E-05 | 6.06E-05 | 0.00012253 | 0.00012551 | 9.25E-05 | 0.0001922 | 0.00015585 | k__Bacteria | p__Actinobact | c__Coriobacte | o__Coriobacteriales | f__Coriobacteriacea | g__Collinsella | 1.52E-03 |
| 2705 | 0.00015172 | 6.86E-05 | 0.00012173 | 0.00010567 | 4.98E-05 | 8.58E-05 | 0.00013076 | 8.33E-05 | 0.00011944 | 0.0001214 | 0.0003633 | 0.00011817 | k__Bacteria | p__Firmicutes | c__Bacilli | o__Lactobacillales | f__Enterococcaceae | g__ | 1.52E-03 |
| 593922 | 7.05E-05 | 8.42E-05 | 6.17E-05 | 9.41E-05 | 0.00017159 | 0.00029565 | 0.00016903 | 2.45E-05 | 9.31E-05 | 0.00020523 | 9.14E-05 | 0.00015071 | k__Bacteria | p__Synergiste | c__Synergistia | o__Synergistales | f__Dethiosulfovibrio | g__TG5 | 1.51E-03 |
| 591402 | 0.00013889 | 0.00090097 | 5.49E-05 | 3.84E-05 | 0 | 1.91E-05 | 6.06E-05 | 3.92E-05 | 2.02E-06 | 0 | 0.00018517 | 6.85E-05 | k__Bacteria | p__Cyanobact | c__Chloroplas | o__Chlorophyta | f__ | g__ | 1.51E-03 |
| 597454 | 8.76E-05 | 0.00029617 | 0.00018859 | 7.69E-05 | 0.00015498 | 0.00013988 | 7.97E-05 | 0.00013968 | 2.23E-05 | 8.96E-05 | 0.00018517 | 2.91E-05 | k__Bacteria | p__SR1 | c__ | o__ | f__ | g__ | 1.49E-03 |
| 280929 | 8.76E-05 | 0.00013717 | 1.37E-05 | 0.00014217 | 5.54E-05 | 8.58E-05 | 0.00057405 | 0.00011763 | 8.70E-05 | 0.00010117 | 3.98E-05 | 4.80E-05 | k__Bacteria | p__Firmicutes | c__Clostridia | o__Clostridiales | f__Veillonellaceae | g__Schwartzia | 1.49E-03 |
| 452363 | 8.55E-05 | 0.00015899 | 0.00017488 | 0.00010375 | 8.86E-05 | 0.00020664 | 0.00013395 | 0.00014949 | 0.00014373 | 0.00011851 | 3.05E-05 | 9.42E-05 | k__Bacteria | p__Actinobact | c__Actinobact | o__Actinomycetales | f__Propionibacteria | g__Microlunatus | 1.49E-03 |
| 610017 | 8.55E-05 | 3.74E-05 | 4.97E-05 | 0.0001537 | 0.00010517 | 0.00019392 | 0.00033805 | 7.35E-05 | 0.00014575 | 7.52E-05 | 9.61E-05 | 0.00013016 | k__Bacteria | p__Firmicutes | c__Clostridia | o__ | f__ | g__ | 1.48E-03 |
| 597412 | 5.77E-05 | 0.00051439 | 9.43E-05 | 8.65E-05 | 3.87E-05 | 0.00014942 | 8.93E-05 | 5.88E-05 | 6.28E-05 | 4.91E-05 | 0.00016407 | 0.00011817 | k__Bacteria | p__Chloroflexi | c__Gitt-GS-13 | o__ | f__ | g__ | 1.48E-03 |
| 405422 | 6.41E-06 | 2.81E-05 | 0.00044576 | 0.0001268 | 0.00017712 | 0.00020664 | 0.00011162 | 5.39E-05 | 6.07E-05 | 1.45E-05 | 0.00012657 | 0.00012331 | k__Bacteria | p__Actinobact | c__Actinobact | o__Actinomycetales | f__Actinomycetacea | g__Mobiluncus | 1.48E-03 |
| 447009 | 7.27E-05 | 9.04E-05 | 0.00015259 | 0.00022671 | 8.86E-05 | 6.99E-05 | 0.00025832 | 6.62E-05 | 0.00010729 | 9.25E-05 | 0.00011719 | 0.00011646 | k__Bacteria | p__Actinobact | c__Actinobact | o__Actinomycetales | f__Intrasporangiace | g__Tetrasphaera | 1.46E-03 |
| 659443 | 1.92E-05 | 4.68E-05 | 0.00051434 | 7.11E-05 | 2.21E-05 | 3.81E-05 | 0.00033805 | 8.82E-05 | 3.44E-05 | 5.78E-05 | 0.00015001 | 6.34E-05 | k__Bacteria | p__Firmicutes | c__Clostridia | o__Clostridiales | f__[Tissierellaceae] | g__ | 1.44E-03 |
| 471947 | 8.12E-05 | 6.24E-06 | 0.00016802 | 9.80E-05 | 1.11E-05 | 0.00016849 | 0.00015627 | 0.00015684 | 0.00015183 | 0.00014163 | 0.00015001 | 0.00014729 | k__Bacteria | p__Actinobact | c__Actinobact | o__Actinomycetales | f__Nakamurellaceae | g__ | 1.44E-03 |
| 405576 | 9.19E-05 | 3.12E-06 | 0.00012173 | 7.30E-05 | 4.98E-05 | 0.00011763 | 8.61E-05 | 0.00044601 | 6.88E-05 | 7.52E-05 | 0.00018048 | 0.00010618 | k__Bacteria | p__Actinobact | c__Actinobact | o__Actinomycetales | f__Nocardioidaceae | g__Pimelobacter | 1.42E-03 |
| 589148 | 8.97E-05 | 0.00014652 | 0.00027432 | 8.45E-05 | 7.75E-05 | 0.00011127 | 0.0001754 | 0.00011518 | 4.45E-05 | 8.38E-05 | 8.67E-05 | 0.0001216 | k__Bacteria | p__TM7 | c__TM7-3 | o__EW055 | f__ | g__ | 1.41E-03 |
| 601221 | 5.56E-05 | 0.00020576 | 5.14E-06 | 3.84E-05 | 3.32E-05 | 6.68E-05 | 0.00066973 | 0.00010538 | 7.09E-05 | 7.52E-05 | 5.63E-05 | 2.57E-05 | k__Bacteria | p__TM7 | c__TM7-3 | o__I025 | f__Rs-045 | g__ | 1.41E-03 |
| 430978 | 6.84E-05 | 6.55E-05 | 0.00010115 | 0.00016715 | 5.54E-06 | 0.00010491 | 0.00028703 | 7.84E-05 | 0.00010324 | 0.00016476 | 6.09E-05 | 0.00019695 | k__Bacteria | p__Actinobact | c__Actinobact | o__Actinomycetales | f__Beutenbergiacea | g__Serinibacter | 1.40E-03 |
| 12259 | 0.00031625 | 4.05E-05 | 0.00040805 | 5.38E-05 | 1.11E-05 | 5.40E-05 | 6.38E-05 | 9.80E-05 | 9.11E-05 | 8.09E-05 | 5.39E-05 | 0.0001216 | k__Bacteria | p__Proteobac | c__Gammapro | o__Vibrionales | f__Pseudoalteromo | g__Pseudoalteromona | 1.39E-03 |
| 12136 | 8.12E-05 | 0.00042399 | 0.00014745 | 3.46E-05 | 0.00022694 | 3.18E-05 | 0.00024238 | 6.86E-05 | 5.06E-05 | 3.18E-05 | 2.34E-05 | 2.40E-05 | k__Bacteria | p__Firmicutes | c__Bacilli | o__Bacillales | f__Planococcaceae | g__Lysinibacillus | 1.39E-03 |
| 34865 | 4.49E-05 | 0.00021199 | 0.00015087 | 0.0001172 | 7.75E-05 | 4.77E-05 | 7.02E-05 | 0.00019605 | 0.00020649 | 4.62E-05 | 7.73E-05 | 0.00013701 | k__Bacteria | p__Firmicutes | c__Erysipelotri | o__Erysipelotrichales | f__Erysipelotrichace | g__[Eubacterium] | 1.38E-03 |
| 447927 | 4.49E-05 | 0.00027434 | 8.06E-05 | 7.49E-05 | 3.32E-05 | 0.00015895 | 0.00011162 | 0.0002132 | 7.09E-05 | 0.00013296 | 0 | 0.00018325 | k__Bacteria | p__Actinobact | c__MB-A2-108 | o__0319-7L14 | f__ | g__ | 1.38E-03 |
| 363860 | 0.00010898 | 2.81E-05 | 3.94E-05 | 0.00017868 | 0 | 6.99E-05 | 6.06E-05 | 0.00017889 | 2.43E-05 | 0.0001214 | 0.00044065 | 0.00012331 | k__Bacteria | p__Bacteroide | c__Cytophagia | o__Cytophagales | f__Cytophagaceae | g__Dyadobacter | 1.37E-03 |
| 439112 | 1.50E-05 | 0.0001777 | 0.00019202 | 9.99E-05 | 3.32E-05 | 0.00010491 | 0.00022643 | 0.00015929 | 6.88E-05 | 6.94E-05 | 0.00012422 | 0.00010105 | k__Bacteria | p__Actinobact | c__Actinobact | o__Actinomycetales | f__Dermatophilacea | g__Dermatophilus | 1.37E-03 |
| 615429 | 1.50E-05 | 0.00013094 | 5.14E-05 | 6.53E-05 | 0 | 0.00075662 | 0.00010205 | 6.13E-05 | 1.82E-05 | 1.73E-05 | 4.45E-05 | 7.71E-05 | k__Bacteria | p__Proteobac | c__Alphaprote | o__Caulobacterales | f__Caulobacteracea | g__Mycoplana | 1.34E-03 |
| 18480 | 3.85E-05 | 0.00016835 | 4.11E-05 | 5.00E-05 | 7.20E-05 | 1.27E-05 | 0.00063465 | 6.37E-05 | 3.44E-05 | 6.94E-05 | 9.38E-05 | 5.14E-05 | k__Bacteria | p__Firmicutes | c__Bacilli | o__Bacillales | f__Planococcaceae | g__Solibacillus | 1.33E-03 |
| 459088 | 5.13E-05 | 0.00022135 | 0.00015259 | 8.26E-05 | 2.77E-05 | 5.72E-05 | 0.00036994 | 9.80E-05 | 3.44E-05 | 4.34E-05 | 5.86E-05 | 9.59E-05 | k__Bacteria | p__Actinobact | c__Actinobact | o__Actinomycetales | f__Nocardiaceae | g__Nocardia | 1.29E-03 |
| 334273 | 9.40E-05 | 2.81E-05 | 0.0002126 | 9.99E-05 | 2.77E-05 | 0.000213 | 4.78E-05 | 4.90E-05 | 0.00010122 | 8.96E-05 | 0.00020157 | 0.00011817 | k__Bacteria | p__Firmicutes | c__Bacilli | o__Bacillales | f__Paenibacillaceae | g__Paenibacillus | 1.28E-03 |
| 76876 | 0.00011753 | 0 | 0.00018002 | 0.00013065 | 9.41E-05 | 5.72E-05 | 4.46E-05 | 0.00022791 | 8.30E-05 | 2.60E-05 | 0.00016407 | 0.00014557 | k__Bacteria | p__Firmicutes | c__Erysipelotri | o__Erysipelotrichales | f__Erysipelotrichace | g__Catenibacterium | 1.27E-03 |
| 584115 | 1.71E-05 | 0 | 0.00013201 | 0.00023632 | 9.41E-05 | 6.04E-05 | 0.000118 | 7.35E-05 | 0.00026114 | 0.00011851 | 5.63E-05 | 0.00010276 | k__Bacteria | p__Fusobacte | c__Fusobacte | o__Fusobacteriales | f__Leptotrichiaceae | g__ | 1.27E-03 |
| 39298 | 6.84E-05 | 0.00014029 | 0.0001183 | 0.00012296 | 4.43E-05 | 6.36E-06 | 0.00026151 | 6.86E-05 | 3.64E-05 | 0.00017343 | 0.00010079 | 0.00010618 | k__Bacteria | p__Proteobac | c__Gammapro | o__Enterobacteriales | f__Enterobacteriace | g__Gluconacetobacter | 1.25E-03 |
| 613990 | 8.76E-05 | 4.05E-05 | 0.00012859 | 7.88E-05 | 5.54E-06 | 0.00014306 | 0.00011162 | 0.0002034 | 0.00012146 | 4.62E-05 | 0.00011016 | 0.00015928 | k__Bacteria | p__Proteobac | c__Alphaprote | o__Rhodospirillales | f__Acetobacteracea | g__Belnapia | 1.24E-03 |
| 410289 | 2.99E-05 | 0.00036163 | 0.0001543 | 3.27E-05 | 5.54E-06 | 0.00010809 | 9.25E-05 | 8.09E-05 | 4.05E-05 | 2.60E-05 | 0.00018048 | 9.59E-05 | k__Bacteria | p__Gemmatim | c__Gemmatim | o__Gemmatimonadales | f__ | g__ | 1.21E-03 |
| 616178 | 0.00013035 | 1.25E-05 | 2.57E-05 | 0.00014602 | 0.0001107 | 3.18E-05 | 7.65E-05 | 0.00015684 | 0.0002328 | 6.07E-05 | 8.44E-05 | 0.0001079 | k__Bacteria | p__Proteobac | c__Gammapro | o__Oceanospirillales | f__Alcanivoracaceae | g__Alcanivorax | 1.18E-03 |

| | | | | | | | | | | | | | | | | | | | |
|---|---|---|---|---|---|---|---|---|---|---|---|---|---|---|---|---|---|---|---|
| 476037 | 0.0001218 | 4.36E-05 | 0.00012001 | 0.00013257 | 0.00017159 | 7.95E-05 | 0.00010205 | 6.13E-05 | 7.90E-05 | 8.38E-05 | 4.92E-05 | 0.00011817 | k__Bacteria | p__Actinobact | c__Actinobact | o__Actinomycetales | f__Microbacteriacea | g__Pseudoclavibacter | 1.16E-03 |
| 9274 | 4.06E-05 | 8.11E-05 | 0.00017659 | 6.92E-05 | 2.77E-05 | 0.00010809 | 9.25E-05 | 0.00013968 | 0.00013766 | 0.00010695 | 0.00010782 | 6.51E-05 | k__Bacteria | p__Proteobac | c__Betaprotec | o__Burkholderiales | f__Comamonadace | g__Polaromonas | 1.15E-03 |
| 578858 | 4.49E-05 | 0.00011223 | 4.11E-05 | 0.00014025 | 0.00025461 | 6.36E-06 | 6.70E-05 | 0.00016174 | 6.48E-05 | 7.23E-05 | 0.00010079 | 7.19E-05 | k__Bacteria | p__Fusobacte | c__Fusobacte | o__Fusobacteriales | f__Leptotrichiaceae | g__Sneathia | 1.14E-03 |
| 610360 | 9.83E-05 | 0.00014341 | 0.00010458 | 9.22E-05 | 2.21E-05 | 1.91E-05 | 0.00018178 | 0.00010538 | 8.50E-05 | 6.65E-05 | 0.0001336 | 8.22E-05 | k__Bacteria | p__Proteobac | c__Alphaprote | o__Rhizobiales | f__Phyllobacteriace | g__ | 1.13E-03 |
| 13889 | 8.76E-05 | 0.00015899 | 9.60E-05 | 7.30E-05 | 9.41E-05 | 8.58E-05 | 0.00016903 | 4.41E-05 | 7.29E-05 | 6.94E-05 | 0.00010079 | 7.88E-05 | k__Bacteria | p__Firmicutes | c__Clostridia | o__Clostridiales | f__Peptococcaceae | g__Peptococcus | 1.13E-03 |
| 610678 | 0.00043592 | 0.00014029 | 0.00016288 | 7.88E-05 | 2.77E-05 | 6.36E-05 | 2.87E-05 | 3.19E-05 | 8.10E-06 | 4.05E-05 | 6.56E-05 | 3.77E-05 | k__Bacteria | p__Proteobac | c__Alphaprote | o__Rhizobiales | f__Rhizobiaceae | g__Rhizobium | 1.12E-03 |
| 405302 | 1.28E-05 | 3.74E-05 | 0.00017488 | 2.88E-05 | 8.86E-05 | 0.00018756 | 7.02E-05 | 0.00017399 | 4.66E-05 | 2.60E-05 | 0.00021329 | 5.82E-05 | k__Bacteria | p__Planctomy | c__Planctomy | o__Pirellulales | f__Pirellulaceae | g__ | 1.12E-03 |
| 12175 | 6.41E-05 | 9.66E-05 | 0.00015602 | 0.00039002 | 5.54E-06 | 2.86E-05 | 2.23E-05 | 2.94E-05 | 0.00021256 | 3.18E-05 | 9.38E-06 | 6.85E-05 | k__Bacteria | p__Proteobac | c__Gammapro | o__Vibrionales | f__Vibrionaceae | g__ | 1.11E-03 |
| 32085 | 2.35E-05 | 3.12E-06 | 0.00026232 | 0.00013641 | 5.54E-06 | 1.91E-05 | 2.87E-05 | 6.86E-05 | 8.30E-05 | 0.00011562 | 0.00013126 | 0.00022093 | k__Bacteria | p__Firmicutes | c__Bacilli | o__Bacillales | f__Bacillaceae | g__Anoxybacillus | 1.10E-03 |
| 611399 | 0 | 0.00020264 | 3.77E-05 | 0.00018444 | 1.11E-05 | 8.90E-05 | 8.61E-05 | 0.00031368 | 3.85E-05 | 3.18E-05 | 4.22E-05 | 4.80E-05 | k__Bacteria | p__Chloroflexi | c__Anaeroline | o__Ardenscatenales | f__Ardenscatenace | g__Ardenscatena | 1.09E-03 |
| 415522 | 2.78E-05 | 9.66E-05 | 0.00019374 | 9.41E-05 | 3.87E-05 | 0.0001208 | 9.89E-05 | 8.82E-05 | 0.00011134 | 4.91E-05 | 4.22E-05 | 0.00010447 | k__Bacteria | p__Actinobact | c__Actinobact | o__Actinomycetales | f__Micrococcaceae | g__Acaricomes | 1.07E-03 |
| 627644 | 4.27E-06 | 0.00025876 | 4.63E-05 | 0.00014217 | 7.20E-05 | 0.0001971 | 6.06E-05 | 0.00012988 | 1.42E-05 | 5.20E-05 | 3.52E-05 | 3.43E-05 | k__Bacteria | p__Firmicutes | c__Clostridia | o__Clostridiales | f__[Tissierellaceae] | g__ph2 | 1.05E-03 |
| 416139 | 3.85E-05 | 0.00018705 | 0.00013373 | 4.61E-05 | 1.66E-05 | 0.00015895 | 9.57E-05 | 7.11E-05 | 9.51E-05 | 8.09E-05 | 6.56E-05 | 5.48E-05 | k__Bacteria | p__Planctomy | c__Planctomy | o__Gemmatales | f__Isosphaeraceae | g__ | 1.04E-03 |
| 437767 | 2.56E-05 | 6.24E-06 | 0.00030003 | 1.92E-05 | 1.11E-05 | 5.72E-05 | 0.00032848 | 4.66E-05 | 0.00012956 | 5.78E-06 | 7.03E-06 | 8.91E-05 | k__Bacteria | p__Actinobact | c__Actinobact | o__Actinomycetales | f__Dermabacterace | g__Devriesea | 1.03E-03 |
| 136919 | 5.77E-05 | 6.86E-05 | 2.57E-05 | 6.15E-05 | 6.09E-05 | 2.54E-05 | 0.00010843 | 0.00014459 | 4.05E-05 | 0.00014163 | 9.38E-05 | 0.00019182 | k__Bacteria | p__Proteobac | c__Gammapro | o__Cardiobacteriales | f__Cardiobacteriace | g__Cardiobacterium | 1.02E-03 |
| 363813 | 3.42E-05 | 0.00018393 | 5.49E-05 | 4.42E-05 | 2.77E-05 | 0.00010173 | 9.57E-05 | 0.00013723 | 5.06E-05 | 9.25E-05 | 3.75E-05 | 0.00015242 | k__Bacteria | p__Bacteroide | c__Bacteroidia | o__Bacteroidales | f__Porphyromonad | g__Tannerella | 1.01E-03 |
| 377133 | 2.56E-05 | 7.79E-05 | 4.46E-05 | 7.49E-05 | 2.21E-05 | 0.00016849 | 5.10E-05 | 0.00017154 | 5.47E-05 | 1.73E-05 | 0.00013594 | 0.00016784 | k__Bacteria | p__Bacteroide | c__Cytophagia | o__Cytophagales | f__Cytophagaceae | g__ | 1.01E-03 |
| 54893 | 4.06E-05 | 0.00011847 | 0.00017316 | 0.00013065 | 2.21E-05 | 6.04E-05 | 5.74E-05 | 9.31E-05 | 0.00011944 | 6.65E-05 | 4.92E-05 | 6.17E-05 | k__Bacteria | p__Proteobac | c__Betaprotec | o__Burkholderiales | f__Comamonadace | g__Acidovorax | 9.93E-04 |
| 13555 | 5.98E-05 | 5.61E-05 | 3.94E-05 | 5.57E-05 | 0.00015498 | 2.86E-05 | 1.28E-05 | 3.19E-05 | 0.00030366 | 1.45E-05 | 3.52E-05 | 0.00018325 | k__Bacteria | p__Firmicutes | c__Clostridia | o__Clostridiales | f__Veillonellaceae | g__Megamonas | 9.76E-04 |
| 423843 | 5.34E-05 | 0.00032734 | 6.17E-05 | 3.65E-05 | 1.66E-05 | 2.23E-05 | 0.00017222 | 6.13E-05 | 6.07E-05 | 2.60E-05 | 3.05E-05 | 0.00010618 | k__Bacteria | p__Actinobact | c__Actinobact | o__Actinomycetales | f__Ruaniaceae | g__Ruania | 9.75E-04 |
| 472018 | 4.91E-05 | 0.00017146 | 4.11E-05 | 0.00011336 | 5.54E-05 | 4.77E-05 | 7.02E-05 | 4.66E-05 | 0.00010324 | 6.36E-05 | 0.00010782 | 0.00010105 | k__Bacteria | p__ | c__ | o__ | f__ | g__ | 9.71E-04 |
| 414321 | 4.27E-05 | 0.00019017 | 8.06E-05 | 0.00011528 | 9.96E-05 | 2.86E-05 | 4.46E-05 | 3.92E-05 | 7.09E-05 | 9.83E-05 | 0.00011719 | 4.28E-05 | k__Bacteria | p__Actinobact | c__Actinobact | o__Actinomycetales | f__Pseudonocardia | g__Saccharopolyspora | 9.70E-04 |
| 12984 | 2.35E-05 | 0.00015276 | 3.09E-05 | 3.65E-05 | 5.54E-06 | 0.00013352 | 0.00010205 | 9.31E-05 | 2.63E-05 | 8.96E-05 | 0.00018985 | 8.56E-05 | k__Bacteria | p__Acidobacte | c__Acidobacte | o__iii1-15 | f__ | g__ | 9.69E-04 |
| 629598 | 8.55E-06 | 0.00038346 | 9.09E-05 | 2.31E-05 | 5.54E-05 | 3.81E-05 | 2.23E-05 | 6.62E-05 | 5.26E-05 | 0.00010984 | 4.69E-05 | 5.82E-05 | k__Bacteria | p__GN02 | c__BD1-5 | o__ | f__ | g__ | 9.55E-04 |
| 449153 | 0.00017736 | 0.00013094 | 6.17E-05 | 0.00010567 | 0 | 3.18E-06 | 7.34E-05 | 0.00017644 | 8.10E-06 | 5.20E-05 | 7.27E-05 | 8.05E-05 | k__Bacteria | p__Actinobact | c__Actinobact | o__Actinomycetales | f__Micromonospora | g__Actinoplanes | 9.42E-04 |
| 612601 | 0.00025215 | 3.43E-05 | 5.83E-05 | 6.92E-05 | 3.32E-05 | 2.86E-05 | 3.51E-05 | 4.41E-05 | 4.25E-05 | 3.18E-05 | 0.00019454 | 0.00011132 | k__Bacteria | p__Proteobac | c__Alphaprote | o__Sphingomonadales | f__Sphingomonada | g__Sphingobium | 9.35E-04 |
| 446745 | 5.56E-05 | 6.55E-05 | 0.00016459 | 3.07E-05 | 3.32E-05 | 2.54E-05 | 0.0003253 | 3.68E-05 | 4.45E-05 | 5.20E-05 | 1.64E-05 | 7.36E-05 | k__Bacteria | p__Actinobact | c__Actinobact | o__Actinomycetales | f__Promicromonosp | g__Cellulosimicrobium | 9.24E-04 |
| 436120 | 2.35E-05 | 0.00015899 | 0.00033604 | 4.42E-05 | 5.54E-06 | 1.59E-05 | 6.70E-05 | 0.00015439 | 3.64E-05 | 5.78E-06 | 1.17E-05 | 5.48E-05 | k__Bacteria | p__Actinobact | c__Actinobact | o__Actinomycetales | f__Micromonospora | g__Micromonospora | 9.14E-04 |
| 411873 | 0.00011325 | 7.48E-05 | 0.0001063 | 7.11E-05 | 1.66E-05 | 5.72E-05 | 0.00015627 | 6.62E-05 | 5.06E-05 | 6.07E-05 | 3.28E-05 | 8.56E-05 | k__Bacteria | p__Actinobact | c__Actinobact | o__Actinomycetales | f__Microbacteriacea | g__Salinibacterium | 8.91E-04 |
| 377163 | 1.07E-05 | 5.61E-05 | 8.23E-05 | 4.23E-05 | 2.21E-05 | 2.23E-05 | 0.00035719 | 7.35E-05 | 0.00012349 | 1.73E-05 | 7.03E-05 | 1.20E-05 | k__Bacteria | p__Bacteroide | c__Cytophagia | o__Cytophagales | f__Cyclobacteriacea | g__ | 8.90E-04 |
| 450257 | 4.27E-05 | 9.35E-05 | 9.26E-05 | 2.88E-05 | 3.87E-05 | 0.00012398 | 0.00011162 | 7.60E-05 | 2.23E-05 | 6.36E-05 | 5.63E-05 | 0.0001353 | k__Bacteria | p__Actinobact | c__Actinobact | o__Actinomycetales | f__Tsukamurellacea | g__Tsukamurella | 8.85E-04 |
| 439282 | 5.98E-05 | 0.00018705 | 4.29E-05 | 5.57E-05 | 0 | 3.50E-05 | 0.00010205 | 6.37E-05 | 8.30E-05 | 3.76E-05 | 2.11E-05 | 0.0001901 | k__Bacteria | p__Actinobact | c__Actinobact | o__Actinomycetales | f__Dermabacterace | g__ | 8.78E-04 |
| 430059 | 6.20E-05 | 1.56E-05 | 0.00011658 | 9.03E-05 | 3.87E-05 | 7.31E-05 | 0.00012119 | 3.68E-05 | 6.28E-05 | 5.20E-05 | 7.97E-05 | 0.00012845 | k__Bacteria | p__Actinobact | c__Actinobact | o__Actinomycetales | f__Dermabacterace | g__Helcobacillus | 8.77E-04 |
| 427333 | 0.00025215 | 1.25E-05 | 3.60E-05 | 5.57E-05 | 1.11E-05 | 4.13E-05 | 0.00022324 | 5.88E-05 | 3.24E-05 | 1.16E-05 | 0.00010079 | 2.23E-05 | k__Bacteria | p__Actinobact | c__Actinobact | o__Actinomycetales | f__Ruaniaceae | g__ | 8.58E-04 |
| 444177 | 1.07E-05 | 0.00050816 | 3.60E-05 | 7.69E-06 | 5.54E-06 | 2.23E-05 | 5.74E-05 | 0.00010293 | 2.02E-06 | 3.76E-05 | 3.75E-05 | 2.40E-05 | k__Bacteria | p__Actinobact | c__Actinobact | o__Actinomycetales | f__Nocardiaceae | g__ | 8.52E-04 |
| 586387 | 7.48E-05 | 0.00011223 | 0.00020745 | 6.53E-05 | 7.20E-05 | 3.18E-06 | 7.02E-05 | 4.90E-05 | 2.02E-05 | 2.89E-05 | 6.33E-05 | 6.68E-05 | k__Bacteria | p__Firmicutes | c__Clostridia | o__Clostridiales | f__Ruminococcacea | g__Oscillospira | 8.33E-04 |
| 109709 | 1.50E-05 | 6.24E-05 | 9.09E-05 | 0.00010375 | 3.32E-05 | 0.00010491 | 4.78E-05 | 0.00013233 | 7.29E-05 | 2.02E-05 | 7.97E-05 | 7.02E-05 | k__Bacteria | p__Firmicutes | c__Erysipelotri | o__Erysipelotrichales | f__Erysipelotrichace | g__ | 8.33E-04 |
| 610692 | 4.49E-05 | 0.00013094 | 9.09E-05 | 4.03E-05 | 1.11E-05 | 3.18E-05 | 8.29E-05 | 0.00023771 | 2.63E-05 | 3.47E-05 | 1.88E-05 | 8.22E-05 | k__Bacteria | p__TM7 | c__TM7-1 | o__ | f__ | g__ | 8.32E-04 |
| 586605 | 1.28E-05 | 0.00010911 | 6.52E-05 | 2.50E-05 | 3.87E-05 | 3.18E-05 | 0.00011481 | 0.00015929 | 4.25E-05 | 5.78E-06 | 0.00021329 | 1.20E-05 | k__Bacteria | p__Actinobact | c__Acidimicrob | o__Acidimicrobiales | f__AKIW874 | g__ | 8.30E-04 |
| 23774 | 7.48E-05 | 1.56E-05 | 2.06E-05 | 0.00011336 | 0.0001107 | 2.23E-05 | 3.51E-05 | 3.92E-05 | 0.00014373 | 1.73E-05 | 3.98E-05 | 0.00019353 | k__Bacteria | p__Proteobac | c__Gammapro | o__Enterobacteriales | f__Enterobacteriace | g__Plesiomonas | 8.26E-04 |
| 593844 | 2.99E-05 | 4.68E-05 | 0.0001063 | 2.31E-05 | 3.87E-05 | 0.00014942 | 9.25E-05 | 7.11E-05 | 4.25E-05 | 0.00011562 | 5.63E-05 | 5.31E-05 | k__Bacteria | p__FBP | c__ | o__ | f__ | g__ | 8.25E-04 |
| 784 | 4.70E-05 | 7.17E-05 | 6.00E-05 | 4.80E-05 | 0 | 6.04E-05 | 4.78E-05 | 9.80E-05 | 3.85E-05 | 6.94E-05 | 0.00016876 | 0.00011303 | k__Bacteria | p__Gemmatim | c__Gemm-3 | o__ | f__ | g__ | 8.23E-04 |
| 611305 | 0 | 8.42E-05 | 8.06E-05 | 1.15E-05 | 6.64E-05 | 7.31E-05 | 4.15E-05 | 0.00015194 | 1.01E-05 | 4.91E-05 | 0.00021095 | 4.28E-05 | k__Bacteria | p__Chloroflexi | c__Anaeroline | o__Caldilineales | f__Caldilineaceae | g__ | 8.22E-04 |
| 406358 | 4.27E-06 | 0.00016211 | 4.80E-05 | 4.23E-05 | 0 | 5.40E-05 | 2.55E-05 | 0.00011273 | 2.63E-05 | 1.16E-05 | 0.00015938 | 0.0001627 | k__Bacteria | p__Gemmatim | c__Gemmatim | o__ | f__ | g__ | 8.09E-04 |
| 444440 | 2.56E-05 | 0.00032734 | 8.57E-06 | 1.15E-05 | 2.21E-05 | 3.50E-05 | 0.00016584 | 2.45E-06 | 8.10E-06 | 4.05E-05 | 6.56E-05 | 9.08E-05 | k__Bacteria | p__Actinobact | c__Actinobact | o__Actinomycetales | f__Gordoniaceae | g__Millisia | 8.03E-04 |
| 20375 | 3.63E-05 | 0.00033981 | 1.03E-05 | 2.69E-05 | 0.00011624 | 5.09E-05 | 9.57E-06 | 4.90E-05 | 7.29E-05 | 3.18E-05 | 2.34E-05 | 2.91E-05 | k__Bacteria | p__Proteobac | c__Betaprotec | o__Methylophilales | f__Methylophilacea | g__ | 7.96E-04 |
| 73893 | 0.00010471 | 3.74E-05 | 1.03E-05 | 2.50E-05 | 1.66E-05 | 9.54E-06 | 1.59E-05 | 3.68E-05 | 8.10E-06 | 5.49E-05 | 3.28E-05 | 0.00044357 | k__Bacteria | p__Proteobac | c__Gammapro | o__Enterobacteriales | f__Enterobacteriace | g__Pantoea | 7.96E-04 |
| 434515 | 1.50E-05 | 0.00015588 | 4.29E-05 | 1.73E-05 | 3.87E-05 | 2.54E-05 | 8.61E-05 | 0.00011518 | 1.82E-05 | 0.00010406 | 9.84E-05 | 5.82E-05 | k__Bacteria | p__Planctomy | c__Planctomy | o__Planctomycetales | f__Planctomycetace | g__Planctomyces | 7.75E-04 |
| 479578 | 1.71E-05 | 0.00012158 | 0.00010973 | 9.03E-05 | 5.54E-06 | 4.45E-05 | 0.0001467 | 1.72E-05 | 9.72E-05 | 4.34E-05 | 5.63E-05 | 2.23E-05 | k__Bacteria | p__Actinobact | c__Actinobact | o__Actinomycetales | f__Cellulomonadace | g__Demequina | 7.72E-04 |
| 431739 | 4.91E-05 | 3.74E-05 | 0.00026917 | 4.03E-05 | 5.54E-05 | 6.36E-06 | 4.78E-05 | 0.00014459 | 4.25E-05 | 1.45E-05 | 2.34E-05 | 3.77E-05 | k__Bacteria | p__Actinobact | c__Actinobact | o__Actinomycetales | f__Actinomycetacea | g__Arcanobacterium | 7.68E-04 |
| 588929 | 1.71E-05 | 8.42E-05 | 2.23E-05 | 7.69E-05 | 3.32E-05 | 5.09E-05 | 0.00011481 | 0.00012498 | 1.01E-05 | 7.80E-05 | 0.00012891 | 2.23E-05 | k__Bacteria | p__TM7 | c__TM7-3 | o__CW040 | f__F16 | g__ | 7.64E-04 |
| 606122 | 2.14E-05 | 0.00020264 | 4.63E-05 | 1.15E-05 | 4.98E-05 | 6.68E-05 | 6.38E-05 | 3.43E-05 | 0.00015993 | 1.45E-05 | 4.45E-05 | 4.28E-05 | k__Bacteria | p__Armatimon | c__[Fimbriimor | o__[Fimbriimonadales] | f__[Fimbriimonadac | g__Fimbriimonas | 7.58E-04 |
| 460367 | 1.92E-05 | 2.49E-05 | 7.03E-05 | 6.53E-05 | 5.54E-05 | 4.45E-05 | 9.57E-05 | 4.66E-05 | 7.49E-05 | 7.52E-05 | 5.39E-05 | 0.00013187 | k__Bacteria | p__Actinobact | c__Actinobact | o__Actinomycetales | f__Actinosynnemata | g__ | 7.58E-04 |
| 650729 | 6.41E-06 | 0.00011535 | 8.23E-05 | 3.46E-05 | 8.86E-05 | 1.59E-05 | 7.34E-05 | 0.00012498 | 5.26E-05 | 1.45E-05 | 8.91E-05 | 5.82E-05 | k__Bacteria | p__Firmicutes | c__Clostridia | o__Clostridiales | f__[Tissierellaceae] | g__GW-34 | 7.56E-04 |
| 452035 | 2.35E-05 | 0.00016211 | 9.26E-05 | 9.61E-06 | 0 | 7.31E-05 | 3.19E-05 | 0 | 3.04E-05 | 4.62E-05 | 7.03E-05 | 0.00020552 | k__Bacteria | p__Actinobact | c__Actinobact | o__Actinomycetales | f__Nocardioidaceae | g__Kribbella | 7.45E-04 |
| 460219 | 5.13E-05 | 3.43E-05 | 1.89E-05 | 4.23E-05 | 0.00014391 | 2.23E-05 | 1.91E-05 | 2.70E-05 | 4.25E-05 | 7.52E-05 | 1.64E-05 | 0.00025176 | k__Bacteria | p__Proteobac | c__Gammapro | o__Enterobacteriales | f__Enterobacteriace | g__Salmonella | 7.45E-04 |
| 36600 | 6.41E-05 | 7.79E-05 | 6.00E-05 | 3.65E-05 | 2.21E-05 | 0.00011127 | 8.93E-05 | 4.90E-05 | 6.88E-05 | 5.49E-05 | 4.92E-05 | 5.31E-05 | k__Bacteria | p__Firmicutes | c__Bacilli | o__Lactobacillales | f__Aerococcaceae | g__Alkalibacterium | 7.36E-04 |
| 373535 | 1.92E-05 | 3.12E-05 | 3.43E-05 | 4.03E-05 | 5.54E-06 | 7.95E-05 | 0.00018816 | 0.00021565 | 1.82E-05 | 8.67E-06 | 3.52E-05 | 5.65E-05 | k__Bacteria | p__Actinobact | c__Actinobact | o__Actinomycetales | f__Nocardiopsaceae | g__Nocardiopsis | 7.32E-04 |
| 427399 | 3.63E-05 | 3.43E-05 | 7.37E-05 | 7.49E-05 | 2.77E-05 | 7.63E-05 | 0.00010843 | 6.37E-05 | 5.47E-05 | 6.36E-05 | 3.98E-05 | 7.88E-05 | k__Bacteria | p__Actinobact | c__Actinobact | o__Actinomycetales | f__Microbacteriacea | g__Rathayibacter | 7.32E-04 |
| 368076 | 7.91E-05 | 4.99E-05 | 0.00010973 | 3.46E-05 | 2.21E-05 | 8.58E-05 | 0.00016265 | 7.84E-05 | 3.04E-05 | 2.60E-05 | 4.69E-06 | 3.77E-05 | k__Bacteria | p__Bacteroide | c__Bacteroidia | o__Bacteroidales | f__Porphyromonad | g__Paludibacter | 7.21E-04 |
| 451699 | 5.98E-05 | 2.18E-05 | 7.20E-05 | 6.34E-05 | 4.98E-05 | 5.72E-05 | 0.000118 | 5.39E-05 | 5.26E-05 | 5.49E-05 | 3.98E-05 | 7.02E-05 | k__Bacteria | p__Actinobact | c__Actinobact | o__Actinomycetales | f__Beutenbergiacea | g__ | 7.14E-04 |
| 36570 | 4.27E-06 | 0.0001247 | 0.0001303 | 8.07E-05 | 2.77E-05 | 1.27E-05 | 3.19E-05 | 8.33E-05 | 4.66E-05 | 2.31E-05 | 6.33E-05 | 7.71E-05 | k__Bacteria | p__Proteobac | c__Betaprotec | o__Burkholderiales | f__Comamonadace | g__Hydrogenophaga | 7.06E-04 |
| 472136 | 1.92E-05 | 0 | 2.57E-05 | 6.72E-05 | 2.77E-05 | 3.81E-05 | 0.00010205 | 3.92E-05 | 0.00013766 | 4.34E-05 | 1.88E-05 | 0.00017983 | k__Bacteria | p__Actinobact | c__Actinobact | o__Actinomycetales | f__Pseudonocardia | g__Amycolatopsis | 6.99E-04 |
| 41606 | 0.00043378 | 2.49E-05 | 2.57E-05 | 1.73E-05 | 5.54E-06 | 9.54E-06 | 3.19E-06 | 2.21E-05 | 5.47E-05 | 2.89E-06 | 3.98E-05 | 3.08E-05 | k__Bacteria | p__Proteobac | c__Betaprotec | o__Burkholderiales | f__Alcaligenaceae | g__Sutterella | 6.70E-04 |
| 76544 | 4.91E-05 | 9.66E-05 | 8.06E-05 | 5.57E-05 | 1.11E-05 | 3.81E-05 | 6.38E-05 | 5.39E-05 | 4.86E-05 | 5.49E-05 | 3.75E-05 | 7.88E-05 | k__Bacteria | p__Proteobac | c__Betaprotec | o__Burkholderiales | f__Comamonadace | g__Methylibium | 6.69E-04 |
| 37102 | 8.55E-06 | 4.36E-05 | 2.57E-05 | 7.30E-05 | 5.54E-06 | 0.00010173 | 1.59E-05 | 6.37E-05 | 8.30E-05 | 8.67E-06 | 0.00016173 | 7.54E-05 | k__Bacteria | p__Proteobac | c__Betaprotec | o__MND1 | f__ | g__ | 6.67E-04 |
| 439293 | 6.41E-06 | 7.48E-05 | 6.52E-05 | 3.46E-05 | 3.32E-05 | 1.59E-05 | 0.00015627 | 3.43E-05 | 8.10E-06 | 1.73E-05 | 0.0001711 | 4.62E-05 | k__Bacteria | p__Actinobact | c__Actinobact | o__Actinomycetales | f__Intrasporangiace | g__Kribbia | 6.63E-04 |
| 593979 | 4.06E-05 | 3.74E-05 | 2.91E-05 | 9.80E-05 | 6.64E-05 | 6.68E-05 | 9.25E-05 | 9.80E-06 | 4.45E-05 | 5.49E-05 | 9.14E-05 | 2.91E-05 | k__Bacteria | p__Firmicutes | c__Clostridia | o__Clostridiales | f__[Acidaminobacte | g__ | 6.61E-04 |
| 615416 | 6.20E-05 | 8.73E-05 | 1.54E-05 | 1.73E-05 | 1.66E-05 | 0.00016531 | 4.46E-05 | 4.41E-05 | 3.04E-05 | 7.52E-05 | 6.09E-05 | 3.94E-05 | k__Bacteria | p__Proteobac | c__Alphaprote | o__Sphingomonadales | f__ | g__ | 6.59E-04 |
| 550 | 3.63E-05 | 5.92E-05 | 3.60E-05 | 0.00010183 | 3.32E-05 | 1.59E-05 | 4.15E-05 | 3.92E-05 | 6.48E-05 | 0.00010117 | 5.63E-05 | 5.65E-05 | k__Bacteria | p__Proteobac | c__Betaprotec | o__Burkholderiales | f__Oxalobacteracea | g__Massilia | 6.42E-04 |
| 612289 | 1.92E-05 | 9.66E-05 | 1.20E-05 | 7.30E-05 | 5.54E-06 | 5.09E-05 | 6.38E-05 | 6.13E-05 | 2.83E-05 | 5.78E-05 | 0.00010313 | 6.68E-05 | k__Bacteria | p__Proteobac | c__Alphaprote | o__Rhizobiales | f__Aurantimonadac | g__ | 6.38E-04 |
| 626088 | 4.27E-06 | 9.04E-05 | 6.69E-05 | 5.00E-05 | 8.86E-05 | 2.86E-05 | 3.19E-06 | 5.15E-05 | 2.23E-05 | 0.0001214 | 7.03E-05 | 1.71E-05 | k__Bacteria | p__Firmicutes | c__Clostridia | o__Clostridiales | f__[Tissierellaceae] | g__Helcococcus | 6.14E-04 |
| 364512 | 4.27E-06 | 0 | 4.80E-05 | 3.07E-05 | 1.11E-05 | 9.22E-05 | 2.87E-05 | 8.33E-05 | 2.83E-05 | 2.89E-05 | 0.00022501 | 3.25E-05 | k__Bacteria | p__Bacteroide | c__Flavobacte | o__Flavobacteriales | f__Flavobacteriacea | g__Gillisia | 6.13E-04 |
| 19322 | 3.63E-05 | 0.00029617 | 1.71E-05 | 5.00E-05 | 2.77E-05 | 2.54E-05 | 1.59E-05 | 2.70E-05 | 1.21E-05 | 2.31E-05 | 1.41E-05 | 5.14E-05 | k__Bacteria | p__Proteobac | c__Gammapro | o__Aeromonadales | f__Aeromonadacea | g__Tolumonas | 5.96E-04 |
| 57194 | 2.14E-05 | 2.49E-05 | 8.57E-06 | 4.03E-05 | 0.0001107 | 1.59E-05 | 1.59E-05 | 3.68E-05 | 3.85E-05 | 2.89E-05 | 0.00019454 | 4.97E-05 | k__Bacteria | p__Proteobac | c__Gammapro | o__Vibrionales | f__Pseudoalteromo | g__Vibrio | 5.86E-04 |
| 449861 | 4.27E-06 | 0.00015588 | 2.91E-05 | 1.92E-06 | 0 | 6.36E-06 | 3.83E-05 | 0.00028917 | 8.10E-06 | 5.78E-06 | 2.34E-06 | 3.08E-05 | k__Bacteria | p__Actinobact | c__Actinobact | o__Actinomycetales | f__Tsukamurellacea | g__Gordonia | 5.72E-04 |
| 2337 | 7.48E-05 | 3.12E-06 | 2.40E-05 | 1.92E-05 | 1.66E-05 | 9.22E-05 | 1.28E-05 | 4.17E-05 | 2.02E-05 | 0.00015031 | 5.86E-05 | 5.48E-05 | k__Bacteria | p__Firmicutes | c__Bacilli | o__Lactobacillales | f__Lactobacillaceae | g__Pediococcus | 5.68E-04 |
| 620328 | 8.55E-05 | 1.56E-05 | 2.74E-05 | 4.03E-05 | 1.66E-05 | 2.23E-05 | 2.87E-05 | 7.11E-05 | 6.48E-05 | 3.18E-05 | 5.86E-05 | 9.93E-05 | k__Bacteria | p__Proteobac | c__Alphaprote | o__Sphingomonadales | f__Sphingomonada | g__Novosphingobium | 5.62E-04 |
| 585808 | 2.14E-06 | 0.00020888 | 3.60E-05 | 0.00010183 | 5.54E-06 | 1.59E-05 | 6.38E-06 | 7.35E-05 | 3.04E-05 | 1.73E-05 | 7.03E-06 | 5.65E-05 | k__Bacteria | p__Firmicutes | c__Clostridia | o__Clostridiales | f__Lachnospiraceae | g__Lachnobacterium | 5.61E-04 |

| | | | | | | | | | | | | | | | | | | | |
|---|---|---|---|---|---|---|---|---|---|---|---|---|---|---|---|---|---|---|---|
| 8114 | 5.98E-05 | 3.12E-06 | 0.00024174 | 4.61E-05 | 5.54E-06 | 4.45E-05 | 3.51E-05 | 7.35E-06 | 2.02E-05 | 3.76E-05 | 1.41E-05 | 3.77E-05 | k__Bacteria | p__Firmicutes | c__Bacilli | o__Bacillales | f__Bacillaceae | g__Geobacillus | 5.53E-04 |
| 136738 | 5.77E-05 | 1.87E-05 | 5.31E-05 | 7.30E-05 | 3.87E-05 | 3.50E-05 | 3.83E-05 | 2.70E-05 | 6.88E-05 | 2.60E-05 | 5.86E-05 | 5.14E-05 | k__Bacteria | p__Proteobac | c__Alphaprote | o__Rhizobiales | f__Rhizobiaceae | g__ | 5.46E-04 |
| 425992 | 4.49E-05 | 0.0001247 | 0.00010973 | 1.34E-05 | 4.43E-05 | 1.27E-05 | 1.28E-05 | 7.35E-06 | 2.63E-05 | 5.20E-05 | 6.33E-05 | 2.40E-05 | k__Bacteria | p__Actinobact | c__Actinobact | o__Actinomycetales | f__Thermomonospo | g__Actinomadura | 5.35E-04 |
| 588884 | 3.21E-05 | 0 | 7.20E-05 | 4.03E-05 | 0 | 0.00010173 | 7.02E-05 | 6.62E-05 | 4.25E-05 | 2.89E-06 | 5.63E-05 | 4.45E-05 | k__Bacteria | p__Actinobact | c__Acidimicrob | o__Acidimicrobiales | f__Iamiaceae | g__Iamia | 5.29E-04 |
| 455404 | 2.78E-05 | 2.18E-05 | 1.89E-05 | 9.03E-05 | 1.66E-05 | 4.13E-05 | 8.61E-05 | 2.70E-05 | 2.23E-05 | 0.00011851 | 3.28E-05 | 2.06E-05 | k__Bacteria | p__Proteobac | c__Alphaprote | o__Rickettsiales | f__mitochondria | g__Oenothera | 5.24E-04 |
| 91647 | 1.92E-05 | 7.17E-05 | 5.31E-05 | 2.31E-05 | 5.54E-06 | 2.86E-05 | 4.15E-05 | 2.70E-05 | 0.00010527 | 6.07E-05 | 1.64E-05 | 6.85E-05 | k__Bacteria | p__Proteobac | c__Betaprotec | o__Neisseriales | f__Neisseriaceae | g__Eikenella | 5.21E-04 |
| 87083 | 0.00019873 | 1.56E-05 | 6.86E-06 | 2.11E-05 | 1.11E-05 | 3.18E-05 | 2.23E-05 | 6.37E-05 | 2.43E-05 | 6.36E-05 | 3.52E-05 | 2.57E-05 | k__Bacteria | p__Proteobac | c__Gammapro | o__Enterobacteriales | f__Enterobacteriace | g__Proteus | 5.20E-04 |
| 120313 | 2.99E-05 | 0.00011535 | 3.09E-05 | 1.73E-05 | 0 | 7.63E-05 | 9.25E-05 | 3.68E-05 | 3.64E-05 | 8.67E-06 | 3.05E-05 | 4.28E-05 | k__Bacteria | p__Actinobact | c__Thermoleo | o__Solirubrobacterales | f__Patulibacteracea | g__ | 5.17E-04 |
| 593763 | 1.92E-05 | 4.99E-05 | 0.00010973 | 5.76E-06 | 5.54E-06 | 0.00011127 | 2.23E-05 | 3.68E-05 | 5.26E-05 | 1.45E-05 | 2.34E-05 | 6.51E-05 | k__Bacteria | p__TM7 | c__TM7-3 | o__I025 | f__ | g__ | 5.16E-04 |
| 26815 | 1.92E-05 | 0.00010911 | 2.40E-05 | 3.46E-05 | 3.32E-05 | 3.81E-05 | 7.02E-05 | 1.96E-05 | 3.04E-05 | 3.76E-05 | 7.03E-05 | 2.91E-05 | k__Bacteria | p__Actinobact | c__Thermoleo | o__Solirubrobacterales | f__Solirubrobactera | g__Solirubrobacter | 5.15E-04 |
| 610497 | 2.35E-05 | 6.86E-05 | 0.00010287 | 3.84E-06 | 5.54E-06 | 3.81E-05 | 4.15E-05 | 6.62E-05 | 4.45E-05 | 2.02E-05 | 3.05E-05 | 6.85E-05 | k__Bacteria | p__Actinobact | c__Acidimicrob | o__Acidimicrobiales | f__EB1017 | g__ | 5.14E-04 |
| 408512 | 8.55E-06 | 4.05E-05 | 3.43E-05 | 3.84E-05 | 1.66E-05 | 5.09E-05 | 0.00021367 | 7.35E-06 | 1.82E-05 | 2.31E-05 | 1.64E-05 | 4.45E-05 | k__Bacteria | p__Actinobact | c__Actinobact | o__Actinomycetales | f__Dietziaceae | g__ | 5.13E-04 |
| 399085 | 0.00013248 | 3.43E-05 | 5.31E-05 | 2.11E-05 | 0 | 1.59E-05 | 3.51E-05 | 8.82E-05 | 5.47E-05 | 3.18E-05 | 1.64E-05 | 2.06E-05 | k__Bacteria | p__Actinobact | c__Actinobact | o__Actinomycetales | f__Jonesiaceae | g__Jonesia | 5.04E-04 |
| 458189 | 1.92E-05 | 9.35E-06 | 8.57E-06 | 2.88E-05 | 2.21E-05 | 0.00017485 | 1.91E-05 | 1.47E-05 | 3.24E-05 | 2.89E-05 | 0.00010782 | 3.25E-05 | k__Bacteria | p__Actinobact | c__Actinobact | o__Actinomycetales | f__Micromonospora | g__Virgisporangium | 4.98E-04 |
| 366460 | 2.56E-05 | 0.00016523 | 1.89E-05 | 1.92E-05 | 0 | 8.27E-05 | 4.46E-05 | 1.72E-05 | 1.21E-05 | 4.91E-05 | 3.28E-05 | 3.08E-05 | k__Bacteria | p__Bacteroide | c__Bacteroidia | o__Bacteroidales | f__Porphyromonada | g__ | 4.98E-04 |
| 100718 | 2.14E-06 | 0.00025876 | 1.71E-05 | 1.15E-05 | 0 | 0 | 3.19E-06 | 6.37E-05 | 2.23E-05 | 2.02E-05 | 9.38E-05 | 3.43E-06 | k__Bacteria | p__Acidobacte | c__Sva0725 | o__Sva0725 | f__ | g__ | 4.96E-04 |
| 1599 | 4.27E-06 | 0.00011535 | 1.89E-05 | 1.73E-05 | 0.00010517 | 1.91E-05 | 2.55E-05 | 3.43E-05 | 3.24E-05 | 5.78E-06 | 6.09E-05 | 4.80E-05 | k__Bacteria | p__Proteobac | c__Deltaprote | o__Bdellovibrionales | f__Bacteriovoracace | g__ | 4.87E-04 |
| 460542 | 0.00010898 | 6.24E-06 | 1.89E-05 | 3.07E-05 | 4.43E-05 | 3.81E-05 | 8.93E-05 | 1.72E-05 | 2.23E-05 | 4.62E-05 | 1.41E-05 | 4.80E-05 | k__Bacteria | p__Actinobact | c__Actinobact | o__Actinomycetales | f__Promicromonosp | g__ | 4.84E-04 |
| 611884 | 8.55E-06 | 4.05E-05 | 5.49E-05 | 5.76E-05 | 1.11E-05 | 4.13E-05 | 1.59E-05 | 6.13E-05 | 5.26E-05 | 2.02E-05 | 4.92E-05 | 6.85E-05 | k__Bacteria | p__Proteobac | c__Alphaprote | o__Rhizobiales | f__Hyphomicrobiace | g__Rhodoplanes | 4.82E-04 |
| 431070 | 1.28E-05 | 0.00013405 | 2.74E-05 | 4.03E-05 | 1.66E-05 | 3.81E-05 | 4.78E-05 | 1.23E-05 | 1.62E-05 | 2.31E-05 | 3.52E-05 | 4.97E-05 | k__Bacteria | p__Actinobact | c__Rubrobact | o__Rubrobacterales | f__Rubrobacteracea | g__ | 4.54E-04 |
| 577937 | 3.21E-05 | 5.61E-05 | 1.37E-05 | 6.72E-05 | 1.66E-05 | 9.54E-06 | 3.51E-05 | 1.23E-05 | 3.64E-05 | 0.00014163 | 1.41E-05 | 1.20E-05 | k__Bacteria | p__Proteobac | c__Alphaprote | o__Rickettsiales | f__mitochondria | g__Sarcandra | 4.47E-04 |
| 420292 | 6.41E-06 | 0.00015899 | 5.66E-05 | 7.69E-06 | 0 | 2.86E-05 | 6.38E-06 | 2.70E-05 | 1.82E-05 | 6.07E-05 | 4.92E-05 | 2.57E-05 | k__Bacteria | p__[Thermi] | c__Deinococci | o__Thermales | f__Thermaceae | g__Thermus | 4.45E-04 |
| 140121 | 1.50E-05 | 0 | 5.83E-05 | 1.54E-05 | 0 | 4.77E-05 | 6.38E-06 | 0.00016664 | 3.85E-05 | 2.89E-06 | 6.56E-05 | 2.06E-05 | k__Bacteria | p__Actinobact | c__Thermoleo | o__Gaiellales | f__ | g__ | 4.37E-04 |
| 19798 | 3.21E-05 | 0.00025876 | 3.43E-06 | 1.34E-05 | 2.77E-05 | 2.23E-05 | 0 | 2.21E-05 | 1.01E-05 | 1.16E-05 | 9.38E-06 | 2.40E-05 | k__Bacteria | p__Proteobac | c__Betaprotec | o__Methylophilales | f__Methylophilaceae | g__Methylotenera | 4.35E-04 |
| 459148 | 3.42E-05 | 3.43E-05 | 2.74E-05 | 4.42E-05 | 5.54E-06 | 3.81E-05 | 5.42E-05 | 3.19E-05 | 3.24E-05 | 5.49E-05 | 3.28E-05 | 4.28E-05 | k__Bacteria | p__Actinobact | c__Actinobact | o__Actinomycetales | f__Microbacteriacea | g__Agrococcus | 4.33E-04 |
| 7064 | 5.13E-05 | 2.49E-05 | 1.89E-05 | 7.30E-05 | 1.11E-05 | 6.36E-06 | 3.19E-05 | 3.19E-05 | 5.26E-05 | 1.45E-05 | 8.67E-05 | 2.23E-05 | k__Bacteria | p__Proteobac | c__Gammapro | o__Alteromonadales | f__[Chromatiaceae] | g__Alishewanella | 4.25E-04 |
| 610700 | 6.84E-05 | 0.00011535 | 4.46E-05 | 5.19E-05 | 0 | 3.81E-05 | 1.28E-05 | 4.90E-05 | 6.07E-06 | 1.16E-05 | 1.64E-05 | 3.43E-06 | k__Bacteria | p__Proteobac | c__Alphaprote | o__BD7-3 | f__ | g__ | 4.18E-04 |
| 41078 | 6.41E-06 | 6.86E-05 | 0.00010973 | 2.69E-05 | 1.66E-05 | 5.09E-05 | 2.23E-05 | 1.96E-05 | 3.24E-05 | 1.16E-05 | 4.45E-05 | 5.14E-06 | k__Bacteria | p__Proteobac | c__Betaprotec | o__Burkholderiales | f__Burkholderiaceae | g__ | 4.15E-04 |
| 459240 | 4.91E-05 | 3.12E-06 | 0.00018345 | 1.15E-05 | 5.54E-05 | 1.27E-05 | 1.59E-05 | 7.35E-06 | 3.24E-05 | 2.89E-06 | 0 | 2.06E-05 | k__Bacteria | p__Actinobact | c__Actinobact | o__Bifidobacteriales | f__Bifidobacteriacea | g__ | 3.94E-04 |
| 369849 | 1.28E-05 | 0 | 3.26E-05 | 4.61E-05 | 1.66E-05 | 2.23E-05 | 3.19E-05 | 3.19E-05 | 2.83E-05 | 2.31E-05 | 0.00012891 | 1.71E-05 | k__Bacteria | p__Proteobac | c__Gammapro | o__Xanthomonadales | f__Sinobacteraceae | g__ | 3.92E-04 |
| 113336 | 3.42E-05 | 4.36E-05 | 8.57E-06 | 8.07E-05 | 4.98E-05 | 3.81E-05 | 3.51E-05 | 0 | 2.43E-05 | 4.05E-05 | 7.03E-06 | 2.74E-05 | k__Bacteria | p__Firmicutes | c__Bacilli | o__Bacillales | f__Listeriaceae | g__ | 3.89E-04 |
| 585628 | 4.27E-06 | 5.30E-05 | 4.46E-05 | 8.65E-05 | 0 | 2.23E-05 | 3.19E-06 | 6.13E-05 | 8.50E-05 | 0 | 0 | 2.91E-05 | k__Bacteria | p__Proteobac | c__Deltaprote | o__Bdellovibrionales | f__Bdellovibrionace | g__Bdellovibrio | 3.89E-04 |
| 76418 | 4.49E-05 | 0.00015276 | 1.89E-05 | 5.00E-05 | 1.11E-05 | 2.23E-05 | 1.59E-05 | 4.66E-05 | 1.01E-05 | 1.16E-05 | 0 | 5.14E-06 | k__Bacteria | p__Proteobac | c__Betaprotec | o__Burkholderiales | f__Comamonadace | g__Lampropedia | 3.89E-04 |
| 89160 | 4.49E-05 | 5.92E-05 | 8.06E-05 | 3.27E-05 | 5.54E-06 | 3.18E-06 | 7.97E-05 | 1.23E-05 | 2.63E-05 | 1.73E-05 | 1.41E-05 | 1.20E-05 | k__Bacteria | p__Proteobac | c__Betaprotec | o__Burkholderiales | f__Alcaligenaceae | g__ | 3.88E-04 |
| 76246 | 1.28E-05 | 1.87E-05 | 3.94E-05 | 0.00012873 | 5.54E-06 | 1.27E-05 | 1.59E-05 | 4.90E-06 | 1.01E-05 | 5.78E-05 | 2.58E-05 | 5.31E-05 | k__Bacteria | p__Proteobac | c__Gammapro | o__Pseudomonadales | f__Moraxellaceae | g__Perlucidibaca | 3.86E-04 |
| 5562 | 8.55E-06 | 0 | 2.06E-05 | 8.45E-05 | 9.41E-05 | 6.36E-06 | 3.51E-05 | 5.15E-05 | 1.01E-05 | 8.67E-06 | 1.88E-05 | 4.62E-05 | k__Bacteria | p__Firmicutes | c__Clostridia | o__Clostridiales | f__Veillonellaceae | g__Phascolarctobacter | 3.84E-04 |
| 610080 | 1.71E-05 | 6.86E-05 | 1.37E-05 | 1.73E-05 | 0 | 3.81E-05 | 9.57E-06 | 6.62E-05 | 5.47E-05 | 0 | 4.22E-05 | 4.62E-05 | k__Bacteria | p__Proteobac | c__Alphaprote | o__Rhizobiales | f__Xanthobacterace | g__Xanthobacter | 3.74E-04 |
| 45332 | 3.42E-05 | 0 | 1.71E-05 | 4.61E-05 | 3.87E-05 | 3.18E-06 | 4.46E-05 | 1.72E-05 | 6.07E-05 | 1.45E-05 | 5.86E-05 | 3.60E-05 | k__Bacteria | p__Proteobac | c__Gammapro | o__Oceanospirillales | f__Endozoicimonac | g__ | 3.71E-04 |
| 405170 | 4.27E-06 | 0 | 4.46E-05 | 3.07E-05 | 3.32E-05 | 2.54E-05 | 4.15E-05 | 5.15E-05 | 8.10E-06 | 3.76E-05 | 7.27E-05 | 1.88E-05 | k__Bacteria | p__Bacteroide | c__Bacteroidia | o__Bacteroidales | f__S24-7 | g__ | 3.68E-04 |
| 593853 | 6.41E-06 | 9.35E-06 | 1.03E-05 | 6.53E-05 | 5.54E-06 | 2.86E-05 | 1.59E-05 | 4.90E-06 | 4.66E-05 | 0.00012718 | 2.81E-05 | 1.88E-05 | k__Bacteria | p__Proteobac | c__Alphaprote | o__Rickettsiales | f__mitochondria | g__Carludovica | 3.67E-04 |
| 402285 | 4.27E-06 | 7.48E-05 | 3.09E-05 | 9.61E-06 | 1.11E-05 | 9.22E-05 | 2.23E-05 | 1.72E-05 | 2.83E-05 | 1.73E-05 | 1.17E-05 | 4.45E-05 | k__Bacteria | p__Bacteroide | c__[Rhodothe | o__[Rhodothermales] | f__Rhodothermacea | g__Rubricoccus | 3.64E-04 |
| 66403 | 1.71E-05 | 3.12E-06 | 8.57E-06 | 7.69E-05 | 6.64E-05 | 4.13E-05 | 9.57E-06 | 2.94E-05 | 3.24E-05 | 5.78E-06 | 1.64E-05 | 5.65E-05 | k__Bacteria | p__Proteobac | c__Betaprotec | o__Burkholderiales | f__Comamonadace | g__Delftia | 3.63E-04 |
| 585576 | 1.71E-05 | 2.49E-05 | 5.14E-06 | 8.84E-05 | 5.54E-06 | 3.50E-05 | 3.51E-05 | 2.21E-05 | 1.82E-05 | 4.91E-05 | 1.41E-05 | 4.62E-05 | k__Bacteria | p__Proteobac | c__Alphaprote | o__Rickettsiales | f__mitochondria | g__Lupinus | 3.61E-04 |
| 44162 | 2.56E-05 | 0 | 3.94E-05 | 3.27E-05 | 2.21E-05 | 3.18E-06 | 9.57E-06 | 7.11E-05 | 6.48E-05 | 5.78E-06 | 9.38E-06 | 6.51E-05 | k__Bacteria | p__Proteobac | c__Gammapro | o__Alteromonadales | f__Psychromonada | g__Psychromonas | 3.49E-04 |
| 417779 | 1.71E-05 | 3.12E-06 | 6.86E-06 | 0.00010951 | 1.66E-05 | 1.91E-05 | 4.15E-05 | 1.47E-05 | 2.23E-05 | 6.65E-05 | 1.17E-05 | 1.20E-05 | k__Bacteria | p__Proteobac | c__Alphaprote | o__Rickettsiales | f__mitochondria | g__Citrullus | 3.41E-04 |
| 77951 | 1.28E-05 | 3.12E-05 | 3.26E-05 | 3.27E-05 | 4.43E-05 | 1.27E-05 | 1.59E-05 | 3.19E-05 | 5.06E-05 | 2.60E-05 | 2.81E-05 | 1.88E-05 | k__Bacteria | p__Proteobac | c__Gammapro | o__ | f__ | g__ | 3.38E-04 |
| 624788 | 1.50E-05 | 9.35E-06 | 2.57E-05 | 1.54E-05 | 5.54E-06 | 1.59E-05 | 6.38E-06 | 2.45E-05 | 3.24E-05 | 5.20E-05 | 6.80E-05 | 6.68E-05 | k__Bacteria | p__Proteobac | c__Alphaprote | o__ | f__ | g__ | 3.37E-04 |
| 71838 | 1.07E-05 | 0 | 1.03E-05 | 2.50E-05 | 4.43E-05 | 1.27E-05 | 3.83E-05 | 3.43E-05 | 5.67E-05 | 3.47E-05 | 4.92E-05 | 2.06E-05 | k__Bacteria | p__Proteobac | c__Gammapro | o__Xanthomonadales | f__Xanthomonadac | g__Thermomonas | 3.37E-04 |
| 26519 | 8.12E-05 | 1.56E-05 | 2.06E-05 | 5.00E-05 | 2.77E-05 | 1.91E-05 | 3.19E-05 | 4.90E-06 | 4.05E-06 | 5.78E-06 | 4.45E-05 | 2.06E-05 | k__Bacteria | p__Proteobac | c__Betaprotec | o__Burkholderiales | f__Comamonadace | g__Rhodoferax | 3.26E-04 |
| 447181 | 1.71E-05 | 9.35E-06 | 1.03E-05 | 4.03E-05 | 1.11E-05 | 5.09E-05 | 3.83E-05 | 7.35E-06 | 4.66E-05 | 2.89E-05 | 4.69E-06 | 5.99E-05 | k__Bacteria | p__Actinobact | c__Actinobact | o__Actinomycetales | f__Actinosynnemata | g__Actinokineospora | 3.25E-04 |
| 5570 | 3.63E-05 | 2.49E-05 | 5.14E-06 | 2.88E-05 | 0 | 1.27E-05 | 9.57E-06 | 3.68E-05 | 1.01E-05 | 6.94E-05 | 6.80E-05 | 2.23E-05 | k__Bacteria | p__Firmicutes | c__Bacilli | o__Bacillales | f__Alicyclobacillacea | g__Alicyclobacillus | 3.24E-04 |
| 466130 | 6.41E-06 | 7.79E-05 | 1.54E-05 | 9.61E-06 | 0 | 6.36E-06 | 2.23E-05 | 5.39E-05 | 5.47E-05 | 5.20E-05 | 7.03E-06 | 1.54E-05 | k__Bacteria | p__Actinobact | c__Actinobact | o__Actinomycetales | f__Actinosynnemata | g__Lentzea | 3.21E-04 |
| 84562 | 6.41E-06 | 6.55E-05 | 2.57E-05 | 2.50E-05 | 1.66E-05 | 1.91E-05 | 6.06E-05 | 7.35E-06 | 2.63E-05 | 2.02E-05 | 3.75E-05 | 1.03E-05 | k__Bacteria | p__Proteobac | c__Betaprotec | o__ASSO-13 | f__ | g__ | 3.21E-04 |
| 142721 | 2.56E-05 | 5.92E-05 | 8.57E-06 | 5.19E-05 | 1.66E-05 | 2.54E-05 | 1.28E-05 | 9.80E-06 | 3.44E-05 | 2.89E-06 | 4.92E-05 | 1.71E-05 | k__Bacteria | p__Proteobac | c__Betaprotec | o__Burkholderiales | f__Comamonadace | g__Limnohabitans | 3.14E-04 |
| 659408 | 2.78E-05 | 9.35E-06 | 1.20E-05 | 5.00E-05 | 1.11E-05 | 9.54E-06 | 1.59E-05 | 2.94E-05 | 1.62E-05 | 8.09E-05 | 1.64E-05 | 2.91E-05 | k__Bacteria | p__Proteobac | c__Alphaprote | o__Rickettsiales | f__mitochondria | g__Carica | 3.08E-04 |
| 7439 | 2.99E-05 | 2.18E-05 | 6.86E-06 | 2.50E-05 | 1.11E-05 | 3.18E-06 | 1.91E-05 | 2.45E-06 | 3.44E-05 | 1.16E-05 | 0.00011719 | 1.03E-05 | k__Bacteria | p__Proteobac | c__Betaprotec | o__Burkholderiales | f__Alcaligenaceae | g__Achromobacter | 2.93E-04 |
| 396704 | 2.56E-05 | 0 | 5.66E-05 | 5.76E-06 | 5.54E-06 | 4.77E-05 | 3.51E-05 | 3.19E-05 | 2.63E-05 | 1.45E-05 | 2.34E-06 | 3.08E-05 | k__Bacteria | p__Bacteroide | c__Cytophagia | o__Cytophagales | f__Cytophagaceae | g__Rhodocytophaga | 2.82E-04 |
| 466157 | 5.56E-05 | 5.61E-05 | 2.74E-05 | 1.34E-05 | 0 | 2.54E-05 | 3.19E-05 | 1.72E-05 | 1.01E-05 | 1.73E-05 | 9.38E-06 | 1.54E-05 | k__Bacteria | p__Actinobact | c__Actinobact | o__Actinomycetales | f__Microbacteriacea | g__Frigoribacterium | 2.79E-04 |
| 365105 | 2.14E-06 | 5.30E-05 | 3.43E-06 | 3.84E-06 | 1.11E-05 | 3.18E-06 | 1.28E-05 | 2.45E-06 | 3.85E-05 | 2.89E-06 | 7.50E-05 | 6.17E-05 | k__Bacteria | p__Bacteroide | c__Flavobacte | o__Flavobacteriales | f__Flavobacteriacea | g__Salinimicrobium | 2.70E-04 |
| 609797 | 2.56E-05 | 8.73E-05 | 1.20E-05 | 7.69E-06 | 2.21E-05 | 3.81E-05 | 1.28E-05 | 0 | 2.23E-05 | 8.67E-06 | 1.17E-05 | 1.88E-05 | k__Bacteria | p__Actinobact | c__Thermoleo | o__Gaiellales | f__AK1AB1_02E | g__ | 2.67E-04 |
| 618698 | 6.41E-06 | 3.74E-05 | 1.37E-05 | 2.11E-05 | 1.66E-05 | 1.27E-05 | 1.28E-05 | 3.43E-05 | 4.05E-06 | 8.67E-06 | 6.33E-05 | 3.60E-05 | k__Bacteria | p__Proteobac | c__Alphaprote | o__Rhodospirillales | f__Acetobacteracea | g__Roseococcus | 2.67E-04 |
| 462943 | 2.14E-05 | 1.25E-05 | 2.57E-05 | 1.15E-05 | 1.66E-05 | 6.36E-06 | 5.42E-05 | 2.45E-05 | 1.01E-05 | 3.47E-05 | 1.17E-05 | 3.60E-05 | k__Bacteria | p__Actinobact | c__Actinobact | o__Actinomycetales | f__Microbacteriacea | g__Clavibacter | 2.65E-04 |
| 577731 | 1.07E-05 | 0 | 3.43E-05 | 1.54E-05 | 2.77E-05 | 2.54E-05 | 1.28E-05 | 3.43E-05 | 1.42E-05 | 5.49E-05 | 2.11E-05 | 1.03E-05 | k__Bacteria | p__Proteobac | c__Alphaprote | o__Rickettsiales | f__mitochondria | g__Abies | 2.61E-04 |
| 411540 | 1.07E-05 | 9.35E-06 | 2.74E-05 | 1.73E-05 | 1.66E-05 | 2.23E-05 | 4.78E-05 | 1.23E-05 | 2.23E-05 | 3.76E-05 | 1.17E-05 | 2.57E-05 | k__Bacteria | p__Actinobact | c__Actinobact | o__Actinomycetales | f__Micrococcaceae | g__Renibacterium | 2.61E-04 |
| 658994 | 1.50E-05 | 3.12E-06 | 1.54E-05 | 1.92E-06 | 5.54E-06 | 3.18E-05 | 7.02E-05 | 1.72E-05 | 3.44E-05 | 8.67E-06 | 2.34E-05 | 3.25E-05 | k__Bacteria | p__Firmicutes | c__Clostridia | o__Clostridiales | f__Clostridiaceae | g__SMB53 | 2.59E-04 |
| 451667 | 1.50E-05 | 6.24E-06 | 2.06E-05 | 2.88E-05 | 2.77E-05 | 2.23E-05 | 5.10E-05 | 2.45E-06 | 1.82E-05 | 5.78E-06 | 1.88E-05 | 4.11E-05 | k__Bacteria | p__Actinobact | c__Actinobact | o__Actinomycetales | f__Promicromonosp | g__Luteimicrobium | 2.58E-04 |
| 263879 | 1.28E-05 | 4.36E-05 | 2.40E-05 | 7.69E-06 | 0 | 3.18E-06 | 1.28E-05 | 2.70E-05 | 3.44E-05 | 2.02E-05 | 4.22E-05 | 2.74E-05 | k__Bacteria | p__Firmicutes | c__Erysipelotri | o__Erysipelotrichales | f__Erysipelotrichace | g__Allobaculum | 2.55E-04 |
| 78052 | 6.41E-06 | 2.81E-05 | 1.37E-05 | 2.50E-05 | 1.11E-05 | 1.91E-05 | 4.15E-05 | 1.23E-05 | 1.62E-05 | 3.18E-05 | 2.34E-05 | 1.37E-05 | k__Bacteria | p__Firmicutes | c__Bacilli | o__Bacillales | f__Planococcaceae | g__Paenisporosarcina | 2.42E-04 |
| 613904 | 1.71E-05 | 3.12E-06 | 6.52E-05 | 4.42E-05 | 0 | 9.54E-06 | 3.19E-06 | 4.90E-06 | 5.06E-05 | 0 | 0 | 4.28E-05 | k__Bacteria | p__Acidobacte | c__Acidobacte | o__Acidobacteriales | f__Acidobacteriacea | g__ | 2.41E-04 |
| 614743 | 1.28E-05 | 3.12E-06 | 7.54E-05 | 1.73E-05 | 2.21E-05 | 4.45E-05 | 0 | 7.35E-06 | 8.10E-06 | 8.67E-06 | 2.81E-05 | 8.56E-06 | k__Bacteria | p__Proteobac | c__Alphaprote | o__Rhodospirillales | f__Rhodospirillacea | g__Azospirillum | 2.36E-04 |
| 370906 | 2.14E-06 | 3.43E-05 | 1.54E-05 | 1.54E-05 | 5.54E-06 | 3.81E-05 | 1.28E-05 | 4.66E-05 | 4.05E-06 | 8.67E-06 | 2.34E-05 | 2.57E-05 | k__Bacteria | p__Actinobact | c__Actinobact | o__Actinomycetales | f__Nocardiopsaceae | g__ | 2.32E-04 |
| 455271 | 6.41E-06 | 6.24E-05 | 1.03E-05 | 9.61E-06 | 0 | 2.86E-05 | 3.19E-06 | 1.47E-05 | 8.10E-06 | 2.60E-05 | 2.34E-06 | 5.65E-05 | k__Bacteria | p__Actinobact | c__Actinobact | o__Bifidobacteriales | f__Bifidobacteriacea | g__Alloscardovia | 2.28E-04 |
| 477237 | 1.07E-05 | 6.24E-06 | 2.23E-05 | 3.46E-05 | 5.54E-06 | 2.54E-05 | 4.15E-05 | 7.35E-06 | 1.42E-05 | 1.45E-05 | 2.11E-05 | 1.88E-05 | k__Bacteria | p__Actinobact | c__Actinobact | o__Actinomycetales | f__Intrasporangiace | g__Intrasporangium | 2.22E-04 |
| 448656 | 6.41E-06 | 2.81E-05 | 3.43E-06 | 2.69E-05 | 5.54E-06 | 9.54E-06 | 1.28E-05 | 4.90E-06 | 3.64E-05 | 1.16E-05 | 1.41E-05 | 6.17E-05 | k__Bacteria | p__Actinobact | c__Actinobact | o__Actinomycetales | f__Micromonospora | g__Pilimelia | 2.21E-04 |
| 221832 | 1.07E-05 | 0 | 2.23E-05 | 7.69E-06 | 1.11E-05 | 1.59E-05 | 3.19E-05 | 2.45E-05 | 1.62E-05 | 1.73E-05 | 7.03E-06 | 5.48E-05 | k__Bacteria | p__Proteobac | c__Gammapro | o__Alteromonadales | f__Alteromonadace | g__ | 2.19E-04 |
| 473425 | 8.55E-06 | 6.24E-06 | 5.31E-05 | 1.15E-05 | 0 | 6.36E-06 | 9.57E-06 | 0.00010293 | 2.02E-06 | 0 | 7.03E-06 | 8.56E-06 | k__Bacteria | p__Actinobact | c__Actinobact | o__Actinomycetales | f__Intrasporangiace | g__Oryzihumus | 2.16E-04 |
| 628631 | 0 | 2.81E-05 | 1.89E-05 | 7.69E-06 | 5.54E-06 | 9.54E-06 | 3.19E-06 | 2.45E-06 | 2.02E-06 | 2.31E-05 | 8.67E-05 | 2.74E-05 | k__Bacteria | p__Cyanobact | c__ML635J-21 | o__ | f__ | g__ | 2.15E-04 |
| 54831 | 4.27E-06 | 0 | 1.71E-06 | 2.11E-05 | 0 | 1.27E-05 | 1.91E-05 | 1.96E-05 | 6.28E-05 | 2.31E-05 | 3.28E-05 | 1.71E-05 | k__Bacteria | p__Proteobac | c__Betaprotec | o__Neisseriales | f__Neisseriaceae | g__Conchiformibius | 2.14E-04 |
| 417156 | 0 | 3.12E-06 | 2.40E-05 | 2.50E-05 | 5.54E-06 | 1.59E-05 | 1.91E-05 | 2.45E-06 | 6.07E-06 | 5.78E-06 | 7.97E-05 | 2.74E-05 | k__Bacteria | p__Nitrospirae | c__Nitrospira | o__Nitrospirales | f__Nitrospiraceae | g__Nitrospira | 2.14E-04 |

| | | | | | | | | | | | | | | | | | | | |
|---|---|---|---|---|---|---|---|---|---|---|---|---|---|---|---|---|---|---|---|
| 138545 | 2.56E-05 | 0 | 1.20E-05 | 3.27E-05 | 0 | 4.13E-05 | 9.57E-06 | 3.68E-05 | 1.01E-05 | 1.45E-05 | 1.17E-05 | 1.71E-05 | k__Bacteria | p__Verrucomi | c__Opitutae | o__Opitutales | f__Opitutaceae | g__Opitutus | 2.11E-04 |
| 463727 | 1.50E-05 | 2.81E-05 | 1.71E-05 | 2.50E-05 | 5.54E-06 | 1.27E-05 | 2.87E-05 | 1.72E-05 | 2.83E-05 | 1.16E-05 | 4.69E-06 | 1.54E-05 | k__Bacteria | p__Actinobact | c__Actinobact | o__Actinomycetales | f__Microbacteriacea | g__Herbiconiux | 2.09E-04 |
| 104655 | 2.14E-05 | 2.49E-05 | 3.60E-05 | 5.19E-05 | 0 | 9.54E-06 | 1.91E-05 | 0 | 6.07E-06 | 2.60E-05 | 9.38E-06 | 1.71E-06 | k__Bacteria | p__Proteobac | c__Betaprotec | o__Burkholderiales | f__Comamonadace | g__Variovorax | 2.06E-04 |
| 98774 | 6.41E-06 | 2.49E-05 | 4.29E-05 | 1.34E-05 | 5.54E-06 | 1.27E-05 | 3.83E-05 | 0 | 2.23E-05 | 8.67E-06 | 1.88E-05 | 8.56E-06 | k__Bacteria | p__Proteobac | c__Betaprotec | o__Burkholderiales | f__Comamonadace | g__Ramlibacter | 2.02E-04 |
| 373556 | 2.14E-06 | 1.25E-05 | 8.57E-06 | 9.61E-06 | 0 | 3.18E-06 | 1.59E-05 | 3.43E-05 | 2.43E-05 | 1.16E-05 | 5.63E-05 | 2.06E-05 | k__Bacteria | p__Bacteroide | c__[Saprospira | o__[Saprospirales] | f__Chitinophagacea | g__Segetibacter | 1.99E-04 |
| 621452 | 8.55E-06 | 1.25E-05 | 3.77E-05 | 7.69E-06 | 0 | 3.81E-05 | 2.87E-05 | 1.47E-05 | 1.42E-05 | 2.02E-05 | 7.03E-06 | 6.85E-06 | k__Bacteria | p__Chloroflexi | c__Thermomic | o__ | f__ | g__ | 1.96E-04 |
| 136800 | 2.14E-06 | 3.12E-06 | 1.03E-05 | 3.84E-06 | 1.11E-05 | 3.18E-06 | 0.00012757 | 2.45E-06 | 1.42E-05 | 2.89E-06 | 2.34E-06 | 8.56E-06 | k__Bacteria | p__Firmicutes | c__Bacilli | o__Bacillales | f__Planococcaceae | g__Viridibacillus | 1.92E-04 |
| 443110 | 1.28E-05 | 9.35E-06 | 1.89E-05 | 2.31E-05 | 0 | 2.54E-05 | 3.19E-05 | 1.23E-05 | 1.82E-05 | 2.31E-05 | 9.38E-06 | 6.85E-06 | k__Bacteria | p__Actinobact | c__Actinobact | o__Actinomycetales | f__Actinosynnemata | g__Kibdelosporangium | 1.91E-04 |
| 621633 | 7.69E-05 | 0 | 6.86E-06 | 3.84E-06 | 2.77E-05 | 3.18E-06 | 6.38E-06 | 2.94E-05 | 1.01E-05 | 1.73E-05 | 7.03E-06 | 1.71E-06 | k__Bacteria | p__Proteobac | c__Alphaprote | o__Rhizobiales | f__Xanthobacterace | g__ | 1.90E-04 |
| 447476 | 2.14E-06 | 9.35E-06 | 5.14E-06 | 5.76E-06 | 0 | 4.45E-05 | 3.19E-06 | 7.35E-06 | 5.26E-05 | 2.89E-06 | 3.05E-05 | 2.06E-05 | k__Bacteria | p__Actinobact | c__Actinobact | o__Actinomycetales | f__Micromonospora | g__Phytohabitans | 1.84E-04 |
| 476050 | 1.92E-05 | 1.25E-05 | 5.14E-06 | 4.23E-05 | 1.66E-05 | 1.27E-05 | 1.91E-05 | 1.47E-05 | 8.10E-06 | 5.78E-06 | 0 | 2.40E-05 | k__Bacteria | p__Proteobac | c__Gammapro | o__Acidithiobacillales | f__ | g__ | 1.80E-04 |
| 340700 | 4.27E-06 | 1.25E-05 | 1.71E-06 | 7.69E-06 | 5.54E-06 | 3.18E-06 | 0 | 6.37E-05 | 3.04E-05 | 1.73E-05 | 1.64E-05 | 1.54E-05 | k__Bacteria | p__Bacteroide | c__Flavobacte | o__Flavobacteriales | f__Blattabacteriacea | g__Blattabacterium | 1.78E-04 |
| 438406 | 8.55E-06 | 9.35E-06 | 1.54E-05 | 2.31E-05 | 0 | 0 | 2.23E-05 | 9.80E-06 | 4.05E-06 | 3.76E-05 | 2.34E-05 | 2.23E-05 | k__Bacteria | p__Actinobact | c__Actinobact | o__Actinomycetales | f__Actinomycetacea | g__Bogoriella | 1.76E-04 |
| 436632 | 3.63E-05 | 9.35E-06 | 1.37E-05 | 2.88E-05 | 0 | 1.59E-05 | 9.57E-06 | 9.80E-06 | 2.02E-05 | 1.45E-05 | 7.03E-06 | 8.56E-06 | k__Bacteria | p__Proteobac | c__Alphaprote | o__Rhodobacterales | f__Rhodobacterace | g__Oceanicella | 1.74E-04 |
| 363900 | 8.55E-05 | 0 | 3.43E-06 | 1.92E-06 | 0 | 1.27E-05 | 3.19E-06 | 4.90E-06 | 4.05E-06 | 1.73E-05 | 7.03E-06 | 2.23E-05 | k__Bacteria | p__Bacteroide | c__Flavobacte | o__Flavobacteriales | f__Cryomorphaceae | g__Fluviicola | 1.62E-04 |
| 373065 | 1.07E-05 | 5.92E-05 | 8.57E-06 | 3.07E-05 | 0 | 3.18E-06 | 3.19E-06 | 1.47E-05 | 8.10E-06 | 0 | 1.41E-05 | 5.14E-06 | k__Bacteria | p__Bacteroide | c__[Saprospira | o__[Saprospirales] | f__Chitinophagacea | g__Sediminibacterium | 1.58E-04 |
| 77857 | 2.14E-05 | 6.24E-06 | 6.86E-06 | 4.42E-05 | 0 | 0 | 1.59E-05 | 1.72E-05 | 8.10E-06 | 1.45E-05 | 2.34E-06 | 1.88E-05 | k__Bacteria | p__Verrucomi | c__Verrucomic | o__Verrucomicrobiales | f__Verrucomicrobia | g__Luteolibacter | 1.55E-04 |
| 578430 | 1.28E-05 | 9.35E-06 | 1.37E-05 | 1.15E-05 | 0 | 0 | 3.19E-06 | 3.43E-05 | 2.02E-05 | 8.67E-06 | 2.81E-05 | 6.85E-06 | k__Bacteria | p__Cyanobact | c__Chloroplas | o__Stramenopiles | f__ | g__ | 1.49E-04 |
| 437958 | 8.55E-06 | 6.24E-06 | 1.37E-05 | 2.69E-05 | 5.54E-06 | 1.91E-05 | 6.38E-06 | 1.23E-05 | 1.21E-05 | 1.45E-05 | 4.69E-06 | 1.54E-05 | k__Bacteria | p__Actinobact | c__Actinobact | o__Actinomycetales | f__Pseudonocardia | g__Jiangella | 1.45E-04 |
| 421538 | 1.07E-05 | 6.24E-06 | 5.14E-06 | 1.54E-05 | 1.66E-05 | 6.36E-06 | 2.87E-05 | 4.90E-06 | 1.82E-05 | 1.73E-05 | 9.38E-06 | 3.43E-06 | k__Bacteria | p__Actinobact | c__Actinobact | o__Actinomycetales | f__Microbacteriacea | g__Curtobacterium | 1.42E-04 |
| 56695 | 1.07E-05 | 1.25E-05 | 1.89E-05 | 7.69E-06 | 0 | 0 | 1.28E-05 | 2.45E-06 | 8.10E-06 | 1.73E-05 | 2.34E-05 | 1.37E-05 | k__Bacteria | p__Proteobac | c__Gammapro | o__Enterobacteriales | f__Enterobacteriace | g__Buchnera | 1.27E-04 |
| 477631 | 6.41E-06 | 0 | 3.43E-06 | 1.54E-05 | 5.54E-06 | 3.18E-06 | 3.19E-05 | 4.90E-06 | 1.62E-05 | 1.73E-05 | 1.17E-05 | 1.03E-05 | k__Bacteria | p__Actinobact | c__Actinobact | o__Actinomycetales | f__Microbacteriacea | g__Mycetocola | 1.26E-04 |
| 471867 | 4.27E-06 | 6.24E-06 | 5.14E-06 | 2.11E-05 | 2.21E-05 | 1.59E-05 | 1.28E-05 | 4.90E-06 | 8.10E-06 | 2.89E-06 | 4.69E-06 | 1.37E-05 | k__Bacteria | p__Actinobact | c__Actinobact | o__Actinomycetales | f__Microbacteriacea | g__Yonghaparkia | 1.22E-04 |
| 10203 | 0 | 3.12E-06 | 5.49E-05 | 1.92E-06 | 5.54E-06 | 6.36E-06 | 3.19E-06 | 4.90E-06 | 6.07E-06 | 2.89E-06 | 9.38E-06 | 1.88E-05 | k__Bacteria | p__Firmicutes | c__Bacilli | o__Bacillales | f__Thermoactinomy | g__Thermoactinomyce | 1.17E-04 |
| 15083 | 2.14E-06 | 3.12E-06 | 1.71E-06 | 1.54E-05 | 0 | 3.18E-06 | 1.59E-05 | 2.70E-05 | 1.42E-05 | 0 | 0 | 1.03E-05 | k__Bacteria | p__Firmicutes | c__Bacilli | o__Lactobacillales | f__Aerococcaceae | g__Marinilactibacillus | 9.29E-05 |
| 455910 | 2.14E-06 | 6.24E-06 | 3.43E-06 | 1.92E-05 | 0 | 3.18E-06 | 9.57E-06 | 7.35E-06 | 8.10E-06 | 1.45E-05 | 2.34E-06 | 1.54E-05 | k__Bacteria | p__Actinobact | c__Actinobact | o__Actinomycetales | f__Micromonospora | g__Verrucosispora | 9.14E-05 |
| 87500 | 1.50E-05 | 9.35E-06 | 2.91E-05 | 7.69E-06 | 0 | 6.36E-06 | 3.19E-06 | 2.45E-06 | 2.02E-06 | 0 | 1.17E-05 | 1.71E-06 | k__Bacteria | p__Proteobac | c__Betaprotec | o__Burkholderiales | f__Burkholderiaceae | g__Burkholderia | 8.86E-05 |
| 439331 | 4.27E-06 | 0 | 1.71E-06 | 1.92E-06 | 0 | 6.36E-06 | 3.19E-05 | 4.90E-06 | 4.05E-06 | 5.78E-06 | 2.34E-06 | 1.88E-05 | k__Bacteria | p__Actinobact | c__Actinobact | o__Actinomycetales | f__Pseudonocardia | g__Thermocrispum | 8.21E-05 |
| 92549 | 2.14E-06 | 1.25E-05 | 1.71E-06 | 1.15E-05 | 0 | 9.54E-06 | 9.57E-06 | 7.35E-06 | 1.42E-05 | 0 | 0 | 5.14E-06 | k__Bacteria | p__Proteobac | c__Betaprotec | o__Rhodocyclales | f__Rhodocyclaceae | g__KD1-23 | 7.36E-05 |
| 109717 | 0 | 9.35E-06 | 3.43E-06 | 3.84E-06 | 5.54E-06 | 3.18E-06 | 1.28E-05 | 1.23E-05 | 8.10E-06 | 2.89E-06 | 4.69E-06 | 5.14E-06 | k__Bacteria | p__Proteobac | c__Gammapro | o__Thiotrichales | f__Piscirickettsiacea | g__ | 7.12E-05 |
| 90931 | 1.07E-05 | 6.24E-06 | 1.71E-06 | 9.61E-06 | 5.54E-06 | 3.18E-06 | 0 | 2.45E-06 | 2.02E-06 | 2.02E-05 | 2.34E-06 | 1.71E-06 | k__Bacteria | p__Proteobac | c__Betaprotec | o__Burkholderiales | f__Comamonadace | g__Leptothrix | 6.57E-05 |
| 95007 | 2.14E-06 | 1.25E-05 | 3.43E-06 | 1.92E-06 | 0 | 0 | 6.38E-06 | 2.45E-06 | 2.02E-06 | 2.02E-05 | 9.38E-06 | 3.43E-06 | k__Bacteria | p__Proteobac | c__Betaprotec | o__Burkholderiales | f__Oxalobacteracea | g__Ralstonia | 6.38E-05 |
| 51325 | 1.07E-05 | 3.12E-06 | 1.71E-06 | 9.61E-06 | 0 | 3.18E-06 | 1.91E-05 | 0 | 2.02E-06 | 2.89E-06 | 4.69E-06 | 5.14E-06 | k__Bacteria | p__Firmicutes | c__Bacilli | o__Lactobacillales | f__Carnobacteriace | g__Isobaculum | 6.22E-05 |
