## Supplemental Table S4 for "Station and train surface microbiomes of Mexico City’s metro (subway/underground)"

Chloroplast

| | |
|---|---|
| S000323143 | Calycanthus floridus var. glaucus (T); AJ428413 |
| S000528956 | Pinus thunbergii; D17510 |
| S000529368 | Cuscuta reflexa; X72584 |
| S000529388 | Zea mays; X86563 |
| S000529451 | Solanum nigrum; Y18934 |
| S000531305 | Cucumis sativus (T); AJ970307 |
| S000531424 | Physcomitrella patens subsp. patens (T); AP005672 |
| S000575181 | Eucalyptus globulus subsp. globulus; AY780259 |
| S000609085 | Lactuca sativa (T); AP007232 |
| S000610747 | Phalaenopsis aphrodite subsp. formosana; AY916449 |
| S000626854 | Glycine max (T); DQ317523 |
| S000641069 | Gossypium hirsutum (T); DQ345959 |
| S000641682 | Lycopersicon esculentum; AY216521 |
| S000641743 | Ricinus communis; L37580 |
| S000641766 | Sphagnum palustre; U24592 |
| S000641768 | Nicotiana tabacum; V00165 |
| S000641769 | Doodia maxima; U24583 |
| S000641771 | Juniperus virginiana; U24586 |
| S000641773 | Pisum sativum; cr. Progress No.9; pBX5; X51598 |
| S000674963 | Helianthus annuus (T); DQ383815 |
| S000675135 | Glechoma hederacea; DQ417652 |
| S001020332 | Buxus microphylla (T); EF380351 |
| S001020365 | Coffea arabica (T); EF044213 |
| S001020373 | Cuscuta exaltata (T); EU189132 |
| S001020385 | Cuscuta reflexa (T); AM711640 |
| S001020389 | Cycas taitungensis (T); AP009339 |
| S001020393 | Daucus carota (T); DQ898156 |
| S001020397 | Dioscorea elephantipes (T); EF380353 |
| S001020405 | Drimys granadensis (T); DQ887676 |
| S001020421 | Ipomoea purpurea (T); EU118126 |
| S001020439 | Liriodendron tulipifera (T); DQ899947 |
| S001020496 | Phaseolus vulgaris (T); DQ886273 |
| S001020502 | Piper cenocladum (T); DQ887677 |
| S001020510 | Populus alba (T); AP008956 |

Mitochondria

k__Bacteria p__Proteobacteria c__Alphaproteobacteria
o__Rickettsiales f__mitochondria g__Abies s__homolepis
k__Bacteria p__Proteobacteria c__Alphaproteobacteria
o__Rickettsiales f__mitochondria g__Anomodon s__rugelii
k__Bacteria p__Proteobacteria c__Alphaproteobacteria
o__Rickettsiales f__mitochondria g__Arabidopsis s__thaliana
k__Bacteria p__Proteobacteria c__Alphaproteobacteria
o__Rickettsiales f__mitochondria g__Aristolochia s__macrophylla
k__Bacteria p__Proteobacteria c__Alphaproteobacteria
o__Rickettsiales f__mitochondria g__Asarum s__canadense
k__Bacteria p__Proteobacteria c__Alphaproteobacteria

```
o__Rickettsiales f__mitochondria  g__Azolla        s__pinnata
k__Bacteria      p__Proteobacteria        c__Alphaproteobacteria
o__Rickettsiales f__mitochondria  g__Calycanthus   s__floridus
k__Bacteria      p__Proteobacteria        c__Alphaproteobacteria
o__Rickettsiales f__mitochondria  g__Carica        s__papaya
k__Bacteria      p__Proteobacteria        c__Alphaproteobacteria
o__Rickettsiales f__mitochondria  g__Carludovica   s__palmata
k__Bacteria      p__Proteobacteria        c__Alphaproteobacteria
o__Rickettsiales f__mitochondria  g__Citrullus     s__lanatus
k__Bacteria      p__Proteobacteria        c__Alphaproteobacteria
o__Rickettsiales f__mitochondria  g__Didymeles     s__perrieri
k__Bacteria      p__Proteobacteria        c__Alphaproteobacteria
o__Rickettsiales f__mitochondria  g__Diplazium     s__pycnocarpon
k__Bacteria      p__Proteobacteria        c__Alphaproteobacteria
o__Rickettsiales f__mitochondria  g__Euptelea      s__polyandra
k__Bacteria      p__Proteobacteria        c__Alphaproteobacteria
o__Rickettsiales f__mitochondria  g__Galbulimima   s__belgraveana
k__Bacteria      p__Proteobacteria        c__Alphaproteobacteria
o__Rickettsiales f__mitochondria  g__Grevillea     s__robusta
k__Bacteria      p__Proteobacteria        c__Alphaproteobacteria
o__Rickettsiales f__mitochondria  g__Gyrocarpus    s__americanus
k__Bacteria      p__Proteobacteria        c__Alphaproteobacteria
o__Rickettsiales f__mitochondria  g__Hypseocharis  s__pimpinellifolia
k__Bacteria      p__Proteobacteria        c__Alphaproteobacteria
o__Rickettsiales f__mitochondria  g__Lupinus       s__luteus
k__Bacteria      p__Proteobacteria        c__Alphaproteobacteria
o__Rickettsiales f__mitochondria  g__Nageia        s__nagi
k__Bacteria      p__Proteobacteria        c__Alphaproteobacteria
o__Rickettsiales f__mitochondria  g__Nelumbo       s__nucifera
k__Bacteria      p__Proteobacteria        c__Alphaproteobacteria
o__Rickettsiales f__mitochondria  g__Oenothera     s__berteroana
k__Bacteria      p__Proteobacteria        c__Alphaproteobacteria
o__Rickettsiales f__mitochondria  g__Phaeoceros    s__laevis
k__Bacteria      p__Proteobacteria        c__Alphaproteobacteria
o__Rickettsiales f__mitochondria  g__Plantago      s__sericea
k__Bacteria      p__Proteobacteria        c__Alphaproteobacteria
o__Rickettsiales f__mitochondria  g__Pleurozia     s__purpurea
k__Bacteria      p__Proteobacteria        c__Alphaproteobacteria
o__Rickettsiales f__mitochondria  g__Raphanus      s__sativus
k__Bacteria      p__Proteobacteria        c__Alphaproteobacteria
o__Rickettsiales f__mitochondria  g__     s__
k__Bacteria      p__Proteobacteria        c__Alphaproteobacteria
o__Rickettsiales f__mitochondria  g__Sarcandra     s__grandifolia
k__Bacteria      p__Proteobacteria        c__Alphaproteobacteria
o__Rickettsiales f__mitochondria  g__Syntrichia    s__ruralis
k__Bacteria      p__Proteobacteria        c__Alphaproteobacteria
o__Rickettsiales f__mitochondria  g__Zea   s__luxurians
```
